# pH as an eco-evolutionary driver of priority effects

**DOI:** 10.1101/2022.04.19.487947

**Authors:** Callie R. Chappell, Manpreet K. Dhami, Mark C. Bitter, Lucas Czech, Sur Herrera Paredes, Katherine Eritano, Lexi-Ann Golden, Veronica Hsu, Clara Kieschnick, Nicole Rush, Tadashi Fukami

**Affiliations:** Department of Biology, Stanford University, Stanford, CA 94305, USA; Biocontrol and Molecular Ecology, Manaaki Whenua - Landcare Research, Lincoln 7608, New Zealand; Department of Plant Biology, Carnegie Institution for Science, Stanford, CA 94305, USA; Department of Ecology, Evolution and Marine Biology, University of California, Santa Barbara, Goleta, CA 93117, USA; Koch Institute for Integrative Cancer Research, Massachusetts Institute of Technology, Cambridge, MA 02139 USA

## Abstract

Priority effects, where arrival order and initial relative abundance modulate local species interactions, can exert taxonomic, functional, and evolutionary influences on ecological communities by driving them to alternative states. It remains unclear if these wide-ranging consequences of priority effects can be explained systematically by a common underlying factor. Here, we identify such a factor in an empirical system. In a series of field and laboratory studies, we focus on how pH affects nectar-colonizing microbes and their interactions with plants and pollinators. In a field survey, we found that nectar microbial communities in a hummingbird-pollinated shrub, *Diplacus aurantiacus*, exhibited patterns indicative of alternative stable states through domination by either bacteria or yeasts within individual flowers. In laboratory experiments, *Acinetobacter nectaris*, the bacterium most commonly found in *D. aurantiacus* nectar, exerted a strongly negative priority effect against *Metschnikowia reukaufii*, the most common nectar-specialist yeast, by reducing nectar pH. This priority effect likely explains the mutually exclusive pattern of dominance found in the field survey. Furthermore, experimental evolution simulating hummingbird-assisted dispersal between flowers revealed that *M. reukaufii* could evolve rapidly to improve resistance against the priority effect if constantly exposed to *A. nectaris*-induced pH reduction. Finally, in a field experiment, we found that low nectar pH could reduce nectar consumption by hummingbirds, suggesting functional consequences of the pH-driven priority effect for plant reproduction. Taken together, these results show that it is possible to identify an overarching factor that governs the eco-evolutionary dynamics of priority effects across multiple levels of biological organization.

## Introduction

Many ecological communities take one of multiple alternative states even under the same species pool and the same initial environmental conditions (Beisner *et al*., 2003; Chase, 2003; Scheffer *et al*., 2001; Schröder *et al*., 2005). Alternative states can vary not just in species composition, but also in functional, ecosystem-level characteristics such as invasion resistance, total biomass, and nutrient cycling (Bittleston *et al*., 2020; Delory *et al*., 2019; Leopold *et al*., 2017; Pausas & Bond, 2020; Sprockett *et al*., 2018; Suding et al., 2004). However, it is often hard to predict which of the alternative states an ecological community will take. Alternative states can be caused by priority effects, an elusive, historically contingent process in which the order and timing of species arrival dictate the trajectory of community assembly (Drake, 1991; Fukami, 2015; Palmgren, 1926; Song *et al*., 2021). To further complicate matters, the strength of priority effects is not always static, but can change rapidly through evolution (De Meester *et al*., 2016; Faillace *et al*., 2022; Faillace & Morin, 2016; Knope *et al*., 2012; Urban & De Meester, 2009; Wittmann & Fukami, 2018; Zee & Fukami, 2018). The broad scope of priority effects makes it challenging to fully understand which of the alternative states is realized under what conditions (Fukami, 2015). For any ecological community, this understanding requires simultaneous examination of the compositional, functional, and evolutionary consequences of priority effects.

Questions that need to be addressed to understand the causes and consequences of priority effects are indeed wide-ranging. For example, how do priority effects happen (mechanism)? When are priority effects particularly strong (condition)? How rapidly can priority effects change in strength (evolution)? And how do priority effects affect the functional, not just taxonomic, properties of the communities being assembled (functional consequences)? These questions are interrelated, but given that they concern different scales of time and biological organization, it seems reasonable to expect each question to involve a different set of species traits and environmental factors. In this paper, we present empirical evidence that, contrary to this expectation, it is possible for one common factor to underlie many aspects of priority effects, including mechanism, condition, evolution, and functional consequences. Specifically, we show that environmental pH is an overarching factor governing priority effects in a microbial system.

Simple microbial systems can help identify general basic principles that organize ecological communities (Cadotte *et al*., 2005; Drake *et al*., 1996; Jessup *et al*., 2004; Vega & Gore, 2018), and the nectar microbiome has recently emerged as a well-characterized simple system for understanding community assembly (Brysch-Herzberg, 2004; Chappell & Fukami, 2018; Lachance *et al*., 2001; Letten *et al*., 2018; Vannette, 2020). Our study system here consists of the bacteria and yeasts that colonize the floral nectar of the sticky monkeyflower, *Diplacus* (formerly *Mimulus*) *aurantiacus*, a hummingbird-pollinated shrub native to California and Oregon of the USA (Belisle *et al*., 2012). Initially sterile, floral nectar in individual flowers is colonized by these microbes via pollinator-mediated dispersal (Belisle *et al*., 2012; de Vega *et al*., 2021). Microbial communities that develop in nectar though this dispersal are often simple, dominated by a few species of specialist bacteria or yeast (Álvarez-Pérez *et al*., 2019; Golonka & Vilgalys, 2013; Herrera *et al*., 2010; Tsuji & Fukami, 2018; Warren *et al*., 2020). Our previous work has shown that hummingbird-mediated dispersal is highly stochastic because pollinators differ in the species identity of yeasts and bacteria they carry on their beaks and tongues (Morris *et al*., 2020; Toju *et al*., 2018; Vannette *et al*., 2021; Vannette & Fukami, 2017). Additionally, the outcome of local antagonistic interactions among these microbial species can be sensitive to relative initial abundances (Dhami *et al*., 2016, 2018; Grainger *et al*., 2019; Mittelbach *et al*., 2016; Peay *et al*., 2012; Tucker & Fukami, 2014; Vannette & Fukami, 2014). Together, the stochastic dispersal and the history-dependent interactions jointly cause priority effects in nectar microbial communities. Moreover, recent studies indicate that whether flowers are dominated by bacteria or yeasts can affect pollination and seed set, although a mechanism for this effect remains unclear (de Vega *et al*., 2022; Good *et al*., 2014; Herrera *et al*., 2013; Jacquemyn *et al*., 2021; Junker *et al*., 2014; Rering *et al*., 2020; Schaeffer & Irwin, 2014; Vannette *et al*., 2013; Vannette & Fukami, 2016, 2018; Yang *et al*., 2019). Despite the increasing amount of knowledge about this system, whether or not there is a common factor that explains the various phenomena associated with priority effects has not been investigated, to our knowledge.

In this paper, we present multiple independent pieces of evidence that all point to the pervasive role of pH across several aspects of priority effects in this nectar microbiome. First, we report the results of a field survey of *D. aurantiacus* flowers across a 200-km coastline, showing that microbial communities in the nectar exhibit distribution patterns that are consistent with the existence of two alternative community states, one dominated by nectar bacteria and the other dominated by yeasts (***Figure 1A, 2***). Next, we describe findings from laboratory experiments that suggest that inhibitory priority effects between bacteria and yeast are responsible for the alternative states we observed in the field survey and that these priority effects are in large part driven by bacteria-induced reduction in nectar pH (***Figure 1B, 3***). Using experimental evolution of nectar yeast in artificial nectar (***Figure 1C***), we then show that the low pH of nectar can cause rapid evolution of yeast populations, resulting in increased resistance to the pH-driven priority effects (***Figure 1D, 5***). Using whole genome resequencing, we then explore genomic differences between evolved yeast strains that are potentially associated with differences in the level of resistance to the priority effects (***Figure 1E, 6***). Finally, we report a field experiment showing that the low pH of nectar reduces nectar consumption by flower-visiting hummingbirds (***Figure 1F, 7***), suggesting that pH reduction is the likely explanation for our earlier observations that nectar bacteria reduced pollination and seed set.

**Figure 1:**
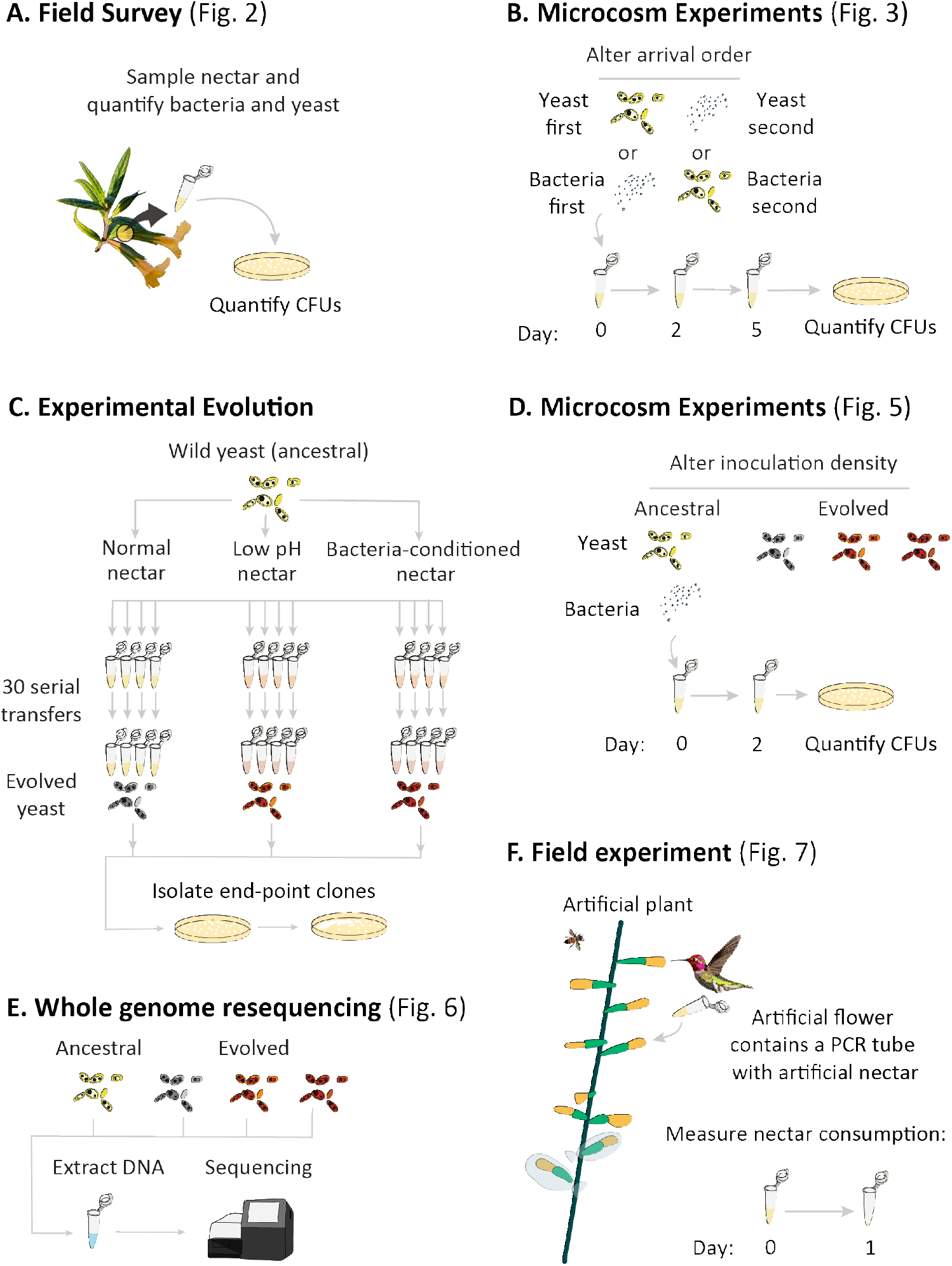
Schematic of the approaches taken in this study. Schematic of approaches undertaken in this study: (**A**) field survey to characterize the distribution of yeast and bacteria in flowers, (**B**) initial microcosm experiments to assess strength of priority effects and identify a potential driver (pH), (**C**) experimental evolution to study adaptation to low-pH and bacteria-conditioned nectar, (**D**) secondary microcosm experiments to study the effect of adaptation to nectar environments, (**E**) whole genome resequencing to identify genomic differences between evolved strains, and (**F**) a field experiment to study the effect low pH on nectar consumption by pollinators.

## Results and Discussion

### Field observations suggest bacterial vs. yeast dominance as alternative states

To study the distribution of nectar-colonizing bacteria and yeast among *D. aurantiacus* flowers, we conducted a regional survey in and around the San Francisco Peninsula of California (***Figure 1A***). Floral nectar was sampled from a total of 1,152 flowers (96 flowers at each of 12 sites) along an approximately 200-km coastline (Dhami *et al.*, 2018; Tsuji *et al.*, 2016) (***Figure 2B, C***). Bacteria and yeast were cultured on differential media to characterize floral nectar communities, which had been corroborated by molecular sequencing (Vannette & Fukami, 2017). Across all 12 sites, we found that *D. aurantiacus* flowers were frequently dominated by either bacteria or yeast, but rarely by both (***Figure 2A***). We used the CLAM classification methods program (Chazdon *et al.*, 2011) to classify flowers into four groups: bacteria-dominated flowers, yeast-dominated flowers, co-dominated flowers, and flowers with too few microbes to be accurately classified. This classification confirmed a scarcity of co-dominated flowers across all sites. This scarcity was probably caused in part by dispersal limitation (Belisle *et al*., 2012), but an additional potential reason is that bacteria and yeasts suppressed each other within flowers. Mutual suppression was suggested previously in a laboratory experiment (Tucker & Fukami, 2014). For example, if a flower first becomes dominated by bacteria, it may prevent yeast from becoming abundant, and vice versa. This mutual suppression would represent inhibitory priority effects. To test for this possibility, for each site, we calculated the proportion of co-dominated flowers that would be expected if bacteria and yeasts were distributed among flowers independently of each other. We found that the observed proportions of co-dominated flowers were lower than expected, suggesting that the observed abundance pattern was consistent with inhibitory priority effects. (**Figure 2D**; P< 0.05).

**Figure 2:**
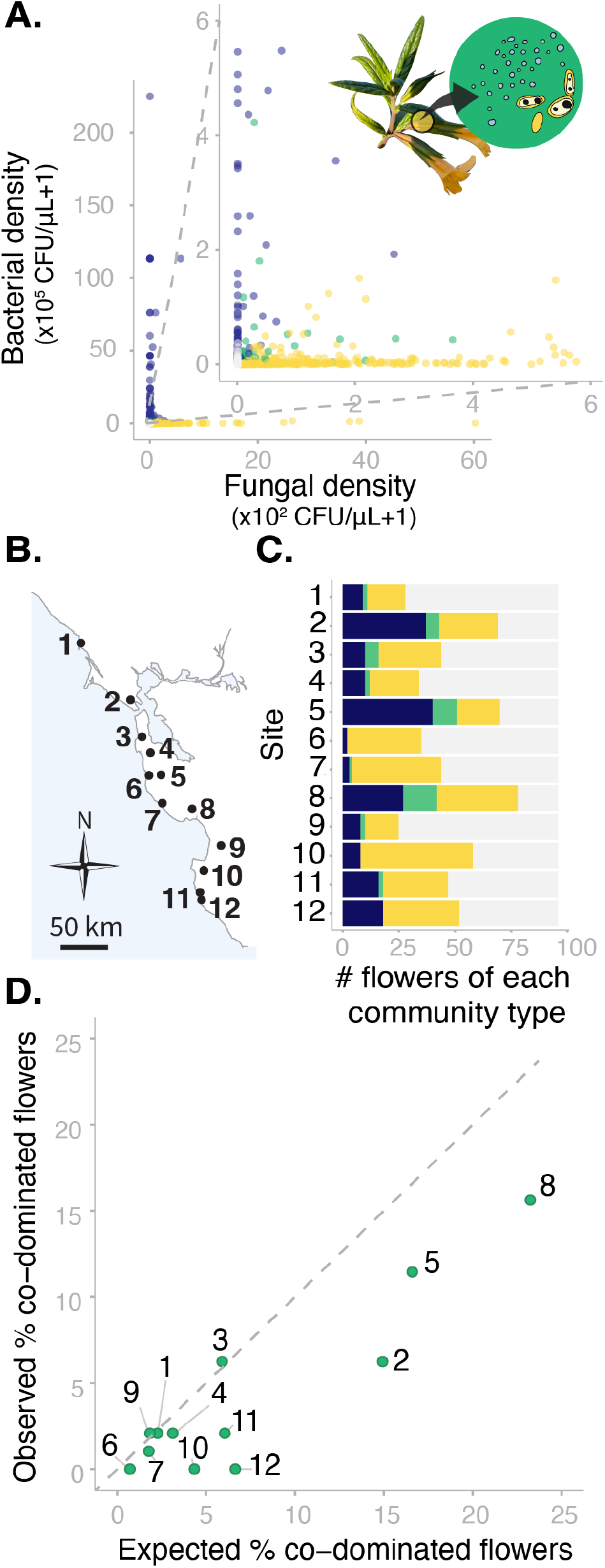
Sites vary in regional dominance of bacteria and yeast. (**A**) Classification of flowers into bacteria-dominated (blue), yeast-dominated (yellow), co-dominated (green), or microbes too rare to determine (grey). Each point represents a floral community. (**B**) *Diplacus aurantiacus* flowers were harvested from 12 field sites in and around the San Francisco Peninsula (California, USA) with (**C**) variable numbers of bacteria-dominated, fungi-dominated, co-dominated flowers, or flowers where microbes were too rare to determine. (**D**) Co-dominated flowers were observed less frequently than expected. Each point represents a site.

### Evidence for pH-mediated priority effects causing alternative states

One way to more directly test for priority effects between bacteria and yeasts is to experimentally manipulate initial relative abundance of the two microbial groups. This manipulation would allow one to determine if communities diverge such that bacteria-dominated communities develop if bacteria were initially more abundant than yeast and vice versa. Our prior work that took this approach yielded evidence for strong priority effects in *D. aurantiacus* nectar (Dhami *et al*., 2016; Peay *et al*., 2012; Toju *et al.*, 2018; Vannette & Fukami, 2014, 2017). However, these studies either looked at priority effects among yeast species or, in the case of priority effects between yeast and bacteria, focused on the yeast *M. reukaufii* and the bacterium *Neokomagataea* (formerly *Gluconobacter*) sp. (Tucker & Fukami, 2014). In a field study (Vannette & Fukami, 2017), this *Neokomagataea* sp. was the second most common bacterial taxon in *D. aurantiacus* nectar at Jasper Ridge, one of the sites used in the field survey above (site 4). The most common bacterial species, *Acinetobacter nectaris*, may also engage in strong priority effects against *M. reukaufii*, but this possibility had not been tested.

To test for priority effects between *M. reukaufii* and *A. nectaris*, we used sterile PCR tubes as artificial flowers. Each tube contained artificial nectar that closely mimicked sugar and amino acid concentrations in field-collected *D. aurantiacus* nectar. We altered the arrival order of *M. reukaufii* and *A. nectaris* and measured growth after five days, which approximated the lifespan of individual *D. aurantiacus* flowers (Peay *et al.*, 2012) (***supplementary table 1***). We found that *A. nectaris* exerted a strong inhibitory priority effect against *M. reukaufii* (***Figure 3A—supplementary table 3***) and vice versa (***figure supplement 1***).

**Figure 3:**
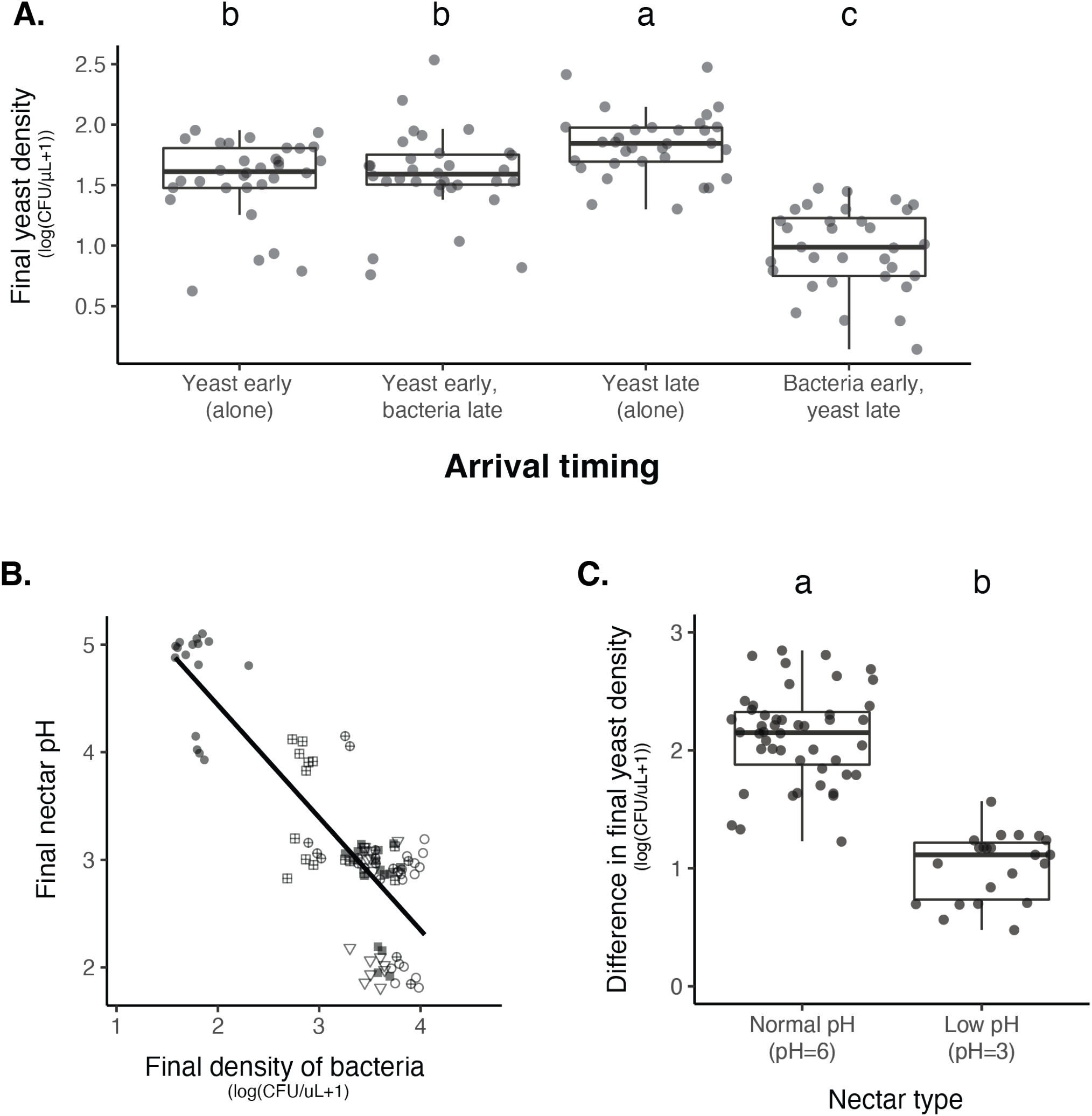
Nectar bacteria exert negative priority effects against nectar yeast, potentially due to a reduction in nectar pH. (**A**) *Metschnikowia reukaufii* (strain MR1) yeast population density after five days of growth with alternating arrival order with *Acinetobacter nectaris* bacteria or growth alone with inoculation on either the first or third day (arriving early or late) of the experiment. (**B**) Final pH of nectar after 5 days of bacterial growth; higher densities of bacteria are associated with lower pH. The shape of each point represents the various treatments represented in panel A, where early arriving yeast and late arriving bacteria are filled circles, early arriving bacteria and late arriving yeast are filled squares, early arriving bacteria and yeast together are cross-hatched squares, late arriving bacteria and yeast together are cross-hatched circles, late arriving yeast alone are open squares, early arriving yeast alone are open diamonds, late arriving bacteria alone are open circles, and early arriving bacteria alone are triangles. Points are jittered on the y-axis. (**C**) Low pH (pH=3) nectar depresses yeast growth when grown in low-density monoculture.

From our previous work with *Neokomagataea* sp. (Tucker & Fukami, 2014), we hypothesized that negative priority effects against yeast were caused by bacteria-induced reduction in nectar pH. Consistent with this hypothesis, we found in our experiments that high densities of bacteria were tightly associated with lower nectar pH (r(192)=-0.69, p<0.05, ***Figure 3B***), whereas high densities of yeast were weakly associated with higher nectar pH (***figure supplement 2***). In addition, monoculture yeast grew poorly in low pH nectar (pH = 3) when inoculated at low density (***Figure 3C***), but not if introduced at high density (***figure supplement 3***). Bacterial growth was not affected by lowered nectar pH (***figure supplement 4***). Together, these data provide further evidence that nectar pH reduction by bacteria is the mechanism of their priority effects on yeast. One likely reason for this density-dependent effect of pH on yeast is that, when yeast arrive first to nectar, they deplete nutrients such as amino acids and limit subsequent bacterial growth, thereby avoiding pH-driven suppression that would happen if bacteria were initially more abundant (Tucker & Fukami, 2014; Vannette & Fukami, 2018). Another possible reason is that larger yeast populations have higher phenotypic heterogeneity, which increases the probability that the population will include cells that are resilient to low pH. This mechanism has been suggested in the baker’s yeast, *Saccharomyces cerevisiae* (Guo & Olsson, 2016), although whether it applies to *M. reukaufii* remains to be tested.

In addition to priority effects, phenotypic plasticity or stochastic phenotypic switching could contribute to increased variation in the extent of bacterial dominance among flowers (Levy, 2016; Morawska *et al.*, 2021). In fact, previous work on *M. reukaufii* suggested that epigenetic changes contribute to its broad niche width (Herrera *et al.*, 2010). However, our experiments manipulating initial abundance provide evidence for priority effects between nectar bacteria and yeast.

### Evidence for pH as an eco-evolutionary driver of priority effects

In the field survey, we found large site-to-site variation in the relative prevalence of bacteria vs. yeast (***Figure 2C***), even though all sites followed the same L-shaped pattern that characterized alternative bacteria vs. yeast dominance (***Figure 2A***). For example, San Gregorio (site 5) had many bacteria-dominated flowers, whereas a nearby site, La Honda (site 6), had more yeast-dominated flowers. We also found through further laboratory experimentation that three genetically divergent strains of *M. reukaufii*, all of which were collected from the same site, Jasper Ridge, were differentially affected by early arrival of *A. nectaris* (***Figure 4—supplementary table 4***). Specifically, one of the three yeast strains, MR1, was more negatively affected by priority effects than the other two strains, MY0182 and MY0202, which were representative of distinct genotypic groups found across the 12 sites (Dhami *et al*. 2018). Given this genotypic and phenotypic variation among strains, it seemed conceivable that the site-to-site differences in the relative prevalence of yeasts and bacteria observed in the field could drive an eco-evolutionary feedback (sensu Post & Palkovacs, 2009), such that sites dominated by bacteria would be characterized by stronger selective pressure for yeasts to become more resistant to priority effects than sites dominated by yeasts (***Figure 2C***).

**Figure 4:**
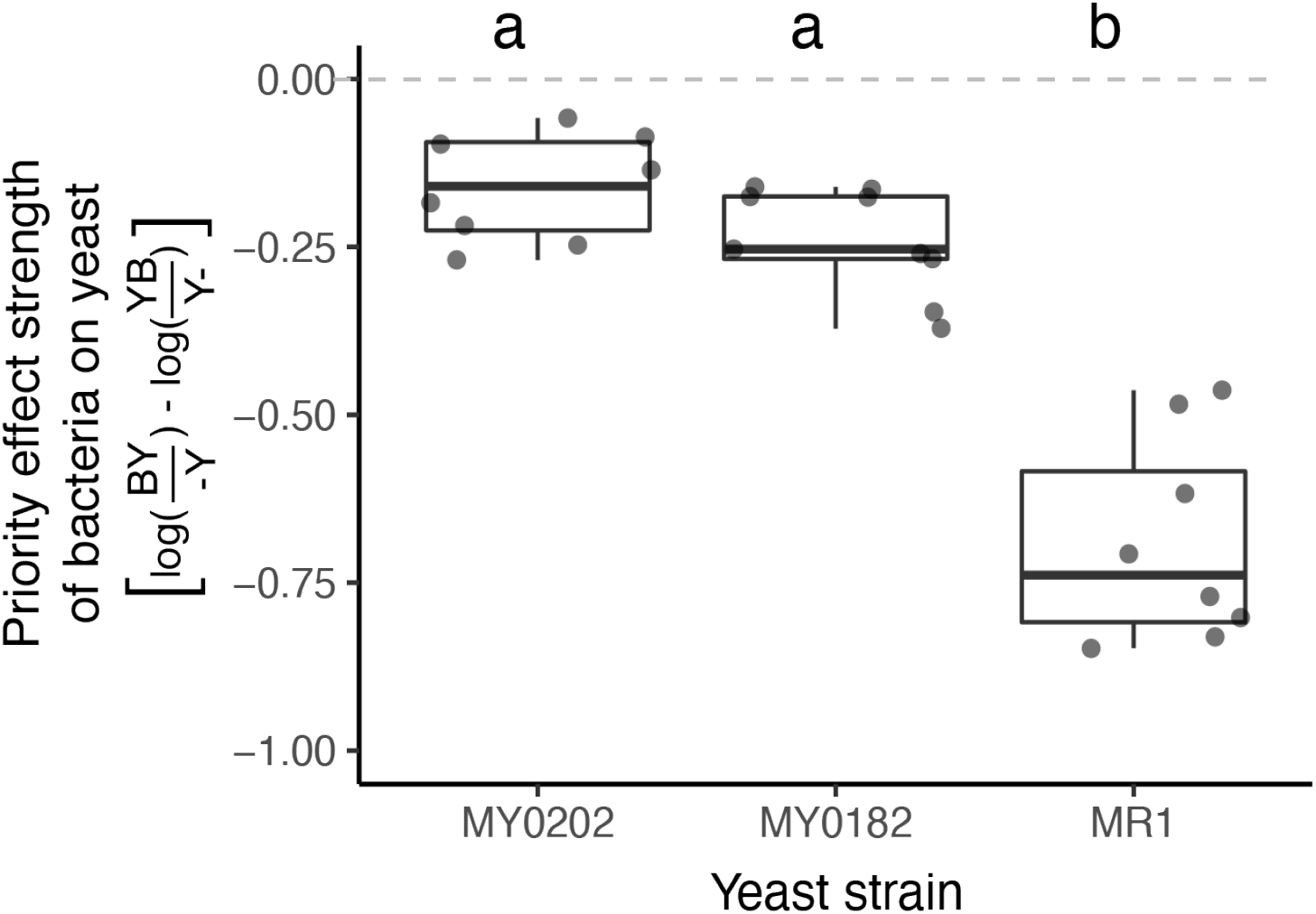
*M. reukaufii* strains differ in susceptibility to bacterial priority effects. Three strains of *M. reukaufii* are differentially affected by early arrival of bacteria. For each strain, we calculated the strength of priority effects using a metric that compares growth between varying initial inoculation densities with a competitor and alone. BY and YB represents initial dominance by bacteria or yeast (e.g. BY represents bacteria arriving early and yeast arriving late). Y- and -Y represent the comparable growth of yeast at either density (early or late) alone.

To test this hypothesis, we experimentally evolved nectar yeast under conditions that resembled repeated exposure to bacterial priority effects (***Figure 1C***). In our experimental evolution, we simulated the serial transfer of microbes between flowers via hummingbirds over the course of an entire flowering season. To this end, over 60 days, we serially transferred a single clone of *M. reukaufii* strain MR1, which we refer to as ancestral, to fresh nectar three times a week (***supplementary text 1***). This strain was isolated from Jasper Ridge. Sixty days roughly corresponds to the duration of a typical flowering season of *D. aurantiacus* at that site (Belisle *et al.*, 2012) and some of the other sites we used for the field survey (***Figure 2B***). We chose strain MR1 for this experiment because it was the most susceptible to priority effects (***Figure 4***).

We serially transferred four independent replicates of strain MR1 in three nectar types: (1) synthetic nectar where pH was 6, simulating fresh *D. aurantiacus* nectar (Vannette *et al.*, 2013); (2) synthetic nectar where pH was 3 (called “low-pH”), mimicking the pH reduction caused by growth of *A. nectaris*, our hypothesized mechanism of priority effects (***figure supplement 2***); and (3) synthetic nectar where pH was initially 6, but then conditioned by growth of *A. nectaris*. The growth of *A. nectaris* reduced pH to approximately 3 (called “bacteria-conditioned”) (***Figure 1C***). To prepare the bacteria-conditioned nectar, we grew a strain of *A. nectaris* for five days in fresh synthetic nectar and then filtered the bacteria out with a 0.2-μm filter. We selected these two experimental treatments to test whether pH reduction alone or, additionally, the production or consumption of chemical compounds by *A. nectaris* influenced *M. reukaufii*’s susceptibility to priority effects. Together, these nectar treatments represent the most extreme selective pressure that yeasts could be exposed to in the field, either never encountering flowers with early arriving bacteria or only encountering flowers where bacteria had already colonized.

To determine whether yeasts evolved in low-pH nectar were more resistant to priority effects by bacteria, we conducted laboratory experiments similar to those described above. We manipulated the initial density of a given *M. reukaufii* strain relative to a co-inoculated standard strain of *A. nectaris* and measured their abundances after three days of growth. Using these data, we quantified the strength of the bacterial priority effect on yeast using the same metric as before (***Figure 4***).

We found that, over the two months of serial transfers, yeasts that evolved in environments that constantly simulated early arrival of bacteria (“bacteria-conditioned” and “low-pH” nectars) were less affected by priority effects than the ancestral yeast or the yeast evolved in the normal nectar (***Figure 5A—figure supplement 5—supplementary table 5***). This difference in the strength of priority effects reflects the fact that the absolute growth of yeast evolved in bacteria-conditioned and low-pH nectars was less negatively affected by bacteria (***Figure 5B—supplementary table 6***). However, their improved growth was apparent only when bacteria were present (***figure supplement 6—supplementary table 7***). Together, these findings suggest that yeast can evolve to resist low pH, the very mechanism by which bacteria exert priority effects against yeast over a duration that roughly corresponds to a single flowering season of *D. aurantiacus*.

**Figure 5:**
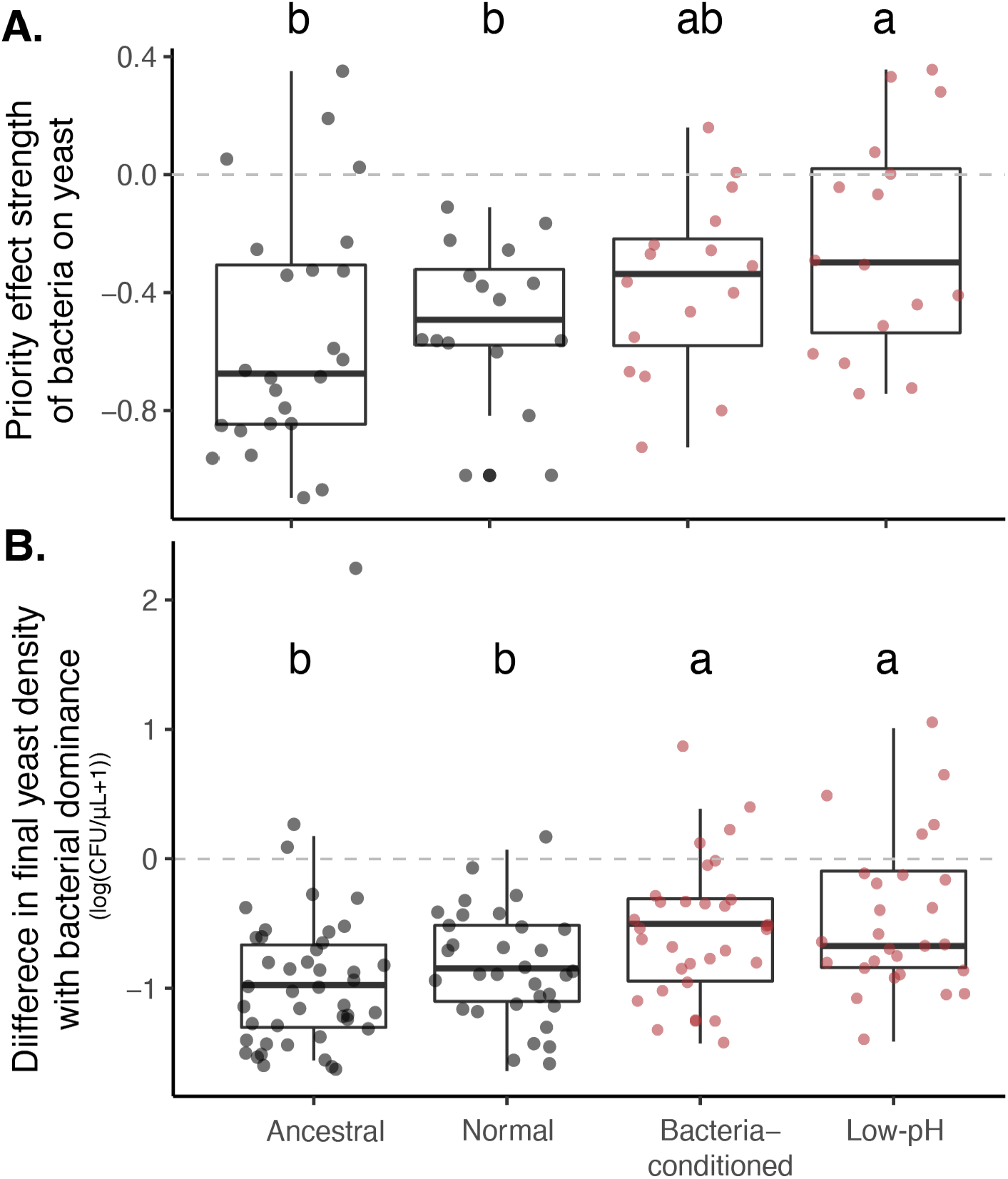
Yeast evolved with bacterial niche modification were more resistant to bacterial priority effects. (**A**) Yeast evolved in low-pH nectar was less affected by bacterial priority effects than other treatments, especially compared to ancestral yeast and yeast evolved in normal nectar. Consequently, (**B**) yeast evolved in bacteria-like nectar (both bacteria-conditioned and low-pH) was less negatively affected by initial bacterial dominance, relative to their growth alone, than ancestral yeast or yeast evolved in normal nectar.

These results point to the possibility that the variation across sites in the prevalence of bacteria-dominated flowers could create a selective pressure that drives rapid evolution of yeast to resist bacterial priority effects. However, strong local priority effects can eventually lead to regional extinction of species that engage in priority effects (Shurin *et al.*, 2004; Wittmann & Fukami, 2018). Thus, one unresolved question is how such strong priority effects can be maintained in the region. We recently used mathematical modeling to propose an “eco-evolutionary buffering” hypothesis, which provides one possible answer to this question (Wittmann & Fukami, 2018). In this model, rapid evolution maintains regional coexistence by preventing priority effects from being eliminated. While our results suggest that rapid evolution to priority effects could affect a species’ regional distribution and abundance, we did not directly explore how the dynamics within a single patch impact regional coexistence. Future work could investigate the regional effects of rapid evolution to priority effects by incorporating spatial heterogeneity and dispersal, using approaches such as experimental enrichment communities (***Estrela et al., 2021***).

### Genome sequencing to explore the genetic changes associated with pH-driven rapid evolution

To understand whether these phenotypic differences in response to priority effects have a genetic basis, we conducted full genome re-sequencing of the ancestral strain and end-point clones isolated from the four independent evolutionary replicates of each treatment (***Figure 1E—figure supplement 7—supplementary table 8***). Genomic variation across samples was characterized by calling variants against the reference genome for this species (Dhami *et al*., 2016). We explored two potential mechanisms of adaptation: loss of heterozygosity through fixation of one of the ancestral heterozygous alleles and *de novo* mutations that emerge and become fixed.

Loss of heterozygosity (LOH) is a common driver of molecular evolution in yeast (Dutta *et al.*, 2021; Peter *et al*., 2018). We explored its role in driving the observed patterns of evolution between the ancestral strain and our evolved lineages. We found that the ancestral strain exhibited heterozygosity at 25,820 sites across the 19 Mb diploid genome. Across all evolved lineages, we observed an average of 142 sites exhibiting LOH per lineage (1,699 total across all lineages). The majority of sites exhibiting LOH were shared across lineages and treatments (97%) and likely indicative of adaptation of this wild strain to the laboratory environment.

To infer genomic loci subject to treatment-specific selection pressure, we compared patterns of LOH between the lineages evolved in the normal nectar treatment and those evolved in the low-pH and bacteria-conditioned treatments. We specifically identified individual sites of LOH that were shared across multiple lineages within the low-pH and/or bacteria-conditioned treatments, but rare or absent in the normal treatment and ancestral strain (***figure supplement 8***). These sites correspond to permutation-derived p-values less than 0.1 and F_ST_ values ranging between 0.3 and 0.5, and thus are indicative of substantial genomic differentiation between treatments.

Despite the limited statistical power owing to the moderate sample size (four evolutionary replicates per treatment), we were able to identify several sites with consistent loss of heterozygosity across the bacteria-conditioned and low-pH treatments (***Figure 6A***). Several of the LOH events most predominant in the low-pH and bacteria-conditioned treatments fell near genes involved in amino acid biosynthesis (***figure supplement 3—supplementary table 9*** for all treatments). Competition for amino acids has been suggested as a mechanism by which yeast species exert priority effects against bacteria and other species of yeast in this system (Dhami *et al.*, 2016; Peay *et al.*, 2012; Tucker & Fukami, 2014). Our results suggest that adaptation to limited amino acid availability could be associated with differential resistance to priority effects in low-pH environments, though further experimentation is needed to determine the precise targets of selection.

**Figure 6:**
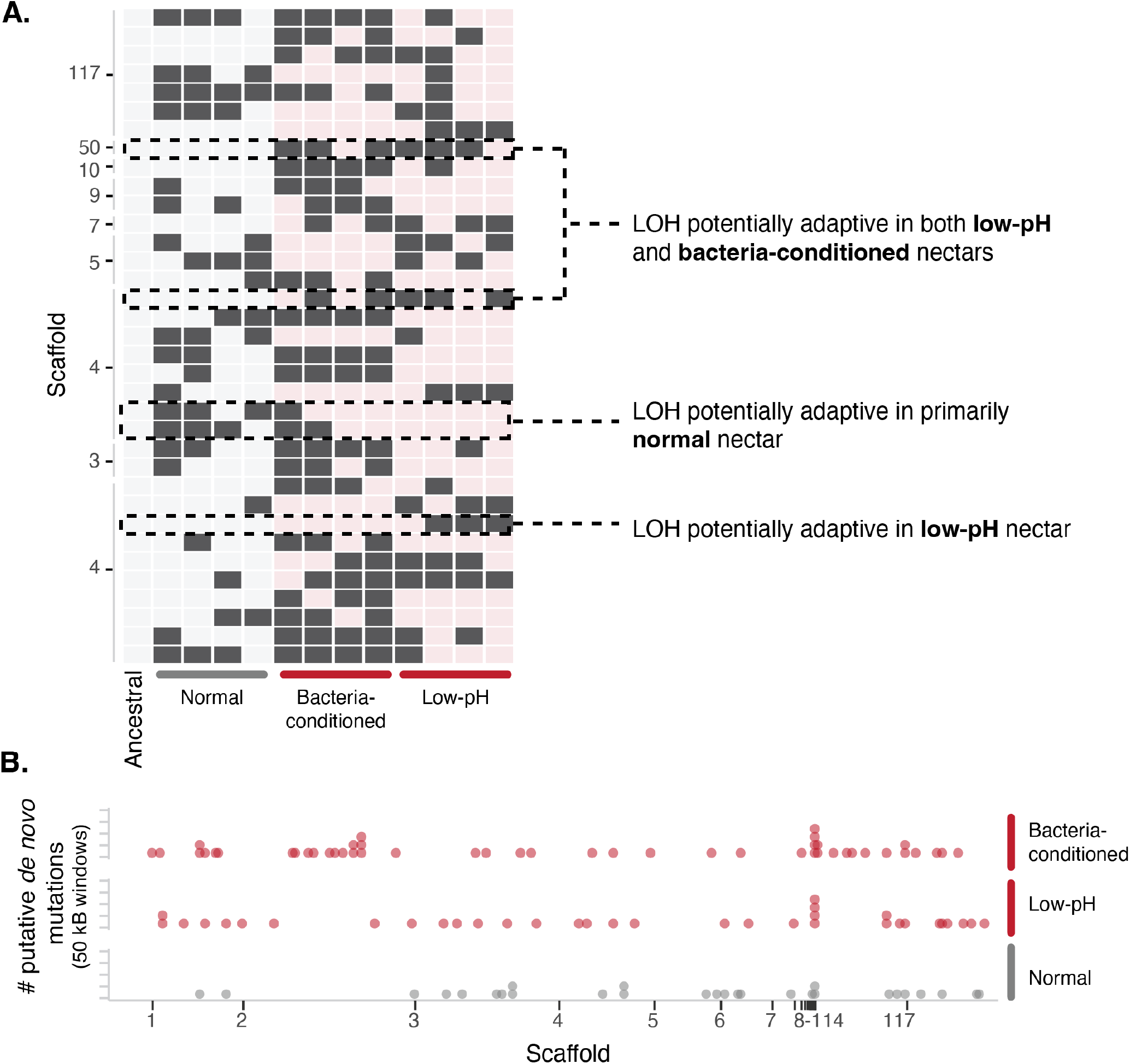
Yeast evolved to synthetic nectar. (**A**) Heat map depicting treatment-specific loss of heterozygosity (LOH). Columns represent individual samples (independent evolutionary trajectories) and rows represent single sites with LOH. White or light red boxes indicate sites without LOH, while dark grey boxes represent a site with LOH. Sites selected for the figure exhibited F_ST_ > 0.3 and permutation-derived p-value < 0.1 when comparing the ancestral strain and one of the evolved clones at that site. Boxes with dashed lines highlight examples of sites that are potentially adaptive in low-pH nectar, both low-pH and bacteria-conditioned nectar, and normal nectar but not the other two nectar types. (**B**) Distribution of putative *de novo* mutations across the genome in 50 kB windows and separated by treatment. Dots represent the sum of putative *de novo* mutations in a 50 kB window, per treatment.

We identified 146 putative *de novo* mutations across all lineages, with an average of 12 putative *de novo* mutations per lineage (***Figure 6B***). The locations of these mutations displayed some spatial concordance with patterns of LOH, corroborating the putative role of these genomic regions in adaptive evolution (***figure supplement 8***). Yeast evolved in low-pH and bacteria-conditioned nectar both exhibited putative *de novo* mutations near genes involved in solute regulation, such as membrane transport, vesicle cargo recognition, and solute carrier proteins (***supplementary table 10***). Solute regulation has been implicated in adaptation to high-osmolarity environments (Pozo *et al*., 2015), where osmotolerant yeast has been found to compete more strongly in sugar-rich nectar. Although we cannot yet directly establish that these are causal variants, they are consistent with our hypothesis that yeasts undergo rapid genetic adaptation to pH-mediated priority effects in nectar.

In summary, these results indicate that observed patterns of phenotypic evolution were concurrent to changes in genomic variation. Future work within this system could elucidate the specific loci underpinning yeast adaptation to bacterial priority effects.

### Reduced consumption of low-pH nectar by flower-visiting animals

Finally, we present evidence that bacteria-induced reduction in nectar pH not only drives microbial priority effects, but also has functional consequences for *D. aurantiacus* plants. Prior work has shown that bacterial growth in nectar can negatively affect nectar consumption by pollinators, the probability of pollination, and the number of seeds produced by *D. aurantiacus* flowers (Good *et al.*, 2014; Vannette *et al.*, 2013; Vannette & Fukami, 2016, 2018). However, the mechanism behind this effect remains unknown. To test whether pH affects nectar consumption by flower-visiting animals, we conducted a field experiment using an array of artificial plants at an outdoor garden at the Stock Farm Growth Facility on the Stanford University campus, located 5 km from Jasper Ridge. We used artificial flowers attached to garden stakes (***Figure 7A***) and measured changes in nectar volume after one day, correcting for reduction in volume through evaporation as opposed to consumption by animals like hummingbirds and bees. Extensive video footage that we recorded of the artificial flowers indicated that the consumption was primarily by hummingbirds.

**Figure 7:**
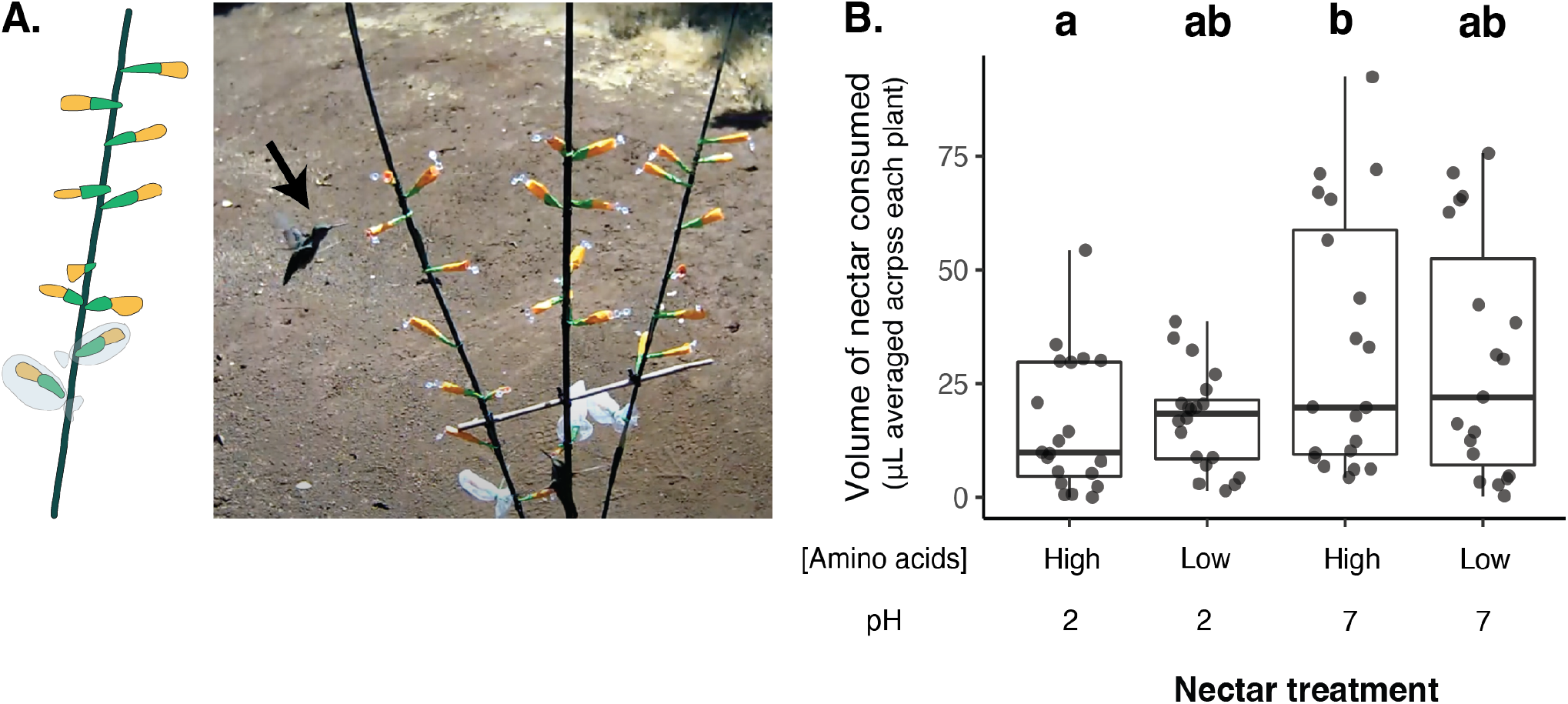
:Low nectar pH reduces nectar consumption by flower-visiting animals. (**A**) Photo of field experiment setup: hummingbird visits artificial flowers containing PCR tubes of nectar attached to a garden stake. See this [link](https://www.youtube.com/watch?v=LbD2r43dvnQ) for video. (**B**) Flower-visiting animals consumed less nectar from flowers containing low pH, high amino-acid nectar than high pH, high amino acid nectar.

We used synthetic nectar with either pH = 7, representing the highest potential nectar pH, or pH = 2, representing the lowest potential nectar pH, to evaluate the effect of pH reduction on nectar consumption. In addition, because nectar-colonizing microbes, particularly *M. reukaufii*, can reduce amino acid concentrations in nectar (Dhami *et al.*, 2016; Vannette & Fukami, 2018), we also manipulated the concentration of amino acids in the artificial nectar to quantify the relative importance of pH compared to amino acid concentration. Nectar contained either a high (0.32 mM) or low (0.032 mM) concentration of amino acids in a factorial design with pH manipulation. We found that for the high amino acid nectar, which resembled the synthetic nectar used in the previously described microcosm experiments, less nectar was consumed from flowers with low-pH nectar (***Figure 7B***). This result indicates that low pH, but not low concentrations of amino acids, caused a reduction in nectar consumption. Because the bacteria lower pH (***Figure 3B***), these findings suggest that bacteria-dominated flowers can negatively affect nectar consumption by pollinators by lowering nectar pH.

Overall, these results suggest that in addition to impacting microbial interactions and community assembly, low nectar pH associated with bacteria-dominated flowers affects the relationship between plants and pollinators via microbial priority effects. pH is thought to be a strong predictor of community assembly in other microbial systems as well (Ratzke & Gore, 2018), including those in the soil (Fierer & Jackson, 2006; Tedersoo *et al.*, 2014) and the human gut (Beasley *et al.*, 2015). Priority effects have been observed in some of these microbial habitats, suggesting that pH may govern priority effects as an overarching factor across many types of microbial communities.

## Conclusion

The independent series of evidence presented here collectively suggests that it is possible to find a single deterministic factor, such as pH, that explains a range of the ecological and evolutionary consequences of priority effects that drive alternative community states (***Figure 8***). Using nectar-inhabiting microbes, this study has suggested that pH-mediated priority effects (***Figures 3 and 4***) give rise to bacterial and yeast dominance as alternative community states (***Figure 2***). However, pH is not merely a mechanism of the priority effects, but it also determines the strength of the priority effects, fuels rapid evolution that changes the strength of the priority effects (***Figures 5 and 6***), and causes functional consequences of the priority effects for the host plants’ reproduction (***Figure 7***). These findings show that historical contingency via priority effects can be understood more simply than generally thought once we identify a common factor that dictates the various aspects of priority effects. We suggest that approaches like ours can promote a fuller understanding of community assembly than otherwise possible.

**Figure 8:**
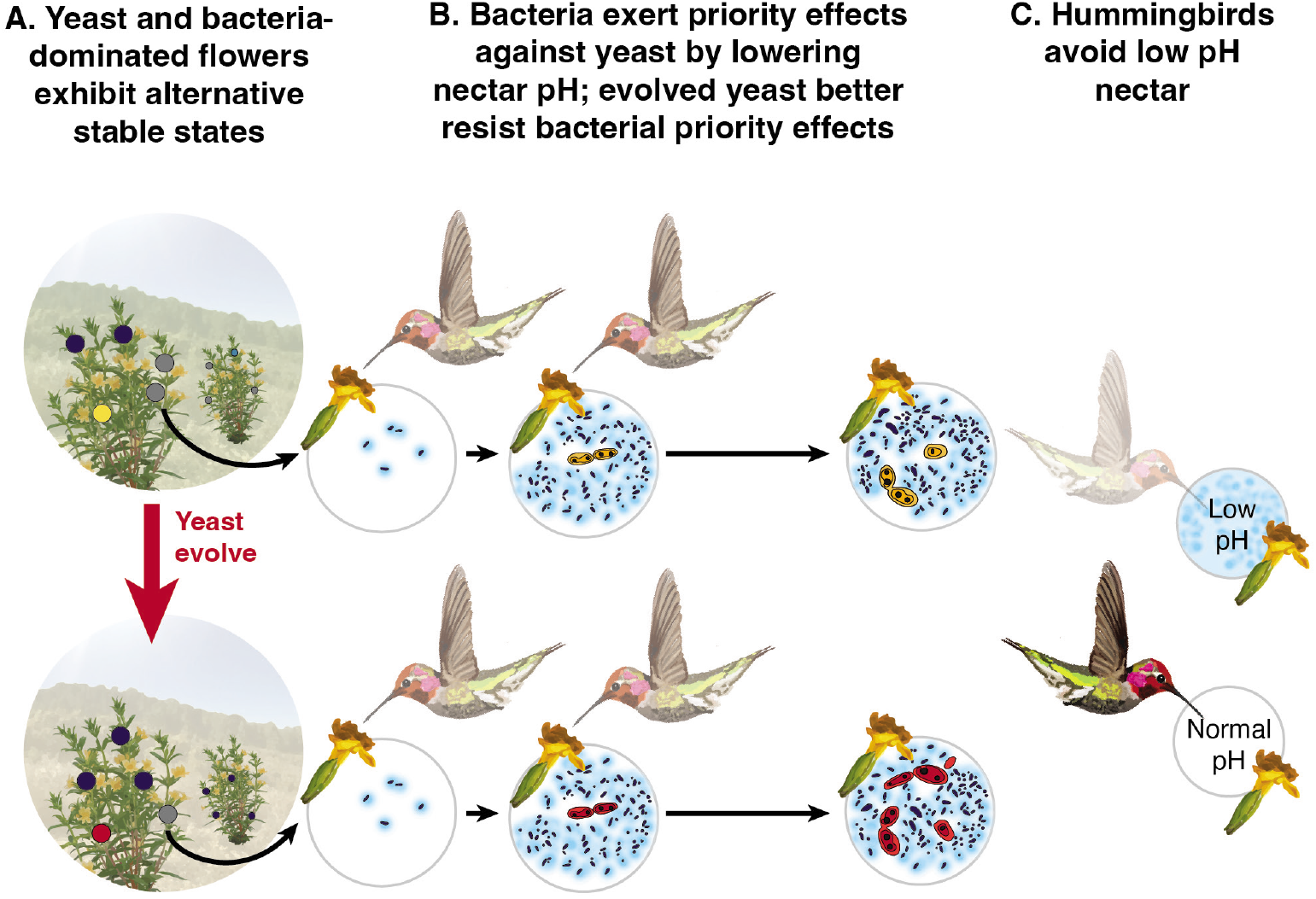
pH is an eco-evolutionary driver of priority effects in nectar microbes. (**A**) Field observations show that nectar yeast and bacteria in individual flowers exhibit alternative stable states, where some flowers are either dominated by bacteria (dark blue), yeast (yellow (ancestral) or red (evolved)) or lack significant microbial growth (gray). (**B**) Laboratory experiments identify negative priority effects between bacteria (dark blue) and yeast (yellow (ancestral) or red (evolved)) that lead to alternative states; for example, where early arriving bacteria lower the pH of nectar (depicted here in light blue), limiting the growth of later arriving yeast. Experimentally evolved nectar yeast (red) was less affected by bacterial priority effects, supporting pH as the key mechanism by which nectar bacteria inhibit yeast growth. (**C**) A field experiment shows one functional outcome of bacterial dominance in nectar: low nectar pH decreases nectar consumption by pollinators. Graphics modified from Chappell & Fukami, 2018.

## Materials and methods

### Field survey

Field sampling was conducted at 12 sites throughout the San Francisco Peninsula (***Figure 2***). At each site, eight *D. aurantiacus* flowers with a closed stigma were harvested from 12 plants. Flowers with a closed stigma are more likely to have been pollinated than flowers with an open stigma (Fetscher & Kohn, 1999). From each flower, nectar was extracted using a clean 10-μL microcapillary tube, measured, and diluted in 40 μL sterile PCR-grade water. Sample size was determined based on previous data (Vannette *et al.*, 2013). In the laboratory, samples were further diluted and plated on yeast malt agar (YMA; Difco, Sparks, MD, USA) supplemented with 100 mg/L of the antibacterial chloramphenicol and on Reasoner’s 2A agar (R2A; BD Diagnostics, Sparks, MD, 31 USA) supplemented with 20% sucrose and 100 mg/L of the antifungal cycloheximide. Plates were incubated at 25°C for five days and colony forming units (CFUs) were counted on the YMA and R2A plates to estimate bacteria and yeast abundance, respectively. Molecular identification of colonies via amplification and sequencing of ribosomal genes (16S rDNA for bacteria and LSU rDNA for fungi) (Good *et al.*, 2014) indicated that *Neokomagataea* sp. was capable of forming colonies on the chloramphenicol-supplemented YMA and that these bacterial colonies tended to be distinctly smaller than yeast colonies. Observations of cells from these small colonies under a compound microscope confirmed that the size of the cells tended to be distinctly smaller than cells found in yeast colonies. Furthermore, we found that the small colonies failed to proliferate during subsequent sub-culturing on YMA, whereas most colonies identified as yeast grew optimally, providing additional indication that the small colonies were bacteria. For these reasons, we removed the small colonies from CFU on YMA for our estimation of yeast abundance. In a previous study (Vannette & Fukami, 2014), Illumina MiSeq metabarcoding conducted on *D. aurantiacus* nectar samples indicated that the dominant 45 taxa of bacteria (e.g., *Acinetobacter, Neokomagataea*, and *Pseudomonas* spp.) and yeast (e.g. *Metschnikowia, Starmerella*, and *Cryptococcus* spp.) recovered from culture-independent analyses also form colonies on R2A and YMA, respectively (Aizenberg-Gershtein *et al.*, 2015). Thus, our culture-based methods for estimating bacterial and yeast abundance should provide useful, though not perfectly precise, information on their abundances.

We used the CLAM (Classification Methods) program (Chazdon *et al.*, 2011) to classify flowers into (1) bacteria-dominated flowers, (2) yeast-dominated flowers, (3) co-dominated flowers, and (4) flowers with too few microbes to be classified into any of the three other groups. The CLAM program was originally developed to classify species into habitat specialists and generalists, focusing on two types of habitat, with species grouped into four categories: habitat A specialists, habitat B specialists, generalists, and species that are too rare to be classified into any of the other three groups. However, the same principle can be applied to classifying microbial communities into bacteria-dominated, yeast-dominated, co-dominated, and microbial communities with too few bacteria and/or yeast for classification. We applied the CLAM method separately for flowers in each of the 12 sites. At each site, the resultant flowers (bacteria-dominated, yeast-dominated, co-dominated, too few to classify) were summed and a two-sided Fisher’s exact test was used to determine differences between sites.

### Synthetic nectar preparation

Synthetic nectars were prepared based on previous chemical analysis of *D. aurantiacus nectar* (Peay *et al.*, 2012). The sugars and amino acids found to be abundant in this analysis, including fructose (4%), glucose (2%), sucrose (20%), serine (0.102 mM), glycine (0.097 mM), proline (0.038 mM), glutamate (0.035 mM), aspartic acid (0.026 mM), GABA1 (0.023 mM), and alanine (0.021 mM), were mixed until dissolved, adjusted using NaOH to pH=6, filtered through a 0.2 μm filter, and stored at −20 °C until used. We made two additional synthetic nectars that were modified from the synthetic nectar recipe above to mimic chemical changes to nectar induced by early arrival of bacteria *A. nectaris*. “Bacteria-conditioned” nectar was prepared by inoculating *A. nectaris* (strain FNA3) at a density of 200 cells/μL into two liters of synthetic nectar. The nectar was conditioned for five days by incubating at 25°C while shaking at 200 RPM. This conditioned nectar was sterilized by filtering through a 0.22μm filter and stored at −20°C until use. This nectar had a pH of 3. The second type of the two additional nectars was “low-pH nectar,” which was prepared by adjusting the pH of the original synthetic nectar to pH=3 using HCl.

### Experimental evolution

The bacterium *A. nectaris* (strain FNA17, isolated in 2017) and the yeast *M. reukaufii* (strain MR1, isolated in 2010) (***supplemental table 1***) were isolated from floral nectar of *D. aurantiacus* growing at Jasper Ridge Biological Preserve and glycerol stocks were kept at −80°C. *A. nectaris* was streaked on tryptic soy agar (TSA) with 100 mg/L cycloheximide and *M. reukaufii* was streaked on yeast mold agar (YMA) with 100 mg/L chloramphenicol from glycerol stocks. Plates were incubated for two days at 25°C. Single colonies from these strains were re-suspended in synthetic nectar and inoculated into 96-well plates containing 120 μL nectar per well at a density of approximately 200 cells/μL. Four replicate isolates of each strain were inoculated into low-pH (pH=3), normal (pH=6), and bacteria-conditioned (pH=3) nectar. Plates were covered with sterile, air-permeable membranes (Breathe-Easy sealing membranes, Millipore Sigma, Darmstadt, Germany) and incubated at 25°C. 10 μL of each culture was transferred to a new 110 μL of nectar for 30 transfers (every Monday, Wednesday, and Friday). Every two weeks, the remaining culture of the evolving strains was frozen in 25% glycerol and stored at −80°C for later experimental use.

To isolate individual clones of yeast evolved in each treatment, 20 μl of culture from the final transfer was diluted with 200 μl sterile phosphate buffered saline and plated onto YMA plates with 100 mg/L chloramphenicol and incubated at 25°C for two days. From each plate, ten colonies were re-streaked and stored for future use as single colony pure cultures. One of these isolates per replicate per treatment was used for subsequent priority effects experiments.

### Priority effects experiments

For the first round of priority effects experiments, *A. nectaris* and different strains of *M. reukaufii* (strains MR1, MY0182, and MY0202) were plated on TSA plates with 100 mg/mL cycloheximide or YMA plates with 100 mg/mL chloramphenicol and incubated for 2 days at 25°C. Individual colonies from strains were resuspended in synthetic nectar and inoculated into PCR tubes at a density of 200 cells/μL in a fully factorial design (***supplementary table 2***) on either day 0 or day 2 of the experiment. Microcosms were covered with sterile, air-permeable membranes (Breathe-Easy Sealing Membrane, Sigma-Aldrich, Darmstadt, Germany) and incubated at 25°C. After five days, cultures were resuspended, serially diluted in sterile 0.85% NaCl and plated on TSA plates with 100 mg/L cycloheximide or YMA plates with 100 mg/L chloramphenicol. After two days of incubation at 25°C, colony forming units were counted to quantify microbial growth in the competition experiment. Experiments were repeated over multiple weeks (rounds) with two or three biological replicates per week.

For the second round of priority effects experiments, ancestral *A. nectaris* (strain FNA17) and ancestral and evolved *M. reukaufii* (strain MR1) were plated on TSA plates with 100 mg/L cycloheximide or YMA plates with 100 mg/L chloramphenicol and incubated for 2 days at 25°C. Single strains subcultured from individual colonies were resuspended in synthetic nectar and inoculated into 96-well plate microcosms at either a density of 10,000 cells/μL (“early”) or 10 cells/μL (“late”) with a competitor species or alone in a fully factorial design (***supplementary table 3***). After inoculation, 96-well plates were covered with sterile, air-permeable membranes and incubated at 25°C. After two days, cultures were resuspended, serially diluted in sterile 0.85% NaCl, and plated on TSA plates with 100 mg/L cycloheximide or YMA plates with 100 mg/L chloramphenicol. After two days of incubation at 25°C, colony forming units were counted to quantify microbial growth in the competition experiment. Experiments were repeated over four rounds with two independent replicates (96-well plates) per week. The data from the first week of each academic quarter was omitted from the final analysis due to variation in training new student researchers. Two additional rounds of this experiment were conducted using low-pH nectar instead of the normal nectar to investigate differential growth of ancestral *M. reukaufii* in low-vs. normal-pH nectar. For both experiments, sample size was determined based on previous data (Belisle *et al.*, 2012; Peay *et al.*, 2012; Tucker & Fukami, 2014) and feasiblity.

### Priority effects statistical analysis

For all analysis, colony counts were log_10_-transformed and analysis was conducted using the *glmmTMB* package (1.7.14) in R (3.6.0).

To calculate the effect of arrival order on bacteria and yeast growth, we used a linear mixed model predicting the final density of yeast or bacteria based on the priority effects treatment. We included the week of the experiment (round) and replicate tubes as nested random effects. Then, we used a post-hoc test using the *emmeans* package (1.7.2) to see whether for each priority effect treatment (representing different arrival orders), there was a difference in microbial growth.

To calculate the effect of bacterial density on final nectar pH, we used a Spearman’s rank order correlation model with the log-transformed final bacterial density as the predictor and final nectar pH as the response variable.

We calculated the differential response of *M. reukaufii* strains to bacterial priority effects by first calculating priority effect strength (PE) as PE = log(BY/(-Y)) - log(YB/(Y-)), where BY and YB represents early arrival (day 0 vs. day 2 of the experiment) by bacteria or yeast, respectively. -Y and Y-represent the comparable growth of yeast at either arrival time alone, and treatment densities were averaged by round of the experiment. Next, we used a linear mixed model predicting the priority effect strength of bacteria on yeast by yeast strain. We included the week of the experiment (round) and replicate tubes as nested random effects. Then, we used a post-hoc test using the *emmeans* package (1.7.2) to see whether for each yeast strain, there was a difference in the strength of priority effects by bacteria.

For experiments comparing the effect of evolutionary history on priority effect strength (PE), we calculated the priority effect strength (PE), as described above, as well as differences in growth by evolutionary history (ancestral or experimental evolutionary treatment) by subtracting yeast growth with initial bacterial dominance (BY) from monoculture growth at the same density (-Y) for each replicate. We used linear mixed models predicting either the final density of yeast or the difference in yeast growth based on priority effects treatment. We included the week of the experiment (round) and 96-well plate as nested random effects and evolutionary replicate (independent evolutionary trajectory) as a random effect. Then, we used a pairwise post-hoc test using the *emmeans* package (1.7.2) to see whether, for each evolution treatment (ancestral, normal, low-pH, bacteria-conditioned), there was a difference in microbial growth by priority effects treatment (representing different arrival orders).

We calculated the effect of nectar type on final yeast density by conducting the evolutionary priority effects experiments described above in both normal and low pH nectars. By combining these experiments, conducted over six total weeks, we used a linear mixed model with the evolution treatment (ancestral or three evolved treatments), the priority effect treatment (related to arrival order), and nectar type as predictors and the final density of yeast as a response. We included independent evolutionary replicates as a random effect and included nested random effects of the week the experiment was conducted and the 96-well plate (experiments were conducted across two 96-well plates). We used a post-hoc test to see whether, for each evolution treatment (ancestral, normal, low-pH, bacteria-conditioned), there was a difference in microbial growth in either low or high pH by priority effects treatment (yeast at a high density, yeast at a low density, etc.).

### Genome sequencing

Ancestral *A. nectaris* (strain FNA17) and ancestral and single clones of evolved *M. reukaufii* (strain MR1) from glycerol stocks were plated on TSA plates with 100 mg/L cycloheximide or YMA plates with 100 mg/L chloramphenicol, respectively and incubated for 2 days at 25°C. Single colonies were selected and subcultured in 10 mL yeast mold broth (yeast) or tryptic soy broth (bacteria). Cultures were grown for 15 hours at 24°C and 200 RPM. Overnight cultures were adjusted to 1×10^7^ yeast cells/μL or 1×10^7^ bacteria cells/μL and yeast cells were treated with Zymolase (100 U/μL) for 30 minutes at 30°C. DNA was extracted using Qiagen Blood and Tissue Kit (Qiagen, Redwood City, CA) and quantified using a Qubit HS Kit (ThermoFisher Scientific, Waltham, MA). Dual-indexed genomic libraries were prepared using a Nextera XT index kit (Illumina, San Diego, CA) and pooled for sequencing. Libraries were sequenced on a single Illumina MiSeq V3 run, which produces 300 bp, paired-end reads. This resulted in a total of 675M mapped reads with an average sequencing depth of 444X per sample.

### Read filtering, sequence alignment, and variant calling

Variants in genomic data were identified with *grenepipe* (0.1.0) (Czech & Exposito-Alonso, 2021), an automated variant calling pipeline for *snakemake* (Köster & Rahmann, 2012). Raw fastq sequencing files were trimmed using *trimmomatic* (0.36) (Bolger *et al.*, 2014). Filtered reads were mapped to the reference genome provided by Dhami *et al*. 2016 using *bwa mem* (0.7.17) (Li & Durbin, 2009). Mapped reads were sorted and indexed using *samtools* (1.9) (Li *et al.*, 2009) and duplicate reads were removed using the *picard* software (2.22.1) (Picard Toolkit, 2019). Genetic variants, both single nucleotide polymorphisms (SNPs) and insertions/deletions (indels) were identified using *freebayes* (1.3.1) (Garrison & Marth, 2012). The resulting VCF file was filtered using *vcftools* (2.22.1) (Danecek *et al.*, 2011), which yielded a total of 3,016,402 polymorphic sites across all samples. Quality of filtered reads and their adapter content was determined by *fastqc* (0.11.9) with additional statistics collected from *qualimap* (2.2.2a) (Okonechnikov *et al.*, 2015) for genome coverage, and *samtools stats* (1.6) (Li *et al.*, 2009) and *samtools flagstat* (1.10) (Li *et al.*, 2009) for mapping and alignment metrics. Output of all tools was then summarized in a *MultiQC* (1.9) report (Ewels *et al.*, 2016), which we provide as ***supplementary table 7***.

### Loss of heterozygosity and de novo mutation analyses

To identify loss of heterozygosity (LOH) between the ancestral strain and the evolved clones, we first identified those sites in which the ancestral strain was heterozygous (26,490 locations, corresponding to ~0.14 % of all sites). LOH events were then characterized as those instances in which any of the evolved strains was homozygous at a site identified as heterozygous in the ancestor, an event occurring 1,699 times across all end point clones. We did not expect to see contiguous segments of the genome where all sites show LOH because we do not see recombination in this yeast species (they are clonal), assessed four independent evolutionary replicates per treatment, and conducted the experiment over a relatively short period of time. To explore evidence of treatment-specific LOH events, we computed an odd’s ratio of alternate to reference allele counts in the control relative to the low-pH or bacteria-conditioned treatment genotypes. For each odds ratio, we generated 1,000 permuted odds ratios in which the genotype data at that site were randomly shuffled across control treatment and either low-pH or bacteria-conditioned samples. P-values were subsequently computed by comparing the observed odds ratio to that of the 1,000 permuted ratios.

Next, to further quantify the genomic differentiation between treatments, we computed a Weir-Cockerham estimator of F_ST_ using *vcftools*. While genome-wide F_ST_ was exceedingly low between the normal and low-pH treatment (0.0013), as well as the normal and bacteria-conditioned treatment (0.0014), a small subset of sites (50/26,380) exhibited extreme divergence with F_ST_ ranging between 0.3-0.5. Unsurprisingly, those sites with elevated F_ST_ largely overlapped with LOH sites with permutation-derived p-values < 0.1 (35/25,545). Putative *de novo* mutations were identified using a custom program to identify SNPs that differed between each pair of ancestral and evolved strains (245 singletons out of 3,016,402 sites). The program was written using the C++ library *genesis* (0.25.0) (Czech *et al.*, 2020). To ensure that these singletons were accurate, we generated a mappability score across the genome using *GenMap* (Pockrandt *et al.*, 2020) and filtered by singletons in sites with a mappability score of 1 (representing a unique k-mer region with high mappability), reducing the number of putative *de novo* mutations to 156 sites.

To explore the functional basis of genomic divergence between ancestral and evolved strains, we identified genes in close proximity to those sites with treatment-specific patterns of LOH (permutation-derived p-value < 0.1; F_ST_ = 0.3-0.5) and/or those containing a putative *de novo* mutation. We re-annotated the *M. reukaufii* reference genome provided by Dhami *et al*. 2016 using the InterPro database (*interproscan* 5.45-80.0) (Blum *et al*., 2021) and identified the closest gene with an InterPro annotation within 5 KB upstream/downstream of each candidate site. This annotation resulted in functional information for 42 of the candidate LOH sites and 94 of the putative *de novo* mutations. Distance to the nearest gene and its associated functional annotation information are provided in ***supplementary tables 9 and 10***.

### Nectar consumption field experiment

To study the effect of nectar chemistry on nectar consumption by flower-visiting animals, an array of artificial plants was constructed at the Plant Growth Facility on Stock Farm Road on the Stanford University campus. Artificial plants were created using 1 m tall garden stakes with 10 artificial flowers arranged vertically on each stake. Each artificial flower consisted of a pipette tip wrapped in orange and green tape and contained a 200 μL PCR tube of nectar to resemble real *D. aurantiacus* flowers in color, shape, and size. Two artificial flowers were covered with small white organza bags (ULINE, Pleasant Prairie, WI) and were used to estimate the loss of nectar volume by evaporation. See ***Figure 7A*** for a schematic of these artificial plants. Artificial plants were placed 3-5 meters apart and positioned near potted *D. aurantiacus* plants.

Across 2016-2018, 188 artificial plants were deployed and several nectar types were tested (n=610). In 2017, nectar treatments included two levels of amino acid concentration (high and low) and two pH levels (7 and 2). The high amino acid nectar contained 0.32 mM casamino acids whereas the low amino acid nectar contained 0.032 mM casamino acids. In 2016 and 2018, nectar treatments had two sugar levels (regular and altered sugars) and two pH levels (7 and 2). The control sugars had 30% sucrose, 1.5% glucose, and 4% fructose. The altered sugars had 20% sucrose, 0.5% glucose and 6% fructose. Results from these findings are reported in ***figure supplement 9***.

Undergraduate students and staff in the BIO 47 (formerly 44Y) course at Stanford University exposed 100 μL nectar in each flower to potential pollinators for 22 hours (approximately 3 PM to 1 PM) and measured the change in nectar using 100μL Drummond calibrated microcapillary tubes. Nectar removal from each tube was calculated by subtracting the volume of nectar remaining in experimental tubes from the average volume of nectar in bagged controls. Over the duration of the experiment, we observed visitation to artificial flowers by primarily Anna’s hummingbirds (*Calypte anna*) and excluded ant visitation by painting the base of each stake with Tanglefoot (Tanglefoot, Marysville, OH, USA). We calculated the difference between treatments and the volume of nectar consumed using a Mann-Whitney U test, and the difference in proportion of flowers consumed using a chi-square test.

## Supporting information

Chappell et al. - supplement

## Acknowledgements

We thank Itzel Arias Del Razo, David Cross, Po-Ju Ke, Carolyn Rice, Nic Romano, and Kaoru Tsuji for assistance with field and laboratory work on the field survey (***Figure 2***); Sergio Álvarez-Pérez for his assistance with preliminary priority effects experiments (***Figure 3***); Adrianna Garner, Jonathan Hernandez, and Paloma Vazquez for their assistance with laboratory work with the priority effects microcosm experiments (***Figure 5***); Jonathan Barros and Briana Martin-Villa for assistance with additional analysis; the students, teaching assistants, and instructors of the Biology 47 (formerly 44Y) class (***Fukami, 2013***) in 2016-2018, for assistance with the artificial flower experiments (***Figure 7***) and in 2019 for assistance with the experimental evolution experiment, in particular Nona Chiariello, Jess Coyle, Bill Gomez, Trevor Hebert, Daria Hekmat-Scafe, Shyamala Malladi, Jesse Miller, and Griselda Morales; Joo Hee Ahn and David Mosko for inspiring us to investigate nectar pH through their Biology 44Y independent project in 2011; Ivana Cvijovic, Moisés Expósito-Alonso, Grant Kinsler, Lauren O’Connell, Kabir Peay, Dmitri Petrov, Gavin Sherlock, and the members of the community ecology group at Stanford for discussion and comments.

## Data and materials availability

Raw sequencing reads are available at NCBI Sequence Read Archive (**BioProject PRJNA825574**).

## Funding

This work was supported by the National Science Foundation (**DEB 1149600, DEB 1737758**), Stanford University’s Terman Fellowship, and donation of sequencing materials from Illumina, Inc. CRC was supported by a National Science Foundation Graduate Research Fellowship (**DGE 1656518**) and a Stanford Graduate Fellowship. LC was supported by the Carnegie Institution for Science at Stanford, California, USA. MKD was supported by Marsden Fund Grant (**MFP-LCR-2002**). SHP was supported by the Life Sciences Research Foundation. Undergraduate researchers KE, LG, and CK were supported by the Stanford Department of Biology VPUE Biology Summer Research Program.

All other data and code reported in this paper are available at: https://gitlab.com/teamnectarmicrobe/n06_nectarmicrobes_ecoevo

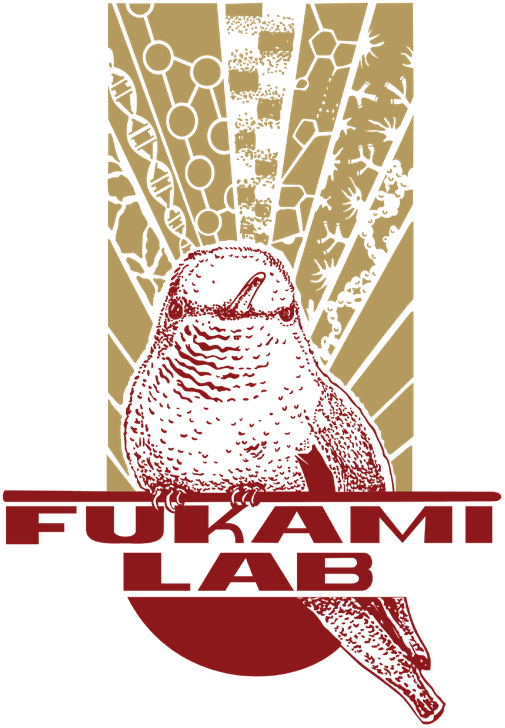

## Supplementary text

### Calculation of generation time

Evolving microbes were transferred 30 times at a dilution of 1/10 (10 uL of each culture was transferred to a fresh 110 mL of nectar). These 30 transfers occurred over 10 weeks (1,680 hours).

To calculate generation time, we estimated the population size of yeast across the experiment by inoculating a series of nectar microcosms with the same initial density of yeast (approximately 2,500 colony forming units (CFUs) and destructively sampling each day.

To calculate generation time, we used the following equation, assuming the yeast growth was exponential before day two:

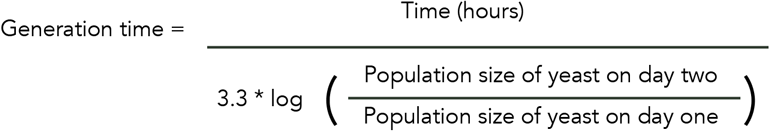

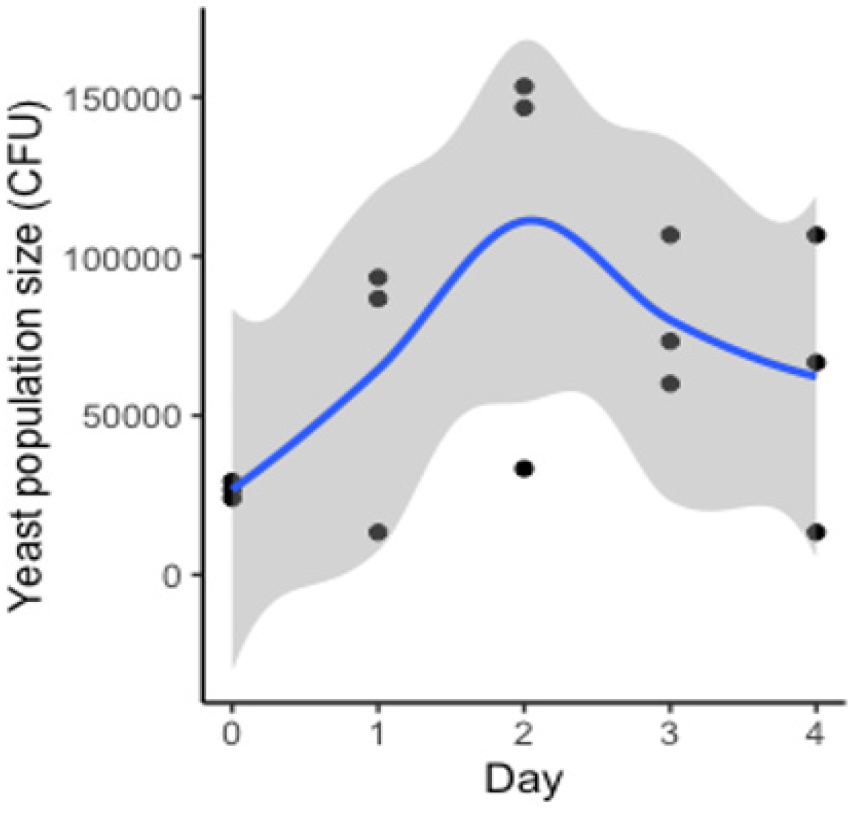

Based on this equation and the data above, we estimate the generation time of *M. reukaufii* yeast (strain MR1) in synthetic nectar is approximately one generation per eight hours. Assuming the generation time stays constant throughout the experiment, we estimate approximately 200 generations over the 30 transfers.

## Supplementary figures

**Figure supplement 1.**
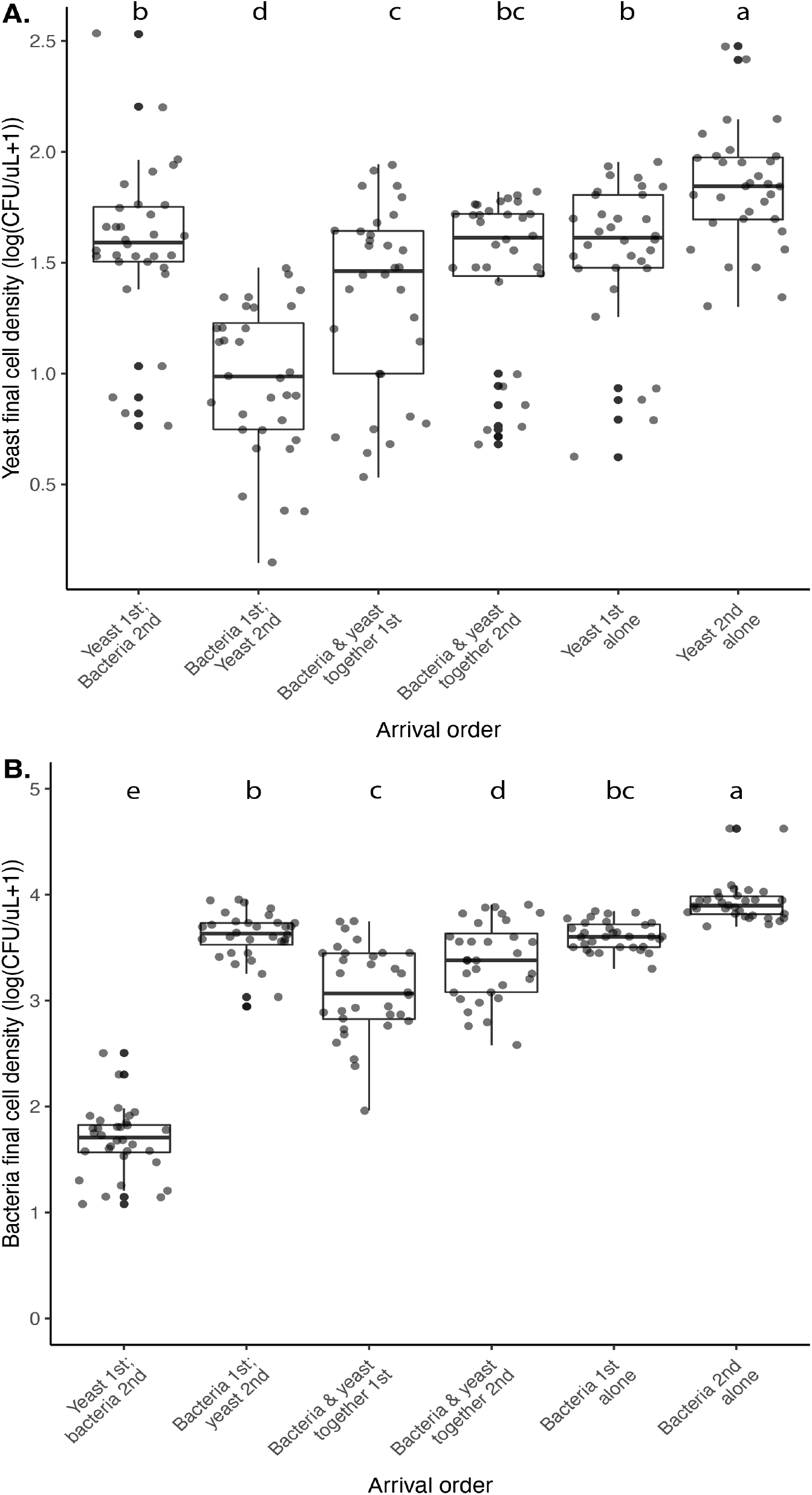
*M. reukaufii* yeast and *A. nectaris* bacteria exhibit negative priority effects against each other, as evidenced by growth in microcosm experiments where arrival order is altered. Statistical analysis for significance codes are in ***Supplementary Table 3*** for yeast (**A**) and bacteria (**B**).

**Figure supplement 2.**
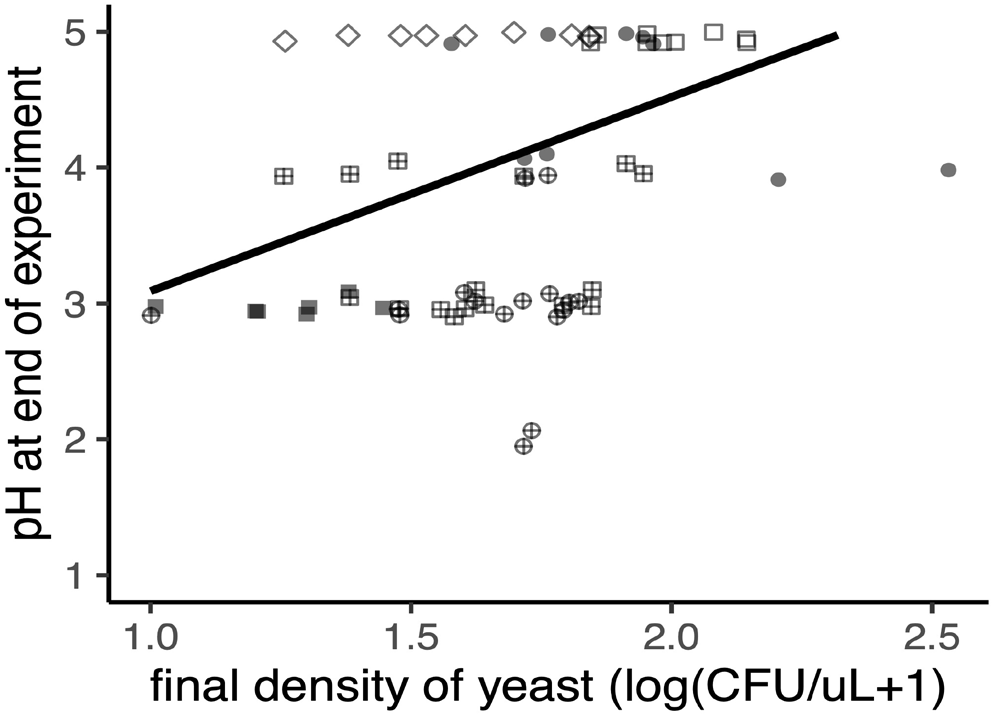
Yeast increases nectar pH (p<0.05, Spearman rank correlation). The shape of each point represents the various treatments (described in ***Figure 3B***), where yeast first, then bacteria (“YB”) are filled circles, bacteria first followed by yeast (“BY”) are filled squares, early arriving bacteria and yeast (“YB-”) are cross-hatched squares, late arriving bacteria and yeast (“BY-”) are cross-hatched circles, late arriving yeast (“-Y) are open squares, early arriving yeast (“Y-”) are open diamonds, late arriving bacteria (“-B”) are open circles, and early arriving bacteria (“B-”) are triangles. Points are slightly jittered on the y axis.

**Figure supplement 3.**
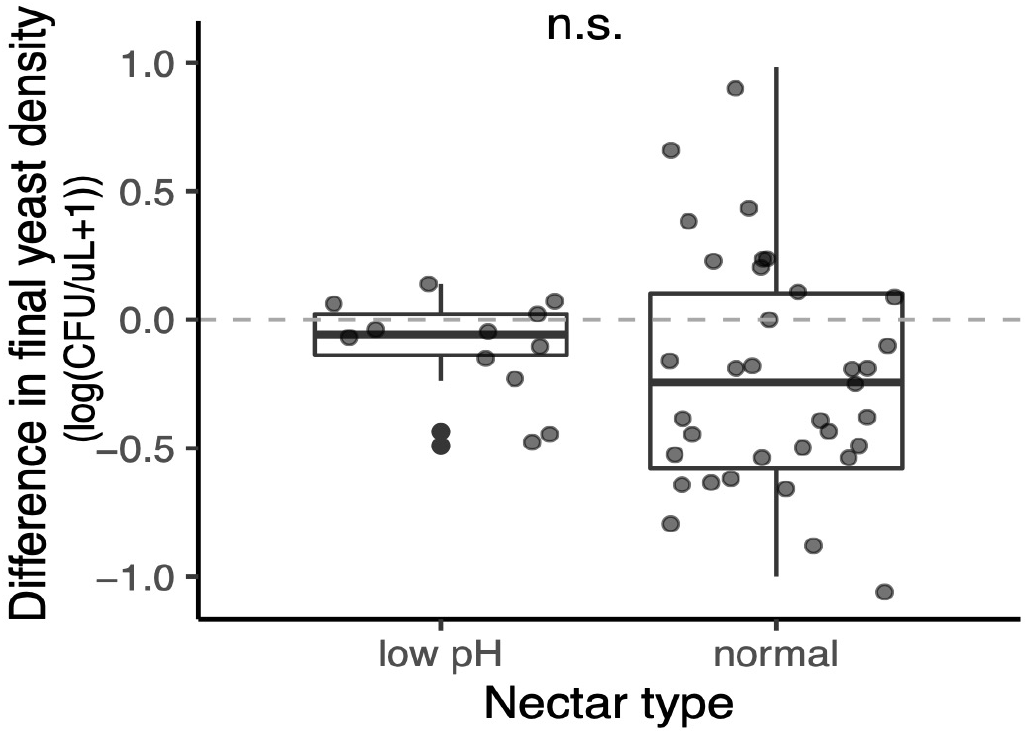
We found no effect of nectar type (pH=3, pH=6) on the growth of *M. reukaufii*, when grown in monoculture at a high density (approximately 10,000 cells/μL). *M. reukaufii* growth was calculated by subtracting the final from initial cell density.

**Figure supplement 4.**
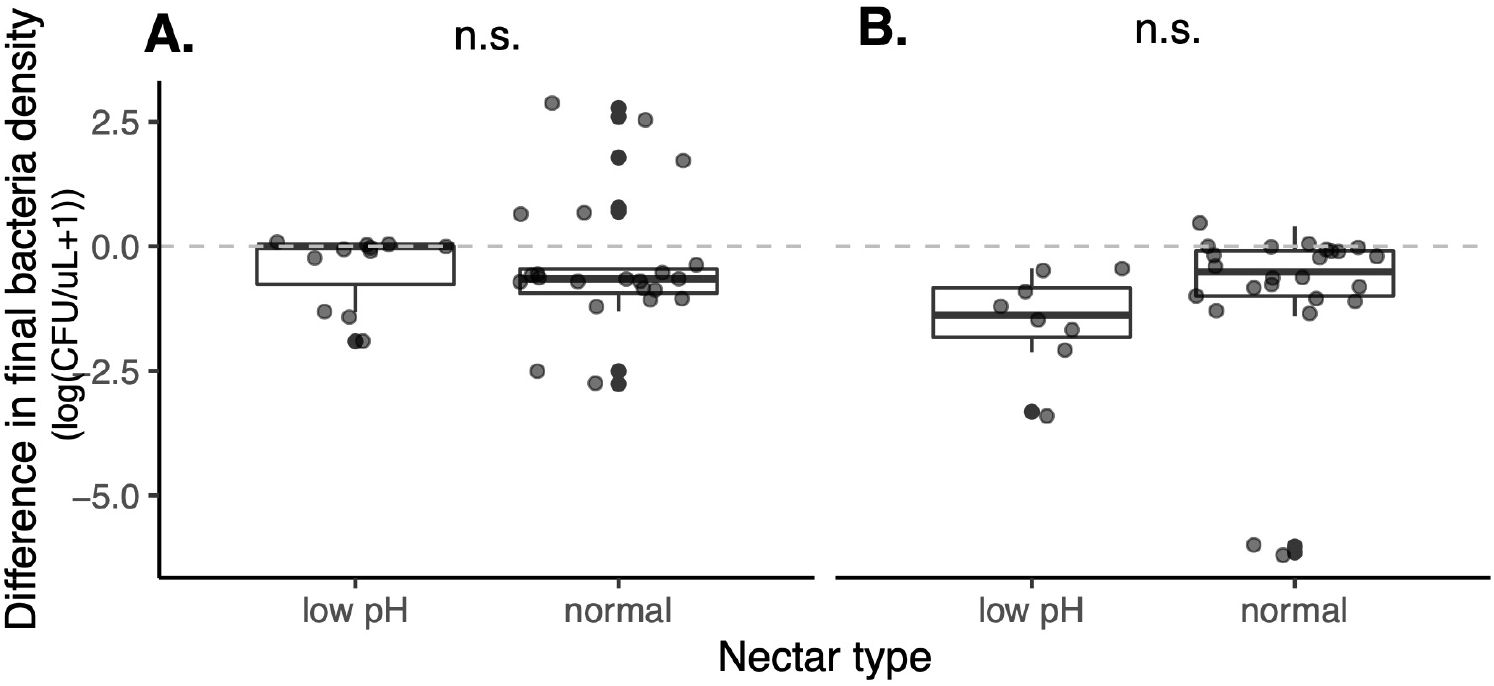
We found no effect of nectar type (pH=3, pH=6) on the growth of *A. nectaris* when grown in monoculture at a low density (approximately 10 cells/μL) (**A**) or high density (approximately 10,000 cells/μL) (**B**). *A. nectaris* growth was calculated by subtracting the final from initial cell density.

**Figure supplement 5.**
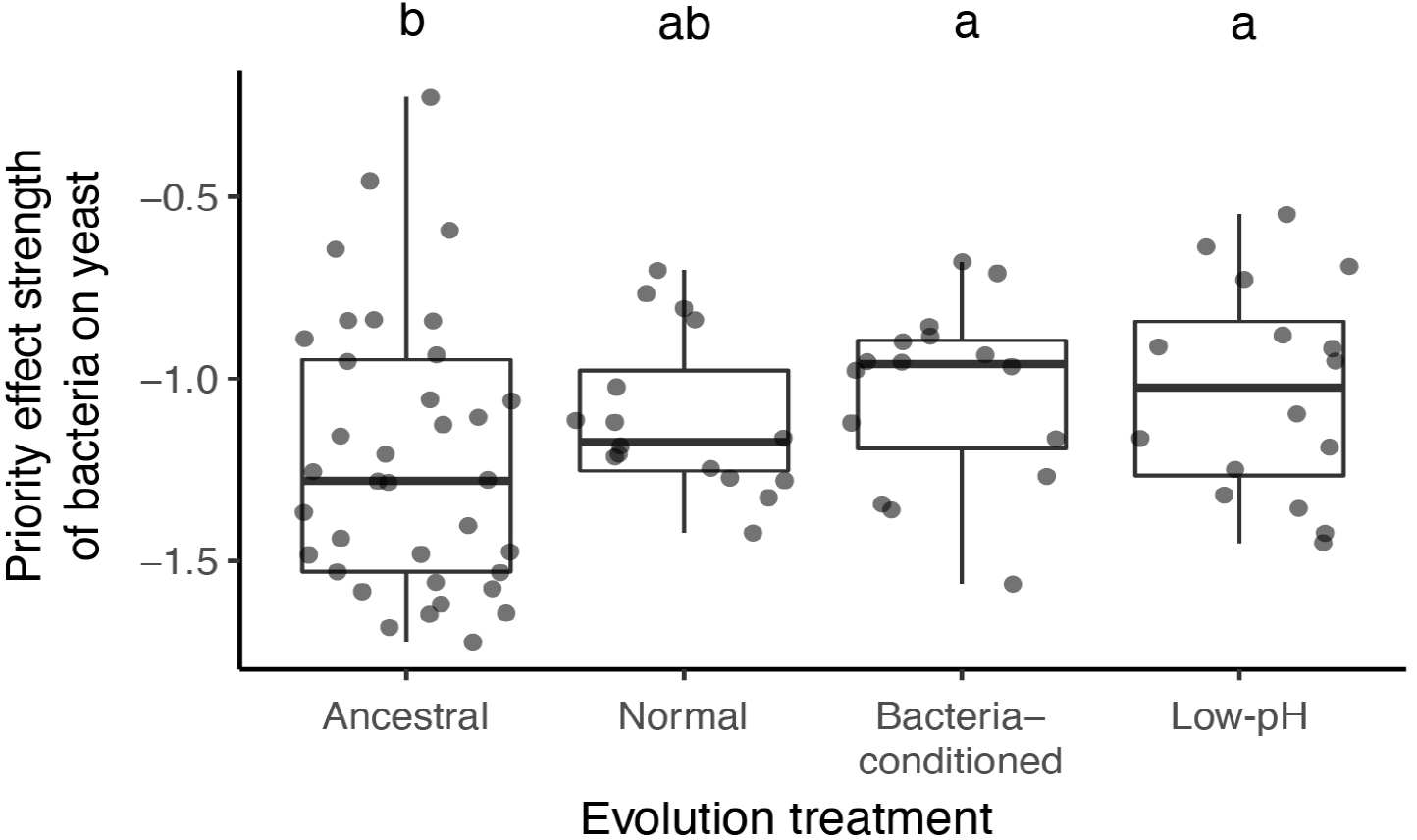
We calculated an additional priority effect metric, which corroborated our main result. This metric is calculated by taking the natural logarithm of growth ratio between different initial dominance: PE = log(BY/YB).

**Figure supplement 6.**
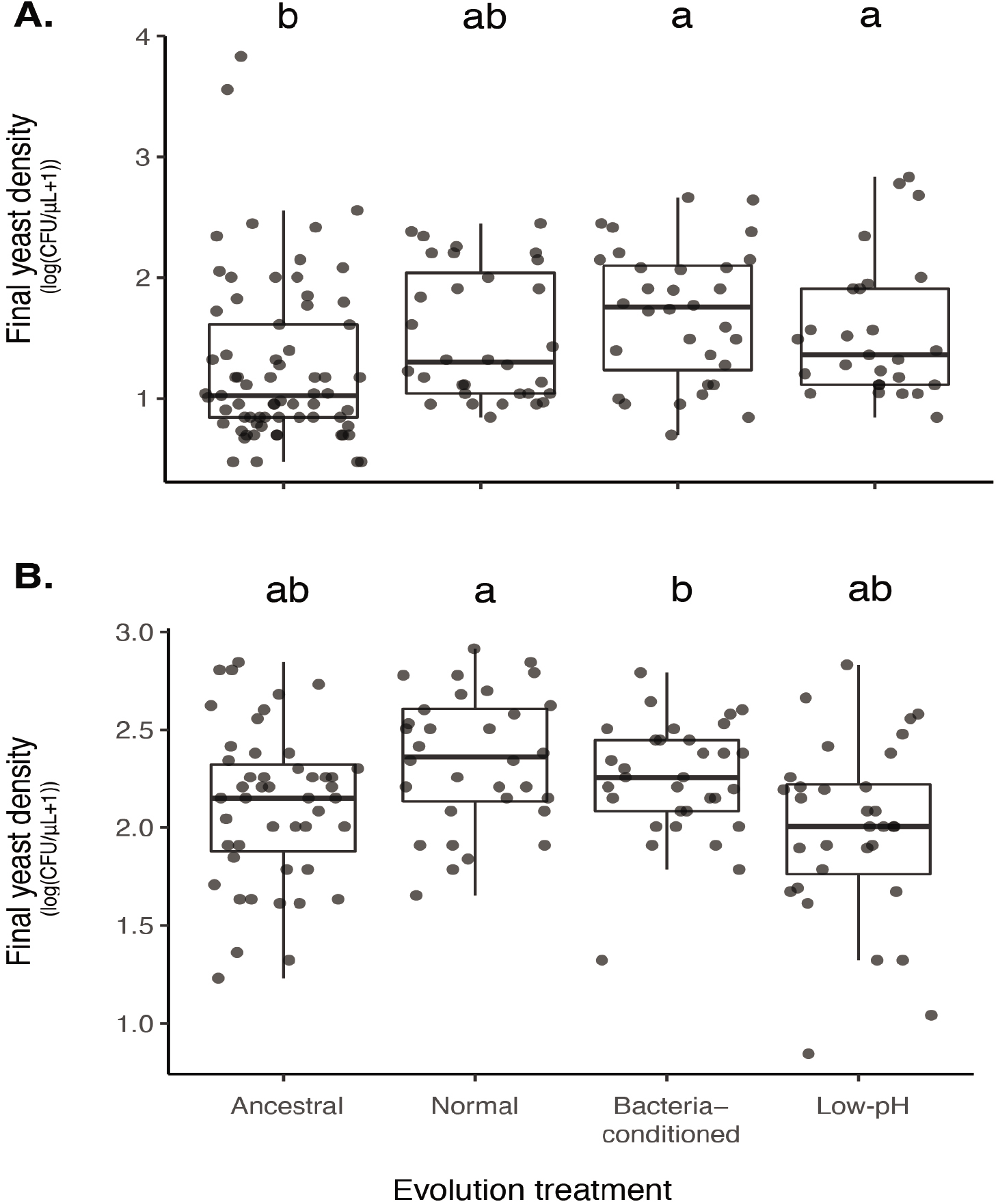
(**A**) Yeast evolved in bacteria-conditioned and low-pH nectars were better able to grow than ancestral yeast when bacteria were dominant (BY bacterial priority effects treatment). (**B**) This difference in growth was not due to a difference in intrinsic growth rate (-Y monoculture treatment).

**Figure supplement 7.**
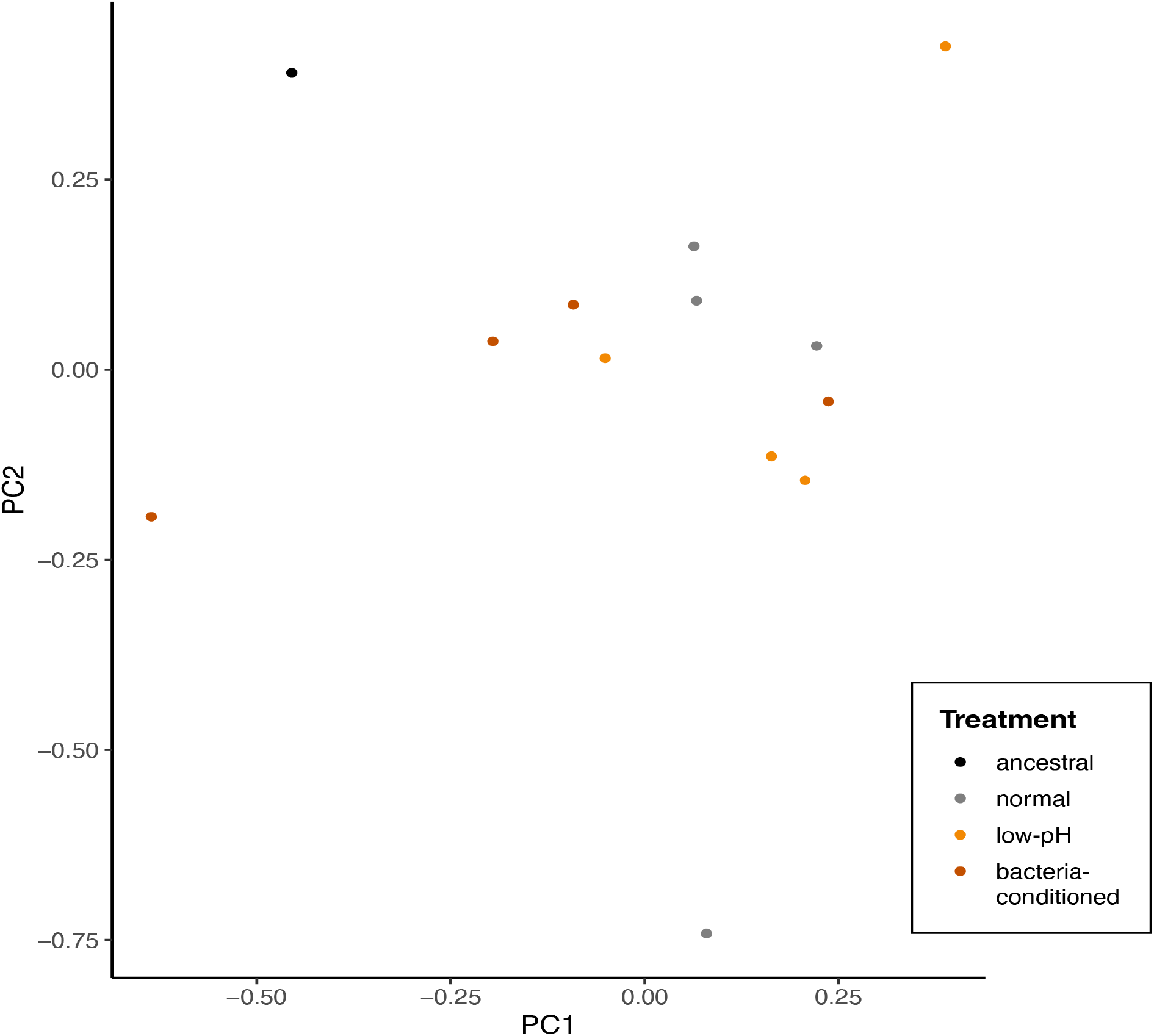
Principal components analysis of single nucleotide polymorphisms that differ between evolved and any ancestral strain (2,319 sites), points represent individual end-point clones (individuals) and are colored by evolution treatment (ancestral in black, neutral in grey, evolved in low-pH nectar in orange, and evolved in bacteria-conditioned nectar in dark red). (perMANOVA, p=0.19, n=12).

**Figure supplement 8.**
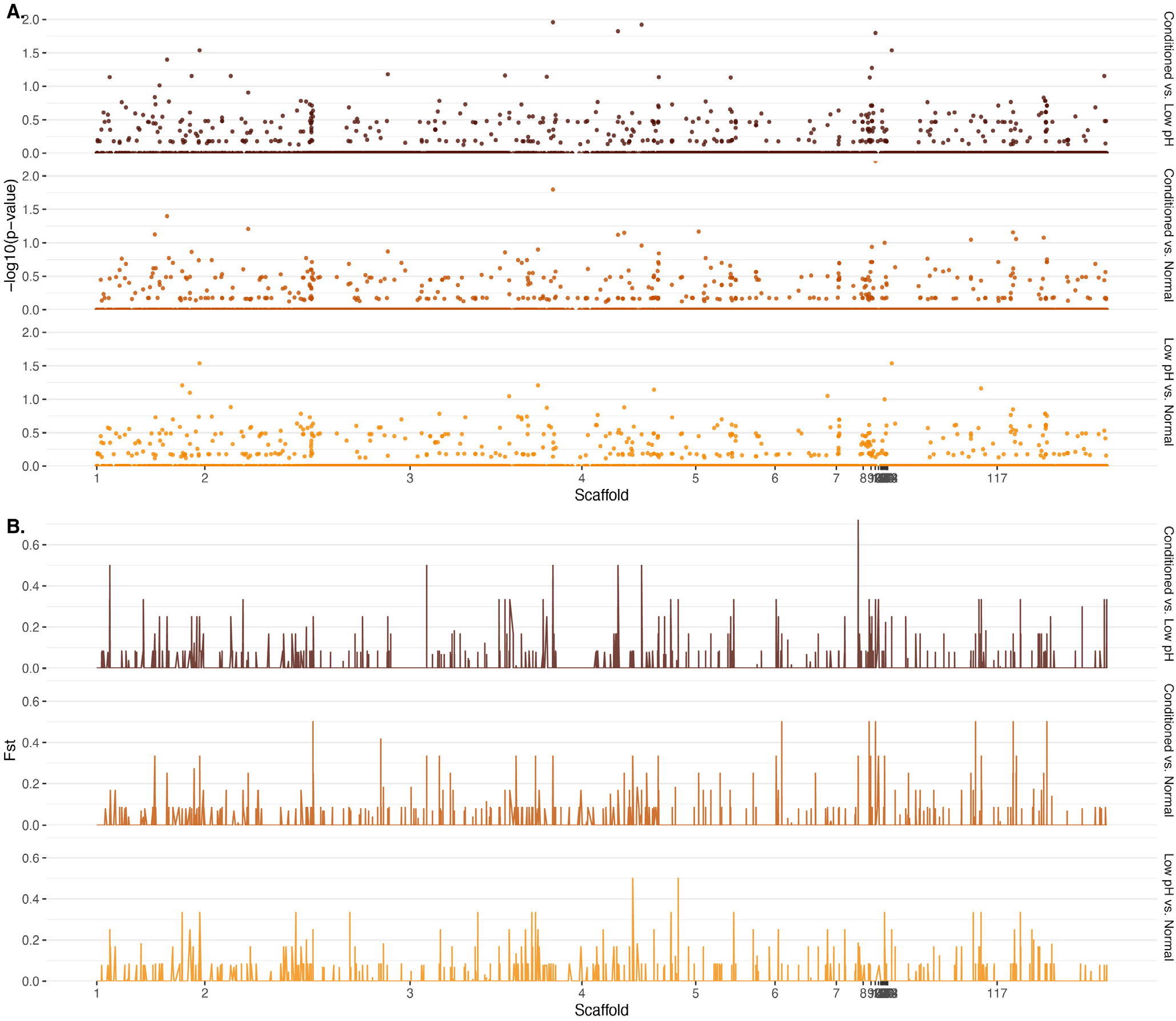
Genome-wide differentiation by (**A**) computed p-values for observed patterns of loss of heterozygosity (LOH) and (**B**) Weir-Cockerham estimator of F_ST_.

**Figure supplement 9.**
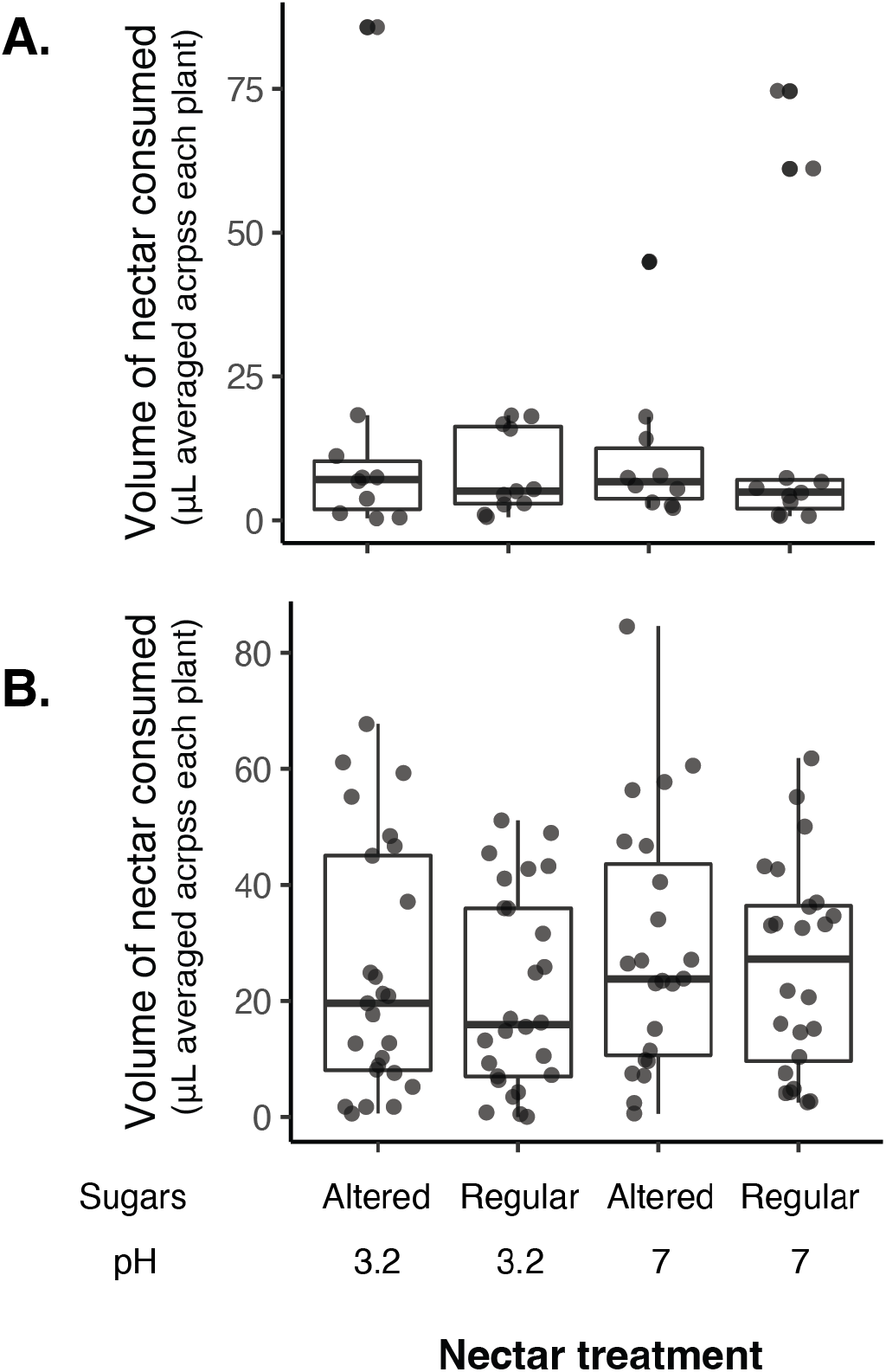
No difference in the volume of nectar consumed by nectar treatment in 2016 (**A**) or 2018 (**B**) when nectar sugars were altered with pH.

## Supplementary tables

**Supplementary table 1 - Priority effect experiment treatments.**
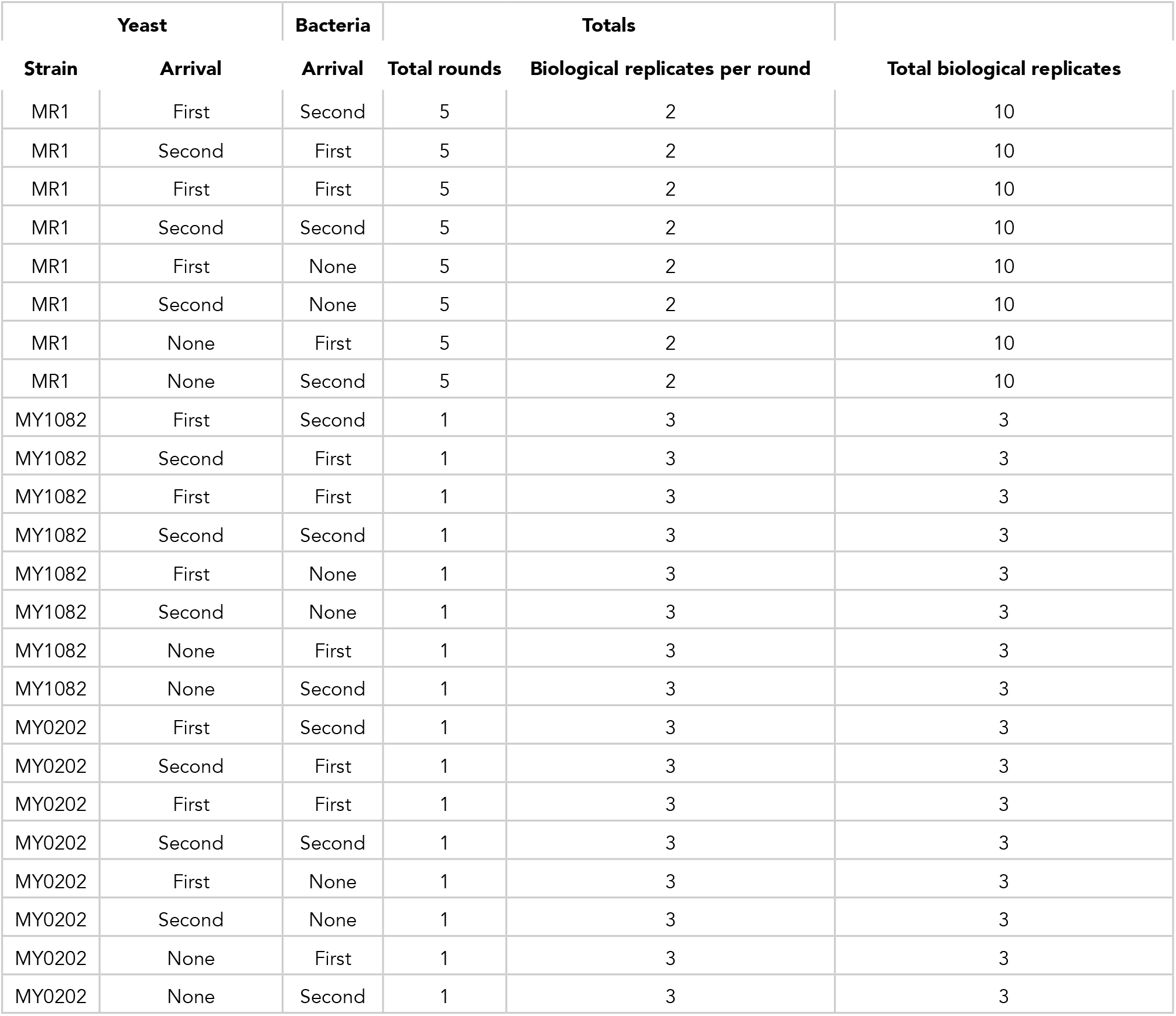
Treatments used in priority effects experiment with ancestral yeast, fully factorial experiment testing the effect of arrival order on priority effects.

**Supplementary table 2 - Evolution treatments.**
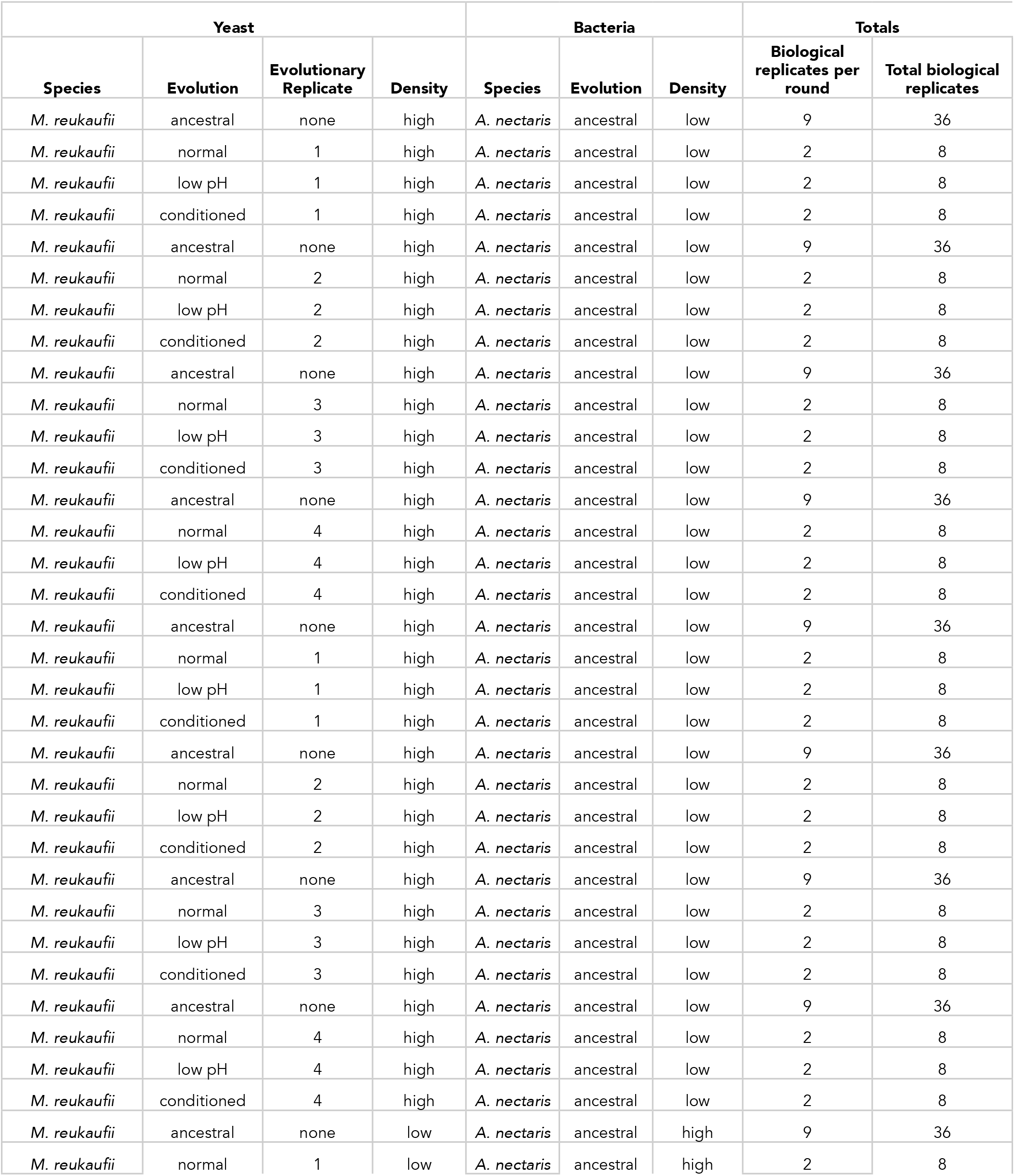
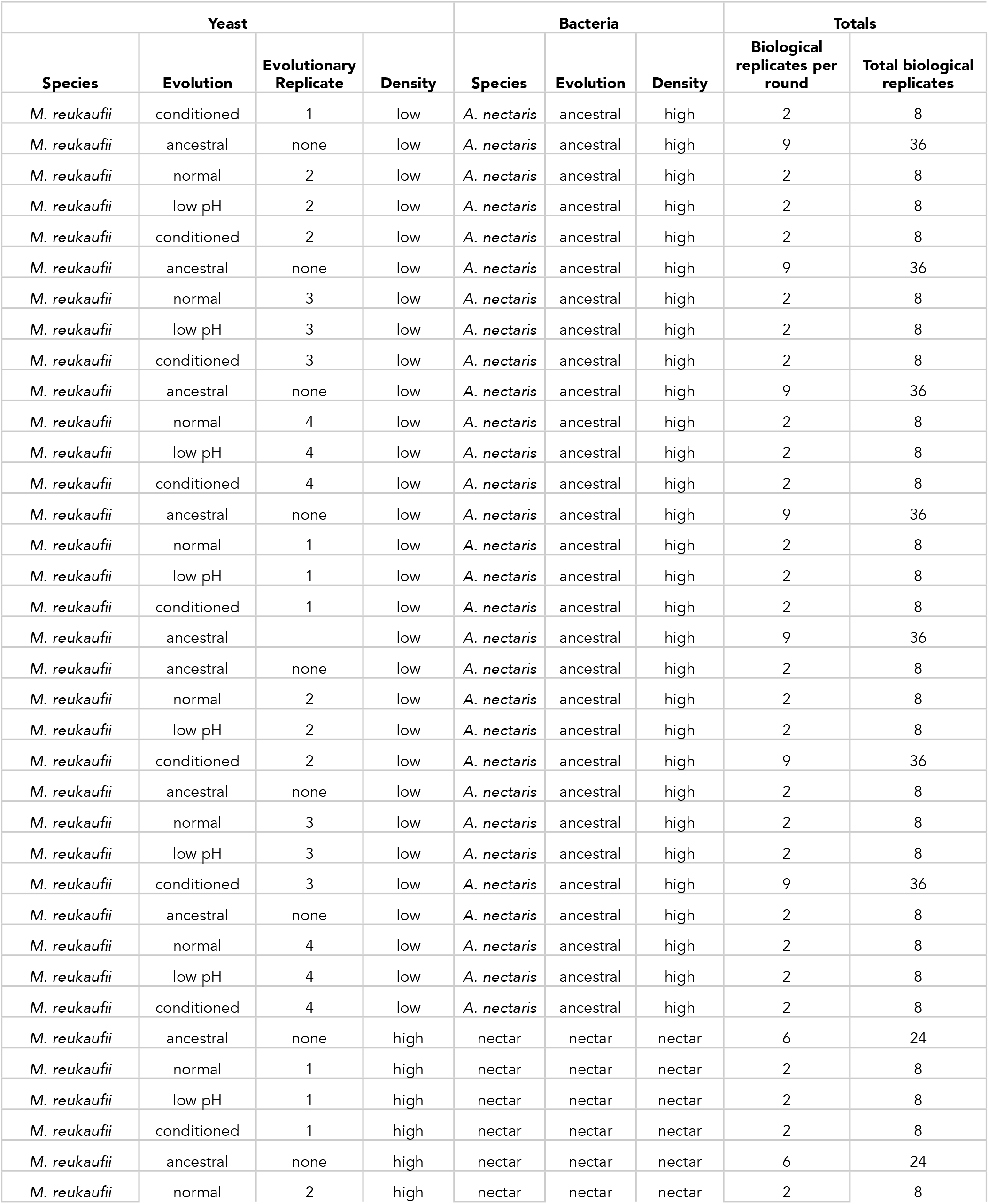
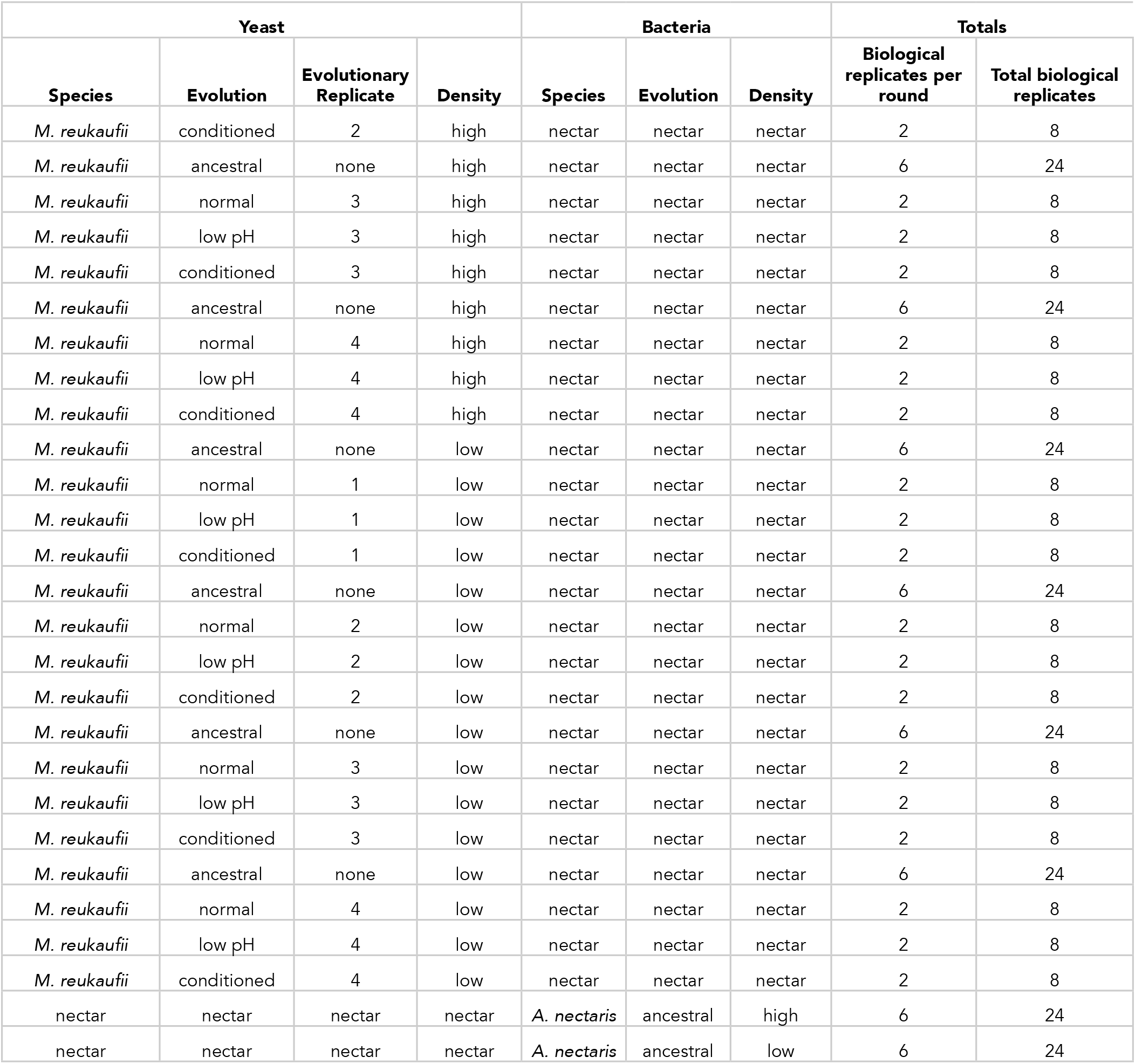
Treatments used in priority effects experiment with experimentally-evolved yeast, fully factorial experiment testing the effect of initial density (10,000 cells/μL (“early”) or 10 cells/μL (“late”)), evolution treatment (ancestral, evolved in normal nectar, low-pH nectar, or bacteria-conditioned nectar), and evolutionary replicate (independent evolutionary lineages) on priority effects.

**Supplementary table 3 - Priority effect experiment results.**
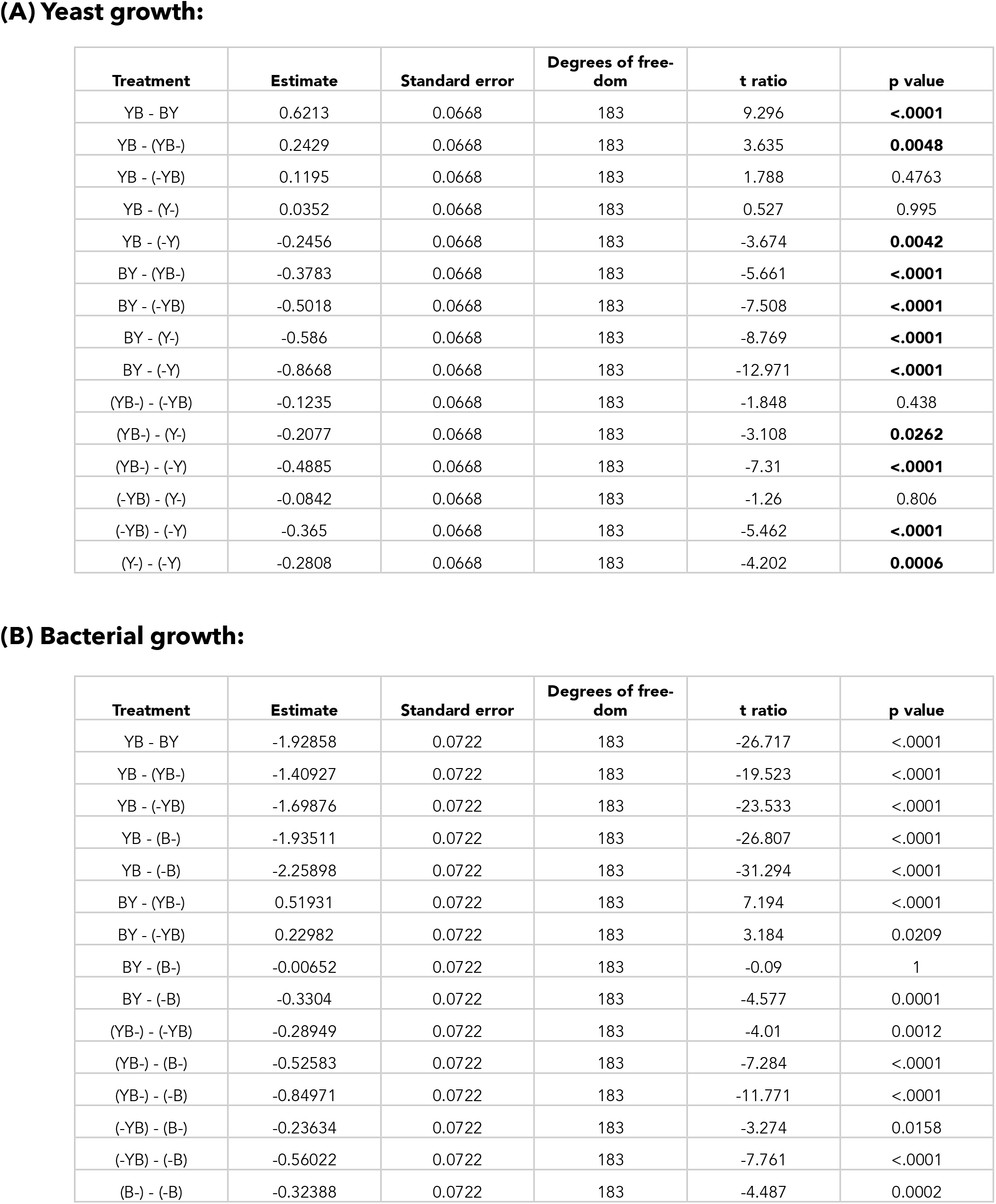
Results from a linear mixed model testing the effect of arrival order on yeast growth, where BY and YB represents initial arrival by bacteria or yeast, respectively. -Y and Y-represent the comparable growth of yeast at either arrival time (day 0 or day 2). Bold text shows p-values less than or equal to 0.05.

**Supplementary table 4 - Priority effect experiment results with wild strains.**
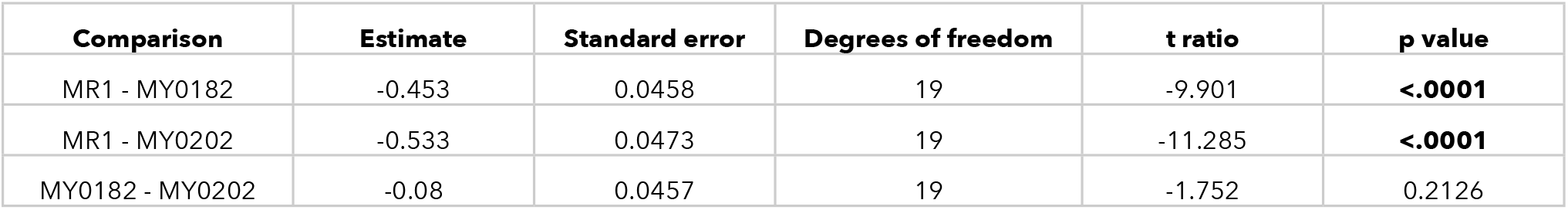
Results from a linear mixed model testing the effect of yeast strain (MR1, MY0182, MY0202) on the strength of priority effects exerted by *A. nectaris* calculated using this metric: PE = log(BY/(-Y)) - log(YB/(Y-)), where BY and YB represents initial dominance by bacteria or yeast, respectively. -Y and Y-represent the comparable growth of yeast at either density, alone and treatment densities were averaged by round of the experiment. Bold text shows p-values less than or equal to 0.05.

**Supplementary table 5 - Priority effect experiment results with evolved strains.**
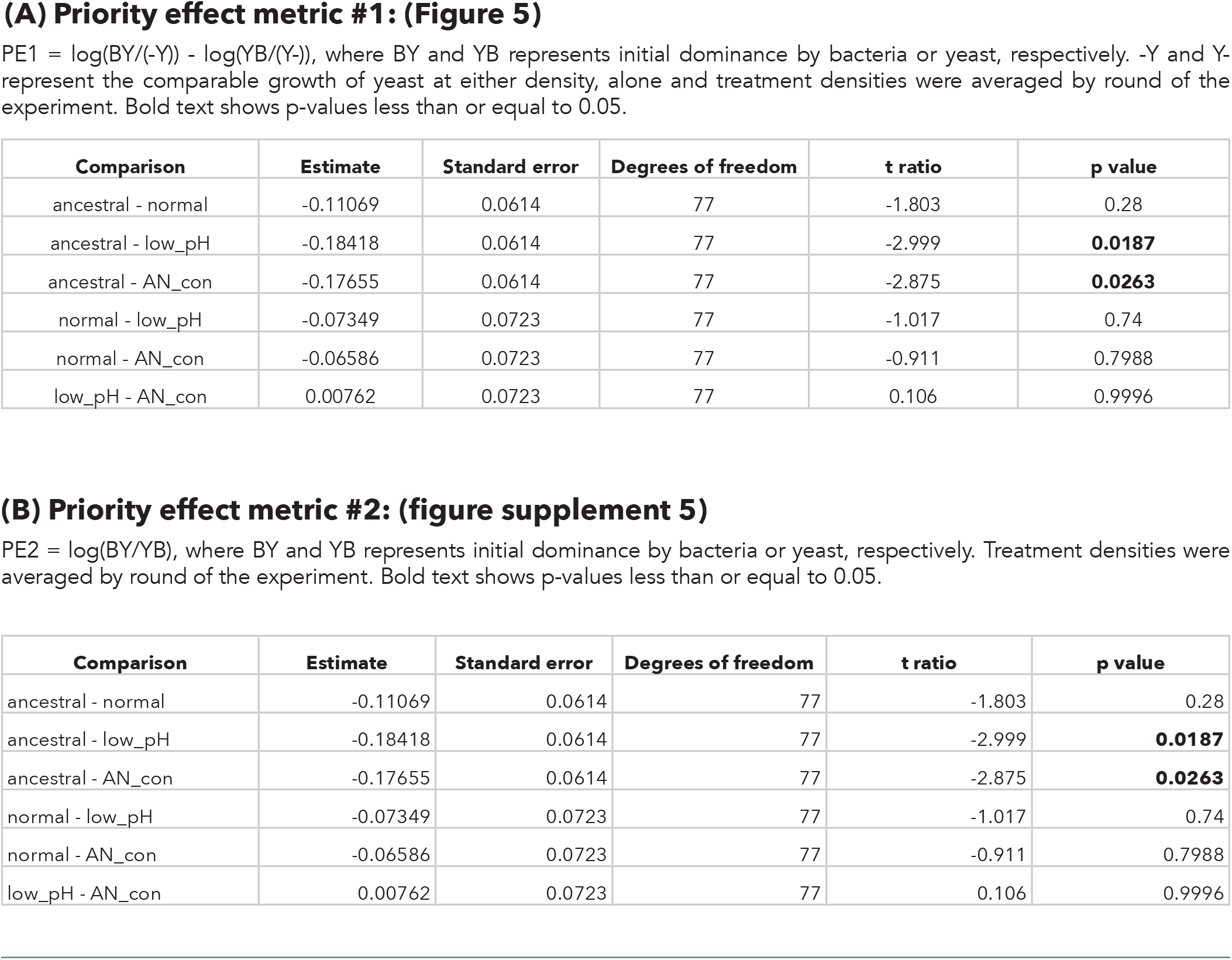
Results from a linear mixed model testing the effect of evolution treatment (ancestral, evolved in normal nectar, low-pH nectar, or bacteria-conditioned nectar) on the strength of priority effects calculated using one of two metrics:

**Supplementary table 6 - Differences in growth between evolved strains with and without bacteria.**
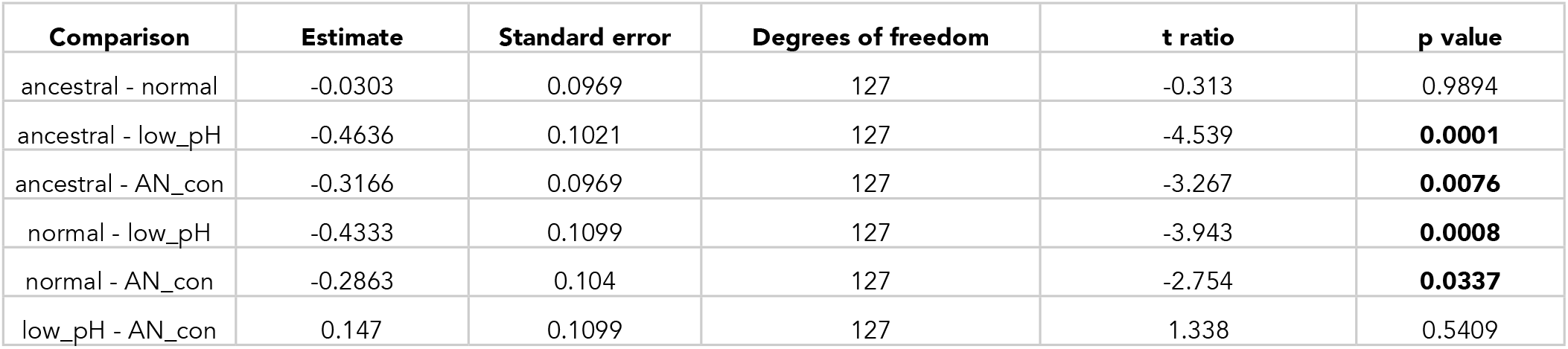
Results from a linear mixed model testing the effect of yeast initial density (10,000 colony forming units/μL (“high”) or 10 cells/ μL (“low”), yeast monoculture or competition with bacteria) treatment and evolution treatment (ancestral, evolved in normal nectar, low-pH nectar, or bacteria-conditioned nectar) on the difference in final yeast density between treatments with a high density of bacteria and a low density of yeast (BY) and yeast grown in monoculture at a low density (-Y). Growth difference was calculated as (BY) - (-Y). Bold text shows p-values less than or equal to 0.05.

**Supplementary table 7 - Difference in final yeast densities between evolved strains with and without bacteria.**
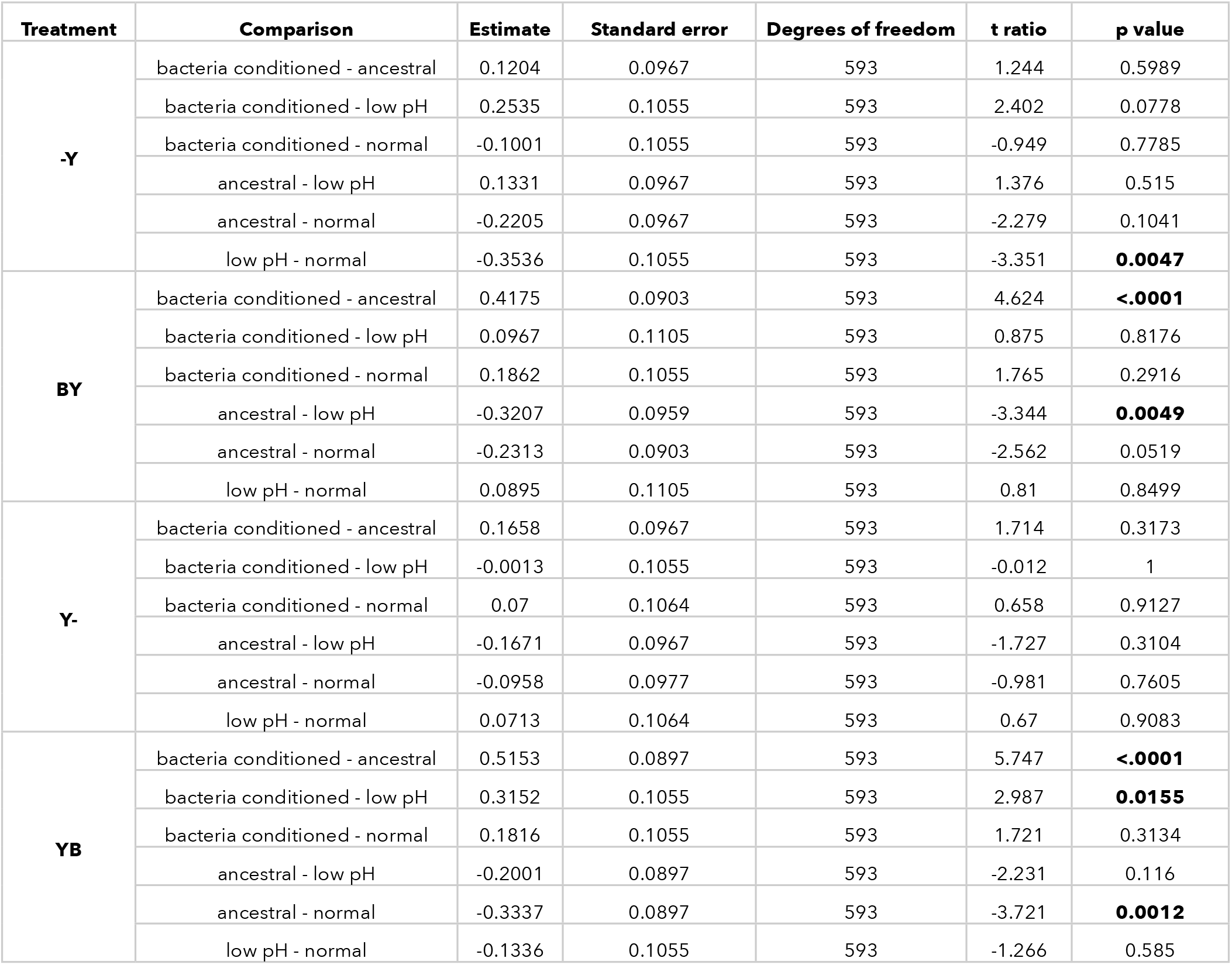
Results from a linear mixed model testing the effect of yeast initial density (10,000 colony forming units/μL (“early”) or 10 cells/ μL (“late”), yeast monoculture or competition with bacteria) treatment and evolution treatment (ancestral, evolved in normal nectar, low-pH nectar, or bacteria-conditioned nectar) on final density of yeast. Bold text shows p-values less than or equal to 0.05.

**Supplementary table 8 - Sequencing and mapping QC.**
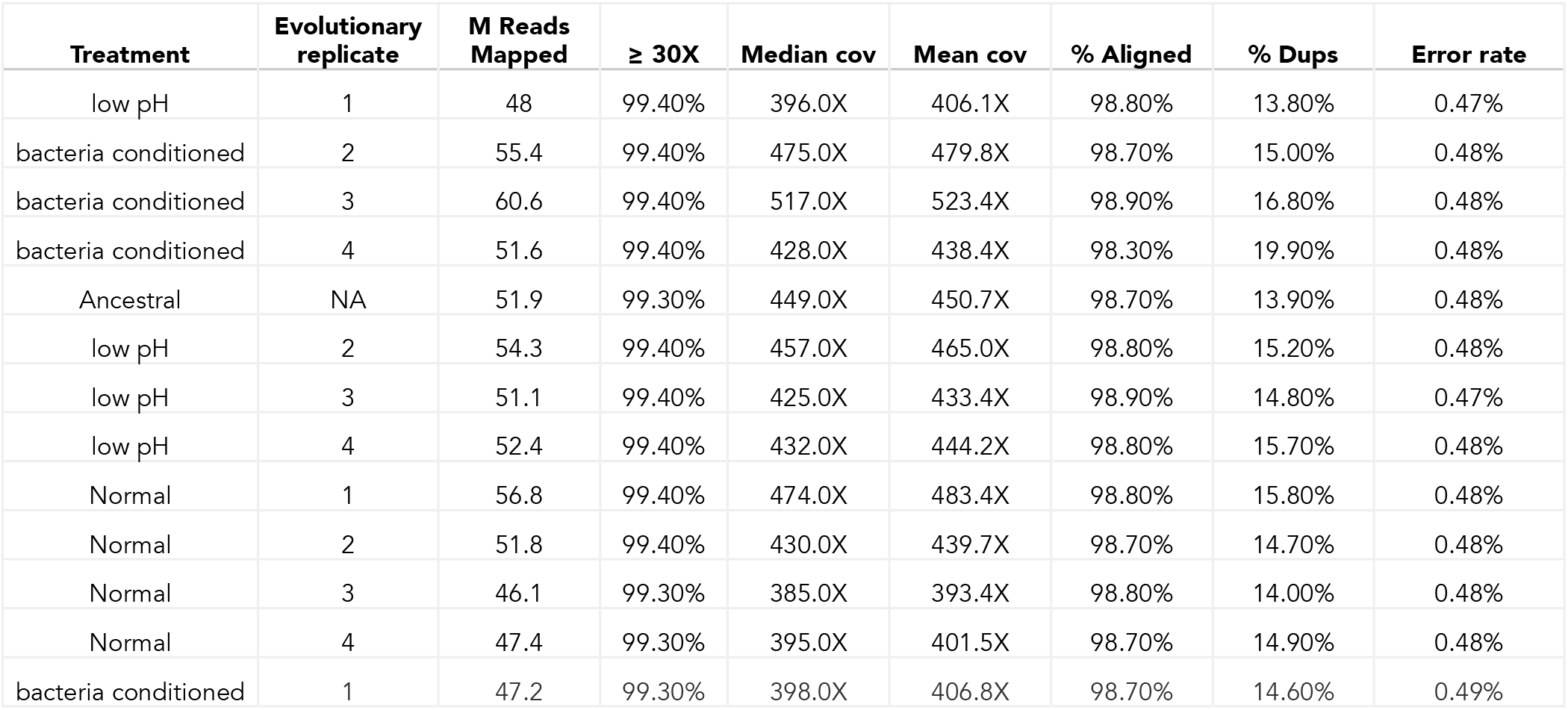
Mapping summary of evolved and ancestral genomes. [Link](https://drive.google.com/file/d/1RftIbyph1yBWrNV5xul38FvX0cztztQ_/view) to full MultiQC report. There was a total of 675M mapped reads and an average coverage depth of 444X.

**Supplementary table 9 - Nearest annotated gene with treatment-specific divergence.**
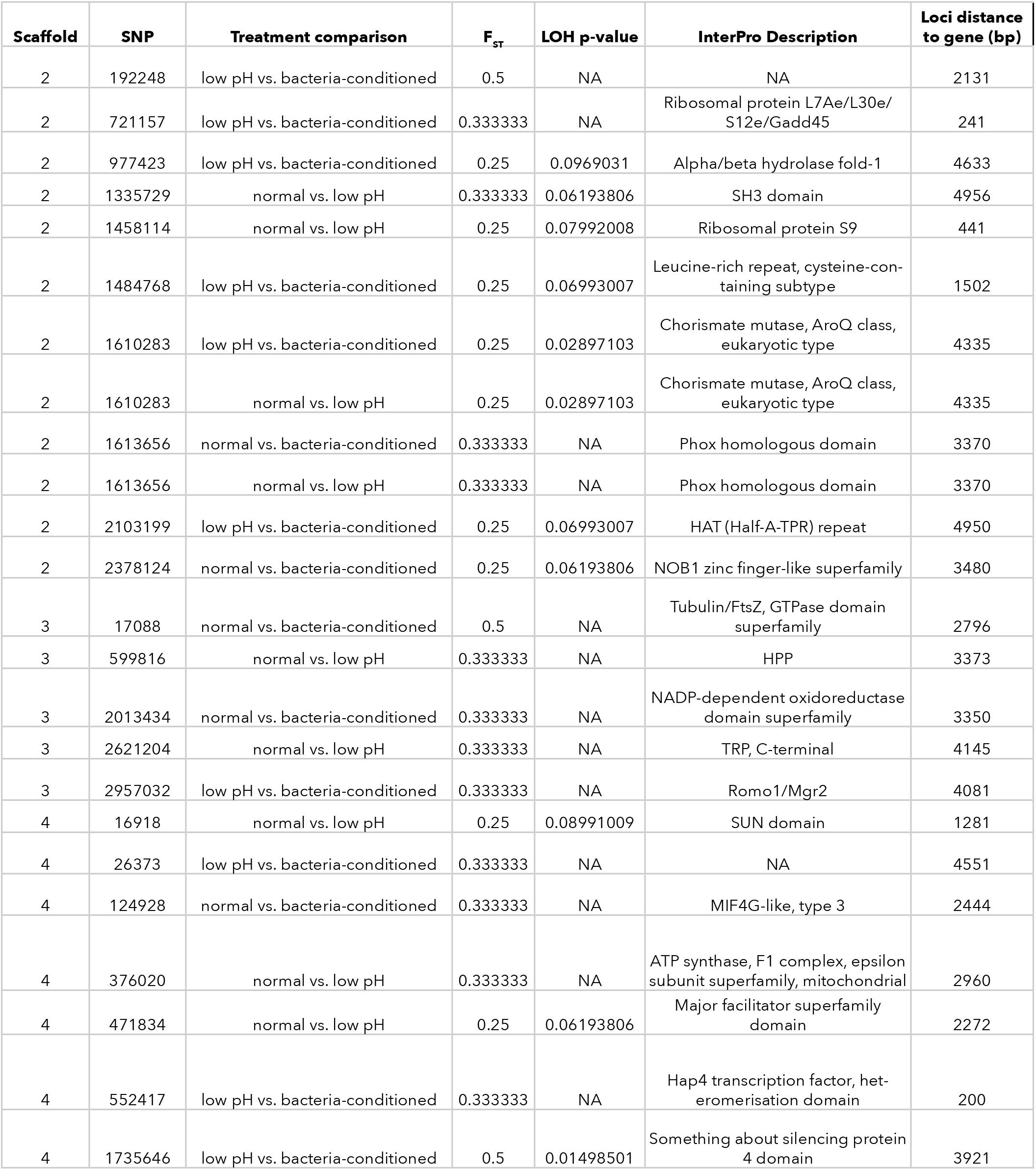
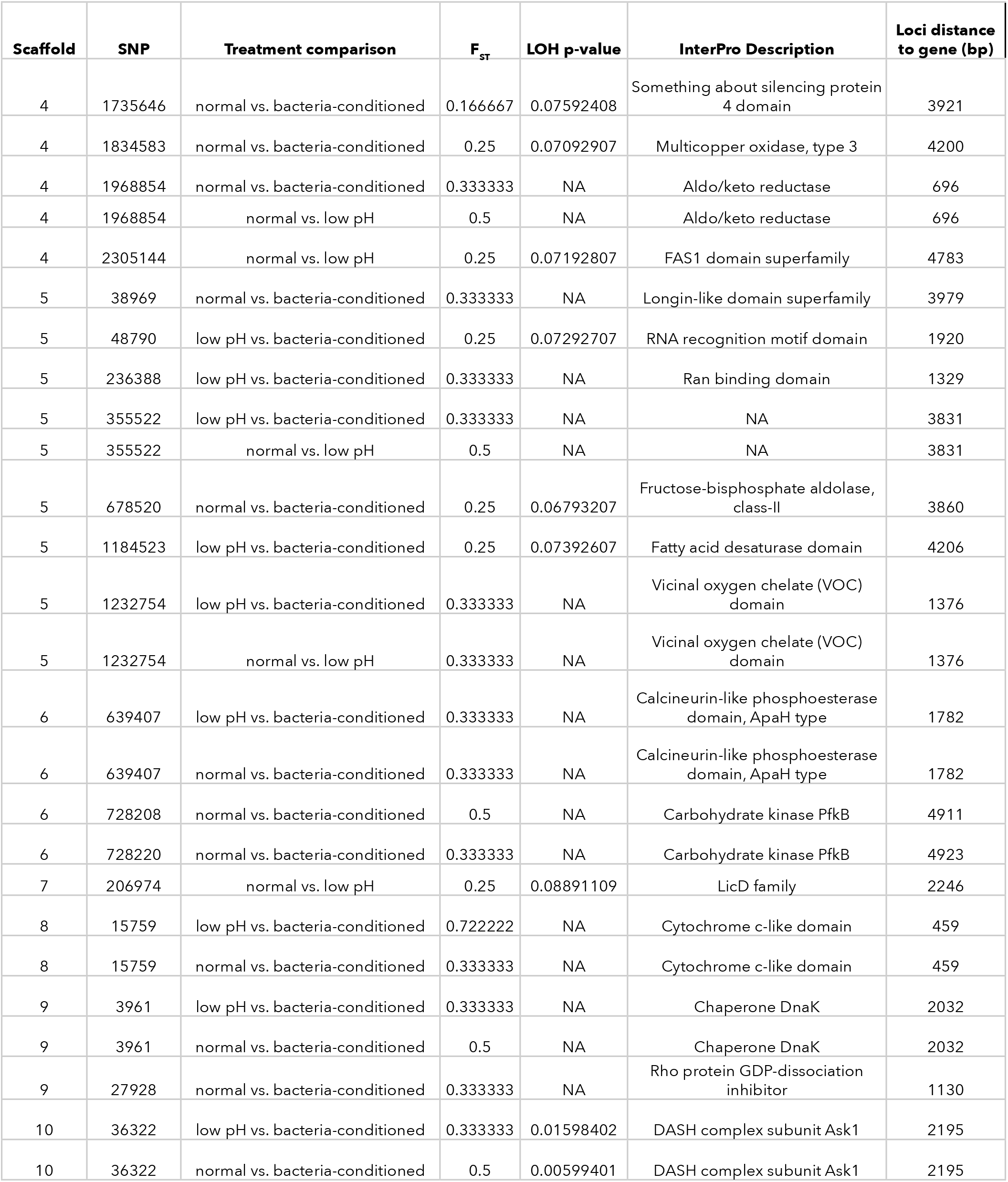
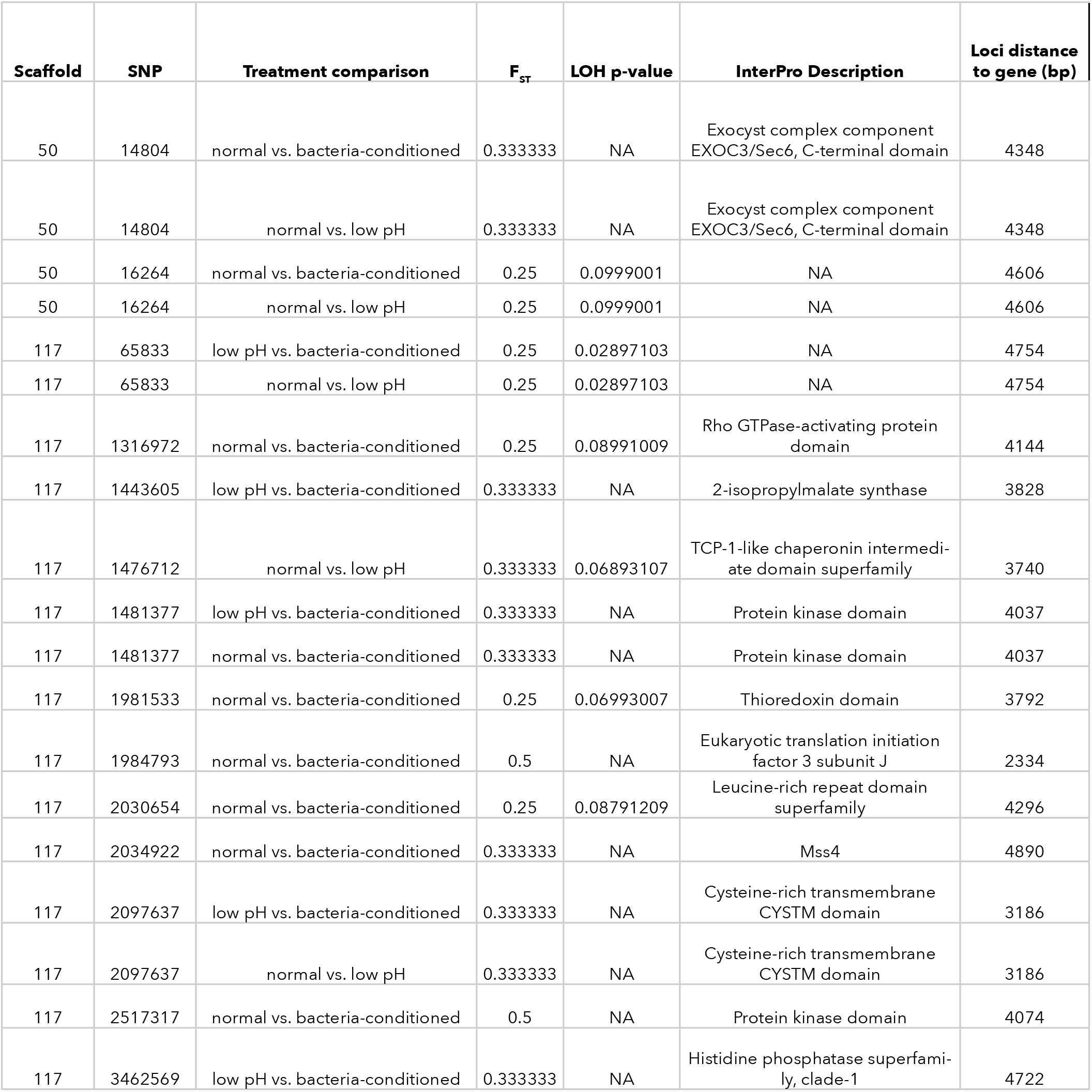
Nearest annotated gene to each loci with treatment specific divergence (loss of heterozygosity permutation test) with p<0.1 and F_ST_ = 0.3-0.5.

**Supplementary table 10 - Nearest annotated gene with punitive de novo singleton mutation.**
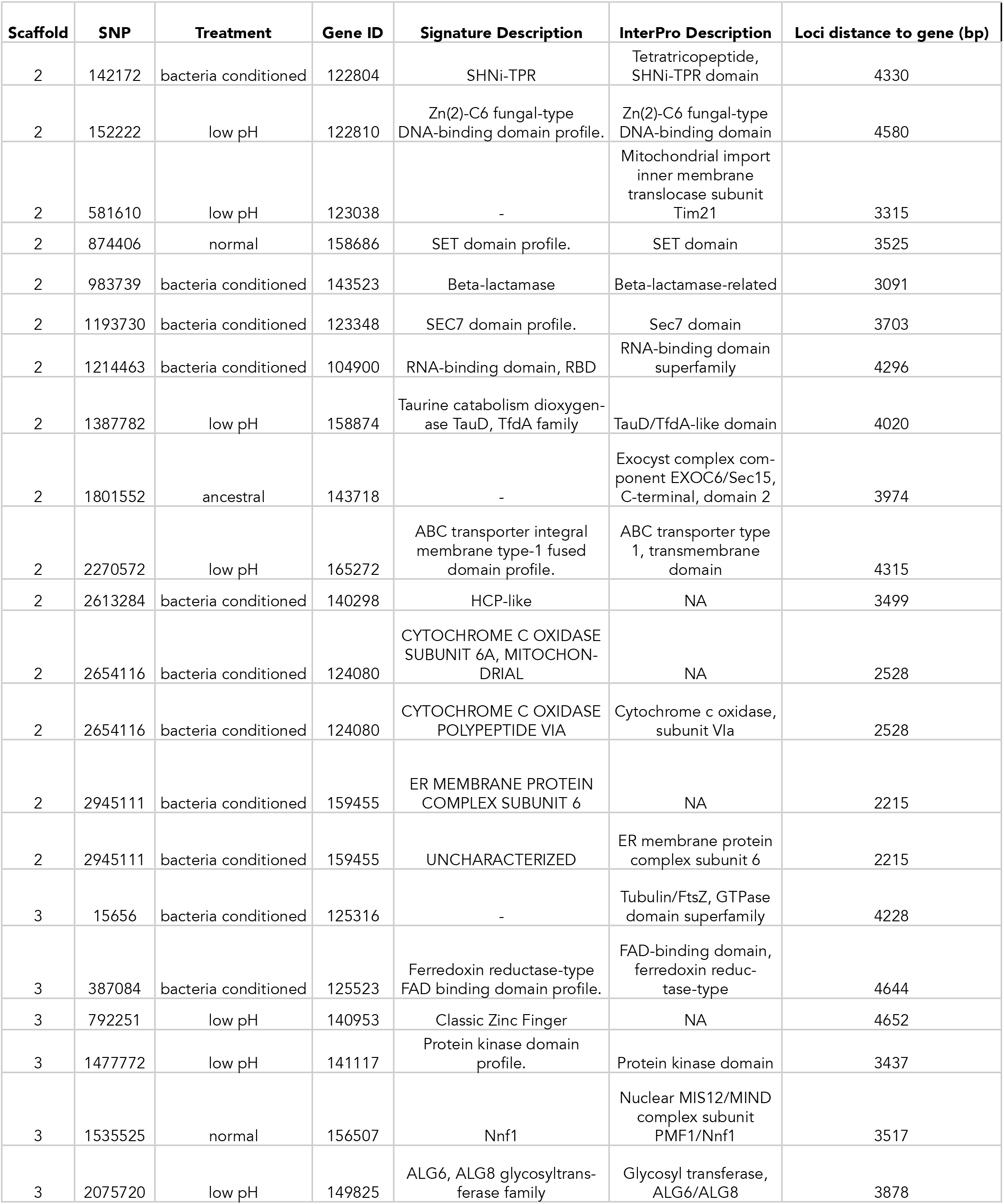
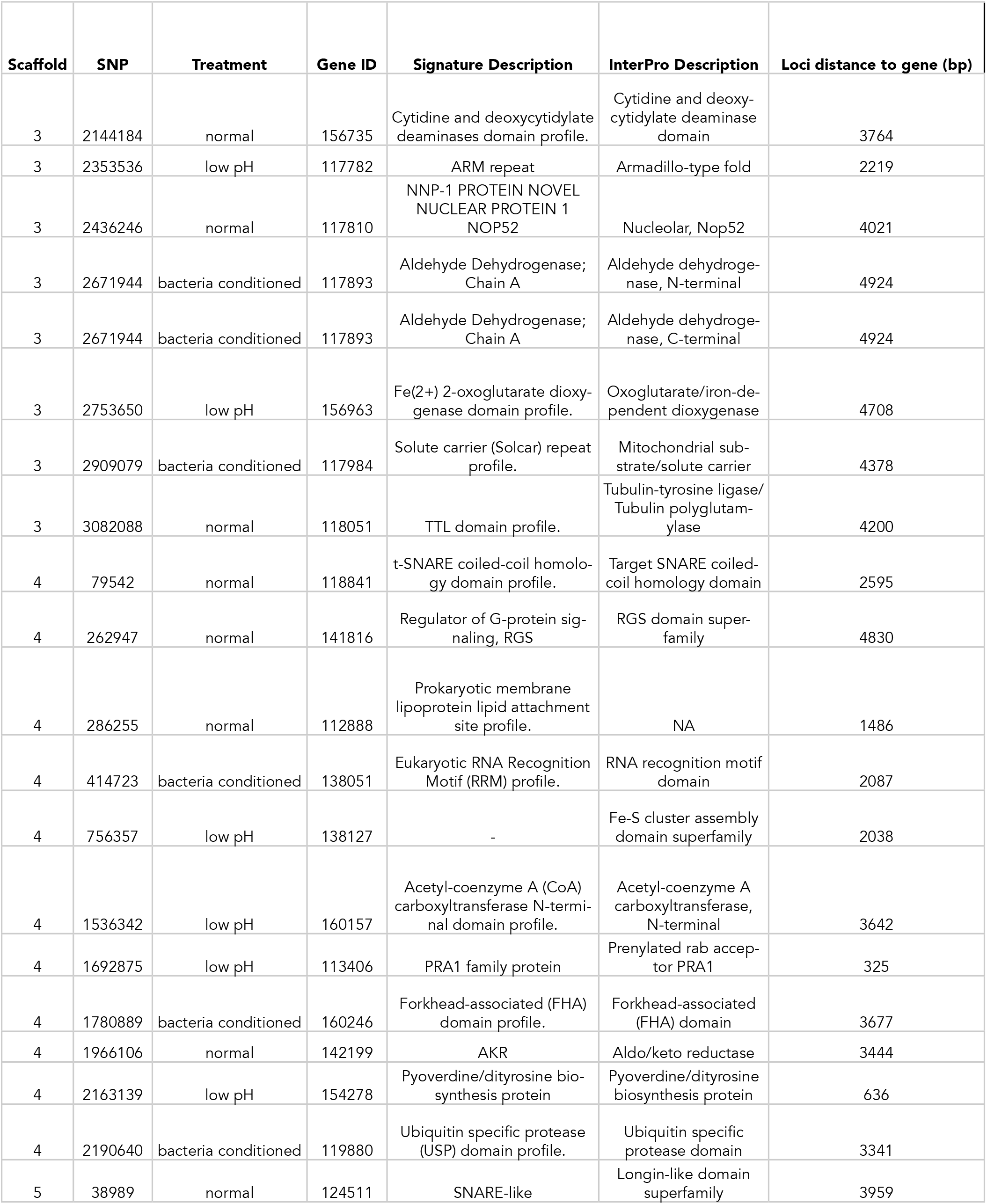
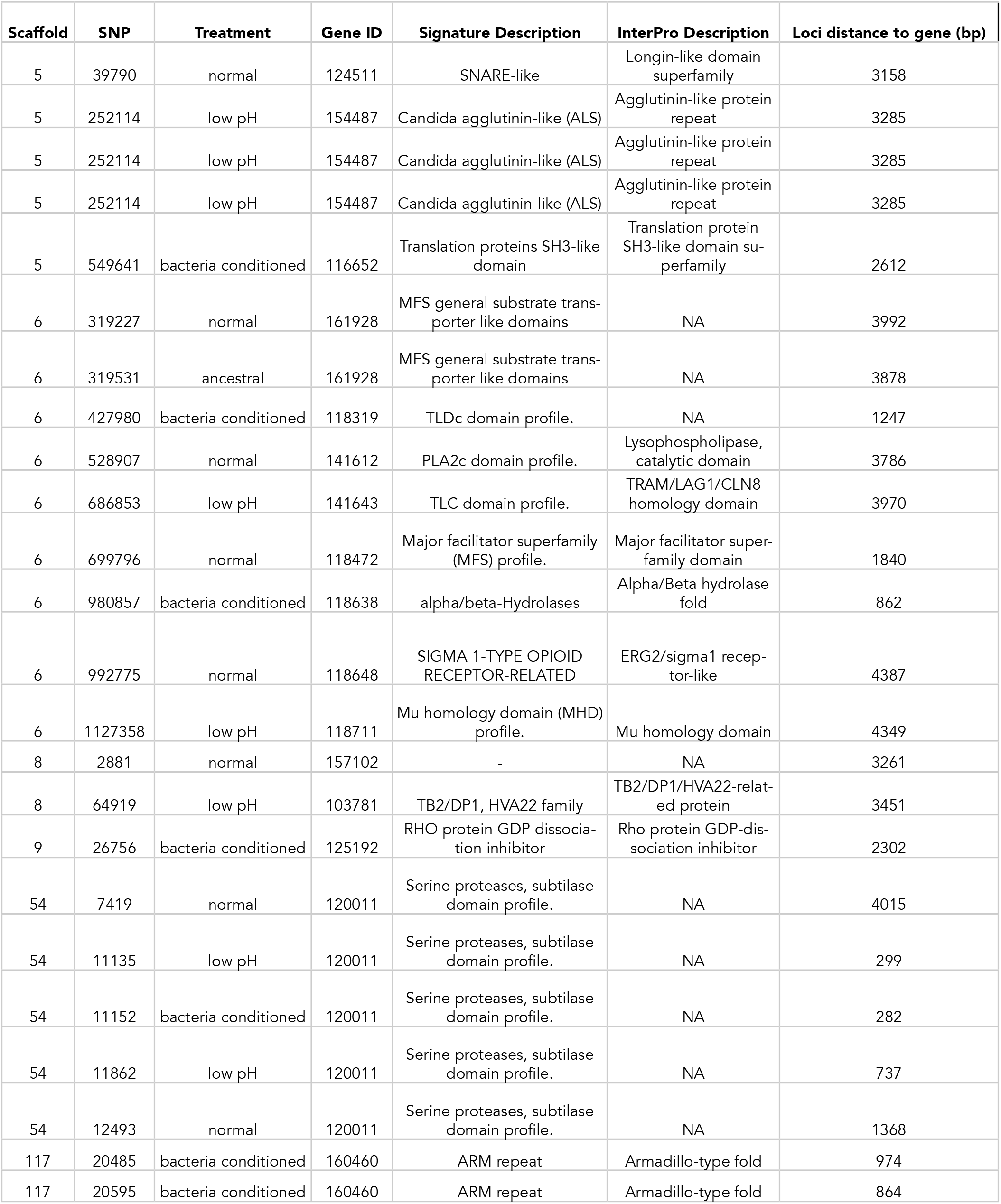
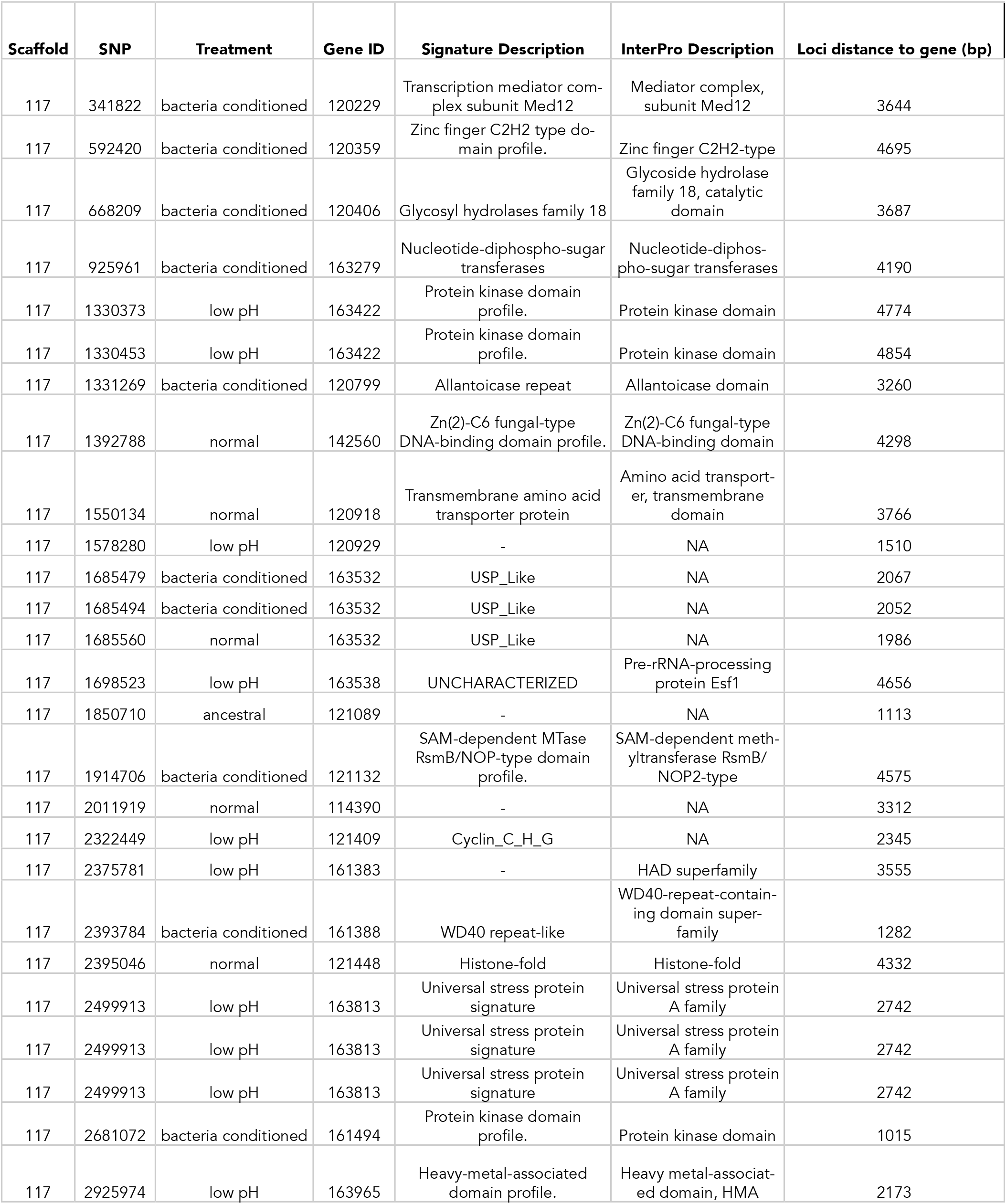
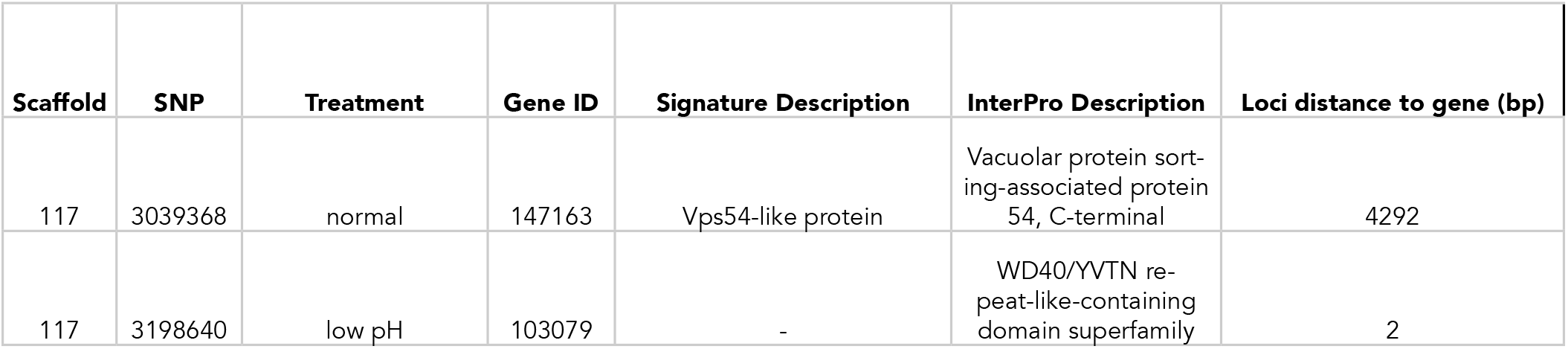
Nearest annotated gene to each putative *de novo* singleton mutation.

