## Supplementary material for "pH as an eco-evolutionary driver of priority effects": Chappell et al. - supplement

### Supplementary text

#### Calculation of generation time

Evolving microbes were transferred 30 times at a dilution of 1/10 (10 uL of each culture was transferred to a fresh 110 mL of nectar). These 30 transfers occurred over 10 weeks (1,680 hours).

To calculate generation time, we estimated the population size of yeast across the experiment by inoculating a series of nectar microcosms with the same initial density of yeast (approximately 2,500 colony forming units (CFUs) and destructively sampling each day.

To calculate generation time, we used the following equation, assuming the yeast growth was exponential before day two:

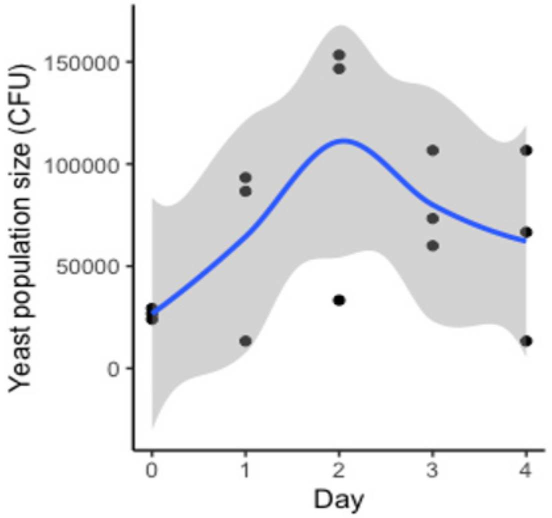


$$\text{Generation time} = \frac{\text{Time (hours)}}{3.3 * \log\left(\dfrac{\text{Population size of yeast on day two}}{\text{Population size of yeast on day one}}\right)}$$

Based on this equation and the data above, we estimate the generation time of *M. reukaufii* yeast (strain MR1) in synthetic nectar is approximately one generation per eight hours. Assuming the generation time stays constant throughout the experiment, we estimate approximately 200 generations over the 30 transfers.

### Supplementary figures

#### Figure supplement 1

*M. reukaufii* yeast and *A. nectaris* bacteria exhibit negative priority effects against each other, as evidenced by growth in microcosm experiments where arrival order is altered. Statistical analysis for significance codes are in ***Supplementary Table 3*** for yeast (**A**) and bacteria (**B**).

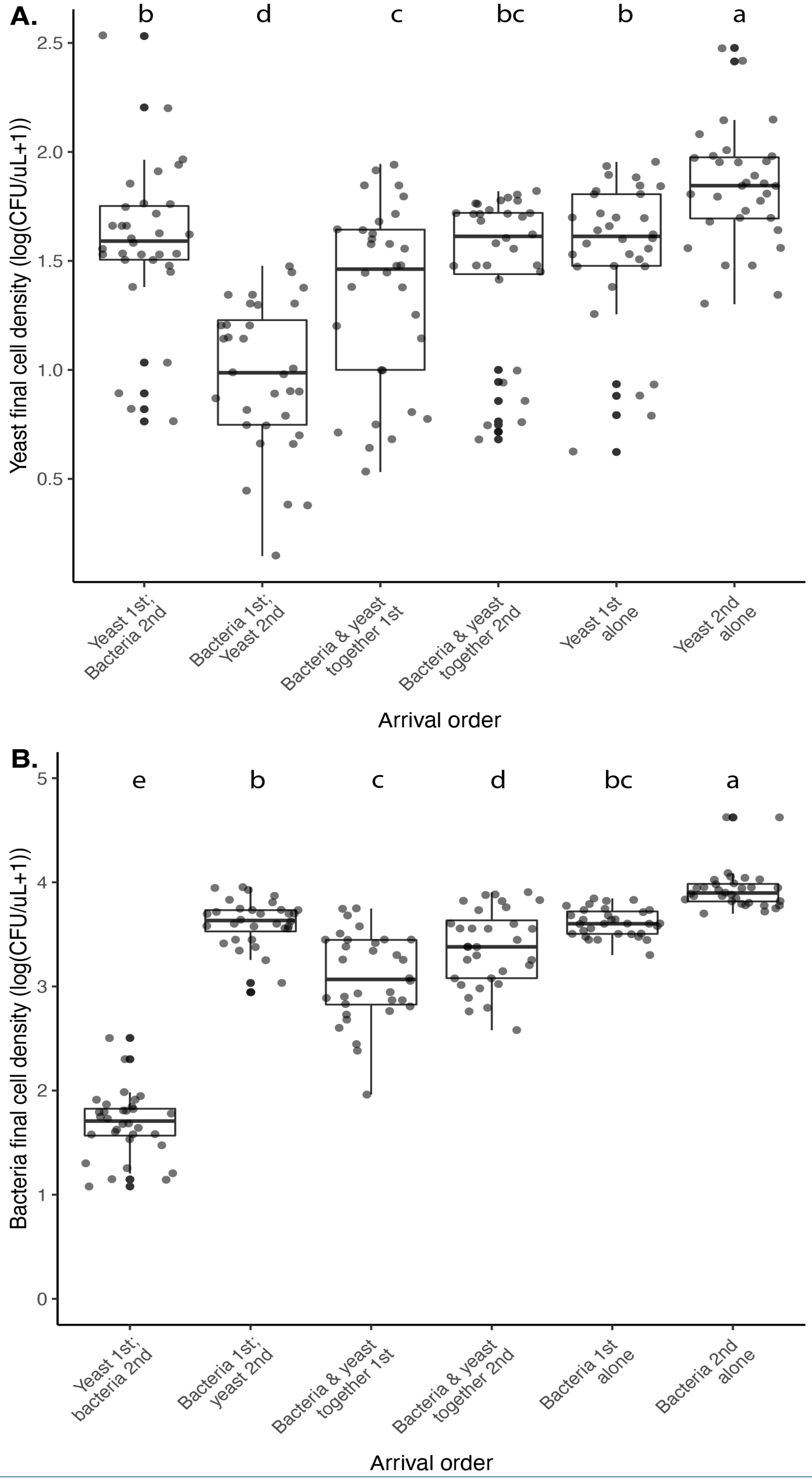

#### Figure supplement 2

Yeast increases nectar pH ($p<0.05$, Spearman rank correlation). The shape of each point represents the various treatments (described in ***Figure 3B***), where yeast first, then bacteria ("YB") are filled circles, bacteria first followed by yeast ("BY") are filled squares, early arriving bacteria and yeast ("YB-") are cross-hatched squares, late arriving bacteria and yeast ("BY-") are cross-hatched circles, late arriving yeast ("-Y) are open squares, early arriving yeast ("Y-") are open diamonds, late arriving bacteria ("-B") are open circles, and early arriving bacteria ("B-") are triangles. Points are slightly jittered on the y axis.

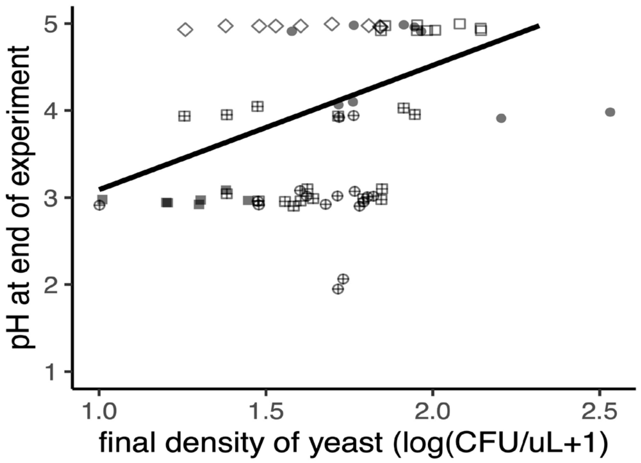


#### Figure supplement 3

We found no effect of nectar type (pH=3, pH=6) on the growth of *M. reukaufii*, when grown in monoculture at a high density (approximately 10,000 cells/μL). *M. reukaufii* growth was calculated by subtracting the final from initial cell density.

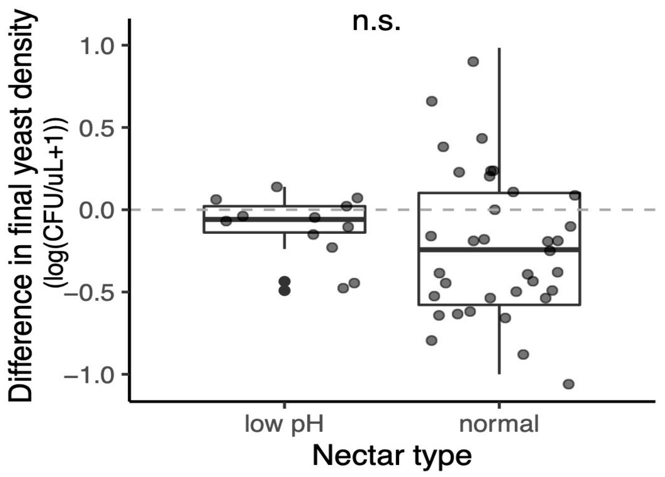

#### Figure supplement 4

We found no effect of nectar type (pH=3, pH=6) on the growth of *A. nectaris* when grown in monoculture at a low density (approximately 10 cells/µL) (**A**) or high density (approximately 10,000 cells/µL) (**B**). *A. nectaris* growth was calculated by subtracting the final from initial cell density.

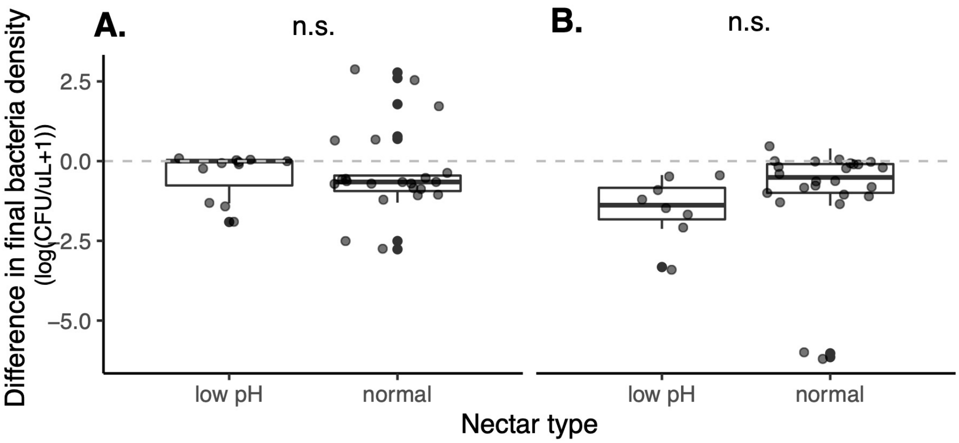


#### Figure supplement 5

We calculated an additional priority effect metric, which corroborated our main result. This metric is calculated by taking the natural logarithm of growth ratio between different initial dominance: PE = log(BY/YB).

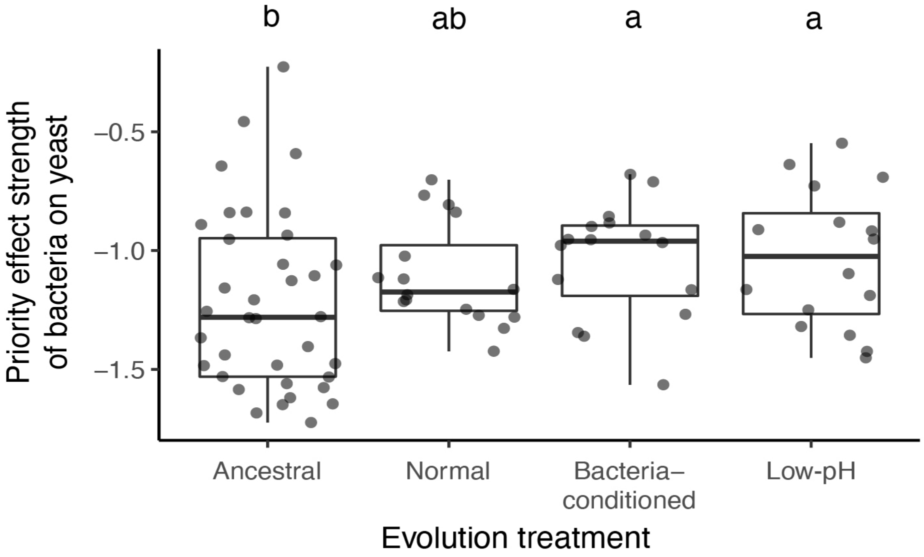

#### Figure supplement 6

(**A**) Yeast evolved in bacteria-conditioned and low-pH nectars were better able to grow than ancestral yeast when bacteria were dominant (BY bacterial priority effects treatment). (**B**) This difference in growth was not due to a difference in intrinsic growth rate (-Y monoculture treatment).

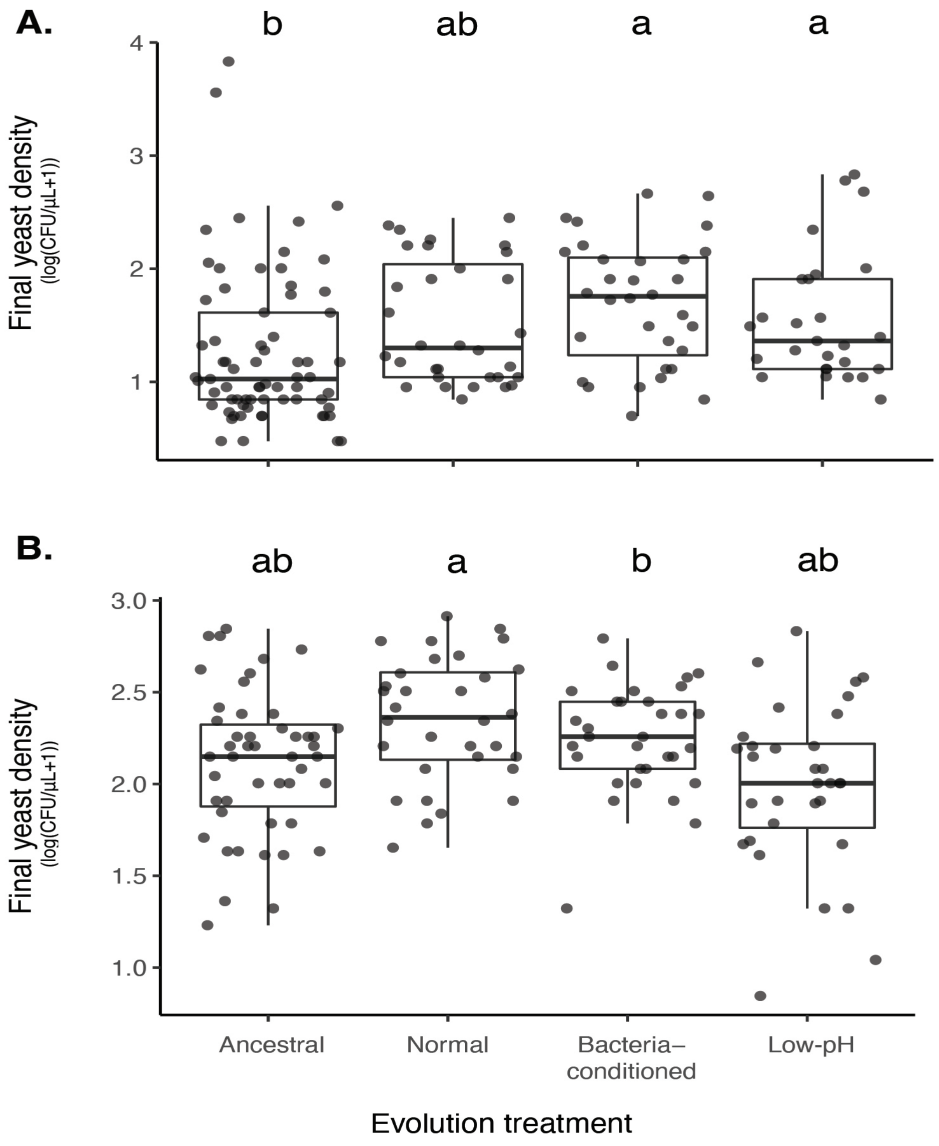

#### Figure supplement 7

Principal components analysis of single nucleotide polymorphisms that differ between evolved and any ancestral strain (2,319 sites), points represent individual end-point clones (individuals) and are colored by evolution treatment (ancestral in black, neutral in grey, evolved in low-pH nectar in orange, and evolved in bacteria-conditioned nectar in dark red). (perMANOVA, p=0.19, n=12).

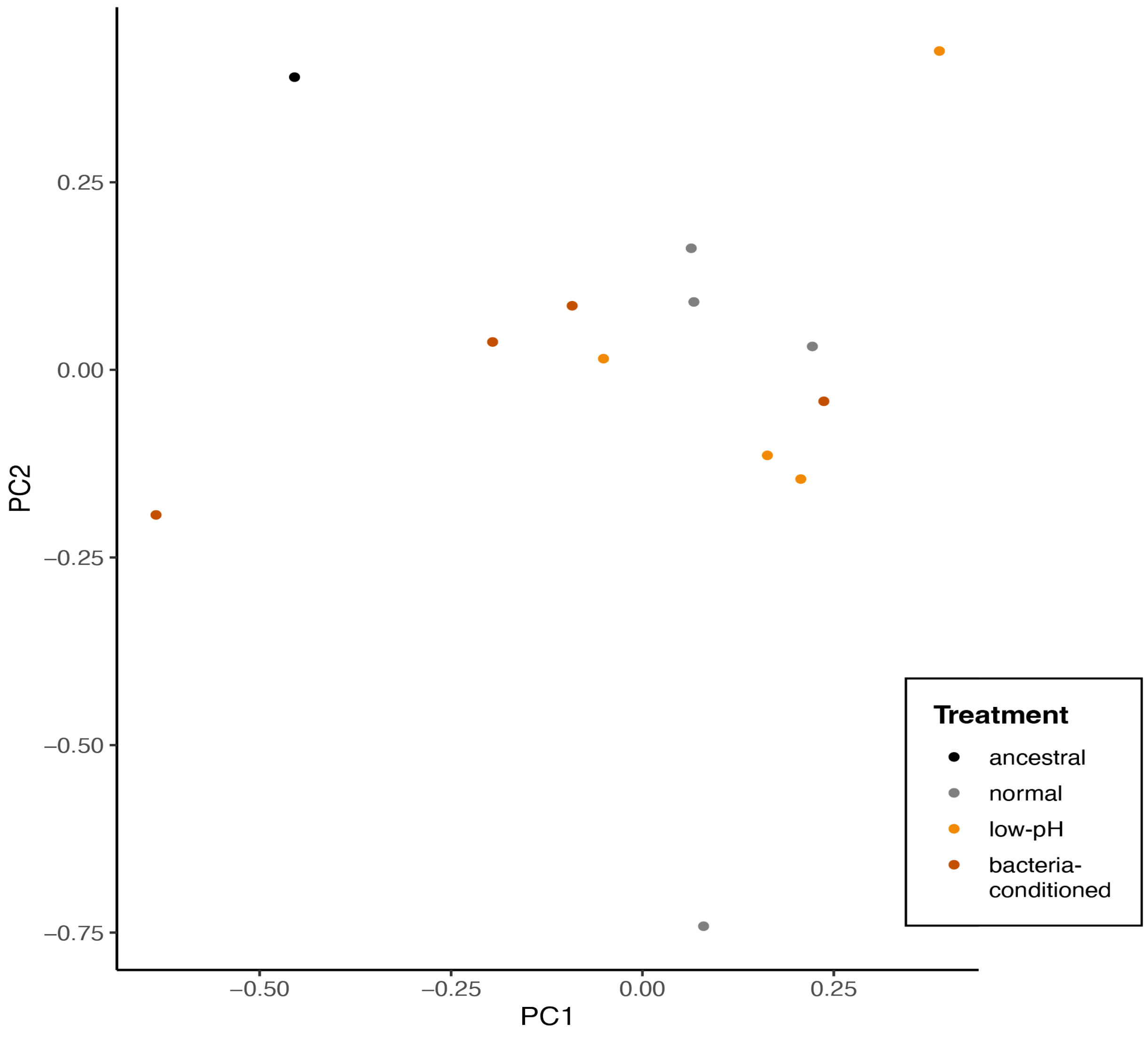

#### Figure supplement 8

Genome-wide differentiation by (**A**) computed p-values for observed patterns of loss of heterozygosity (LOH) and (**B**) Weir-Cockerham estimator of $F_{ST}$.

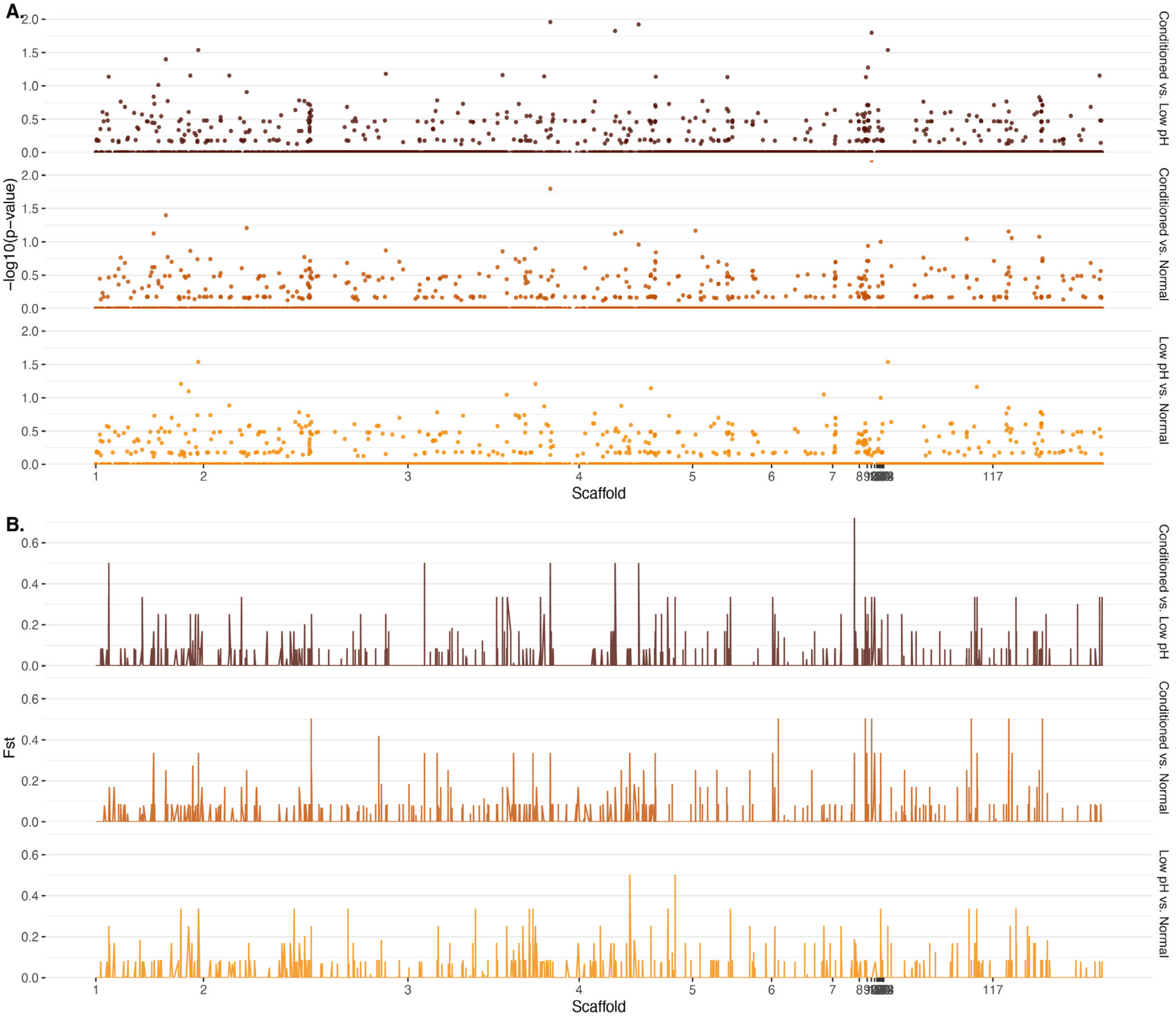

#### Figure supplement 9

No difference in the volume of nectar consumed by nectar treatment in 2016 (**A**) or 2018 (**B**) when nectar sugars were altered with pH.

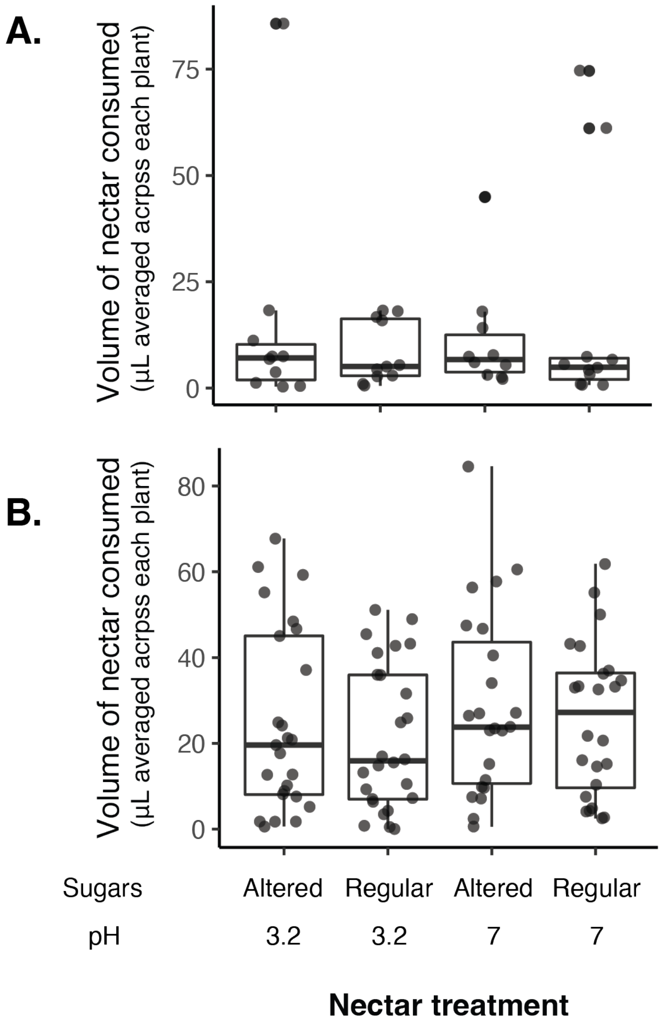

### Supplementary tables

#### Supplementary table 1 - Priority effect experiment treatments

Treatments used in priority effects experiment with ancestral yeast, fully factorial experiment testing the effect of arrival order on priority effects.

| Yeast | | Bacteria | Totals | | |
|---|---|---|---|---|---|
| Strain | Arrival | Arrival | Total rounds | Biological replicates per round | Total biological replicates |
| MR1 | First | Second | 5 | 2 | 10 |
| MR1 | Second | First | 5 | 2 | 10 |
| MR1 | First | First | 5 | 2 | 10 |
| MR1 | Second | Second | 5 | 2 | 10 |
| MR1 | First | None | 5 | 2 | 10 |
| MR1 | Second | None | 5 | 2 | 10 |
| MR1 | None | First | 5 | 2 | 10 |
| MR1 | None | Second | 5 | 2 | 10 |
| MY1082 | First | Second | 1 | 3 | 3 |
| MY1082 | Second | First | 1 | 3 | 3 |
| MY1082 | First | First | 1 | 3 | 3 |
| MY1082 | Second | Second | 1 | 3 | 3 |
| MY1082 | First | None | 1 | 3 | 3 |
| MY1082 | Second | None | 1 | 3 | 3 |
| MY1082 | None | First | 1 | 3 | 3 |
| MY1082 | None | Second | 1 | 3 | 3 |
| MY0202 | First | Second | 1 | 3 | 3 |
| MY0202 | Second | First | 1 | 3 | 3 |
| MY0202 | First | First | 1 | 3 | 3 |
| MY0202 | Second | Second | 1 | 3 | 3 |
| MY0202 | First | None | 1 | 3 | 3 |
| MY0202 | Second | None | 1 | 3 | 3 |
| MY0202 | None | First | 1 | 3 | 3 |
| MY0202 | None | Second | 1 | 3 | 3 |

#### Supplementary table 2 - Evolution treatments

Treatments used in priority effects experiment with experimentally-evolved yeast, fully factorial experiment testing the effect of initial density (10,000 cells/μL ("early") or 10 cells/μL ("late")), evolution treatment (ancestral, evolved in normal nectar, low-pH nectar, or bacteria-conditioned nectar), and evolutionary replicate (independent evolutionary lineages) on priority effects.

| Yeast | | | | Bacteria | | | Totals | |
|---|---|---|---|---|---|---|---|---|
| Species | Evolution | Evolutionary Replicate | Density | Species | Evolution | Density | Biological replicates per round | Total biological replicates |
| *M. reukaufii* | ancestral | none | high | *A. nectaris* | ancestral | low | 9 | 36 |
| *M. reukaufii* | normal | 1 | high | *A. nectaris* | ancestral | low | 2 | 8 |
| *M. reukaufii* | low pH | 1 | high | *A. nectaris* | ancestral | low | 2 | 8 |
| *M. reukaufii* | conditioned | 1 | high | *A. nectaris* | ancestral | low | 2 | 8 |
| *M. reukaufii* | ancestral | none | high | *A. nectaris* | ancestral | low | 9 | 36 |
| *M. reukaufii* | normal | 2 | high | *A. nectaris* | ancestral | low | 2 | 8 |
| *M. reukaufii* | low pH | 2 | high | *A. nectaris* | ancestral | low | 2 | 8 |
| *M. reukaufii* | conditioned | 2 | high | *A. nectaris* | ancestral | low | 2 | 8 |
| *M. reukaufii* | ancestral | none | high | *A. nectaris* | ancestral | low | 9 | 36 |
| *M. reukaufii* | normal | 3 | high | *A. nectaris* | ancestral | low | 2 | 8 |
| *M. reukaufii* | low pH | 3 | high | *A. nectaris* | ancestral | low | 2 | 8 |
| *M. reukaufii* | conditioned | 3 | high | *A. nectaris* | ancestral | low | 2 | 8 |
| *M. reukaufii* | ancestral | none | high | *A. nectaris* | ancestral | low | 9 | 36 |
| *M. reukaufii* | normal | 4 | high | *A. nectaris* | ancestral | low | 2 | 8 |
| *M. reukaufii* | low pH | 4 | high | *A. nectaris* | ancestral | low | 2 | 8 |
| *M. reukaufii* | conditioned | 4 | high | *A. nectaris* | ancestral | low | 2 | 8 |
| *M. reukaufii* | ancestral | none | high | *A. nectaris* | ancestral | low | 9 | 36 |
| *M. reukaufii* | normal | 1 | high | *A. nectaris* | ancestral | low | 2 | 8 |
| *M. reukaufii* | low pH | 1 | high | *A. nectaris* | ancestral | low | 2 | 8 |
| *M. reukaufii* | conditioned | 1 | high | *A. nectaris* | ancestral | low | 2 | 8 |
| *M. reukaufii* | ancestral | none | high | *A. nectaris* | ancestral | low | 9 | 36 |
| *M. reukaufii* | normal | 2 | high | *A. nectaris* | ancestral | low | 2 | 8 |
| *M. reukaufii* | low pH | 2 | high | *A. nectaris* | ancestral | low | 2 | 8 |
| *M. reukaufii* | conditioned | 2 | high | *A. nectaris* | ancestral | low | 2 | 8 |
| *M. reukaufii* | ancestral | none | high | *A. nectaris* | ancestral | low | 9 | 36 |
| *M. reukaufii* | normal | 3 | high | *A. nectaris* | ancestral | low | 2 | 8 |
| *M. reukaufii* | low pH | 3 | high | *A. nectaris* | ancestral | low | 2 | 8 |
| *M. reukaufii* | conditioned | 3 | high | *A. nectaris* | ancestral | low | 2 | 8 |
| *M. reukaufii* | ancestral | none | high | *A. nectaris* | ancestral | low | 9 | 36 |
| *M. reukaufii* | normal | 4 | high | *A. nectaris* | ancestral | low | 2 | 8 |
| *M. reukaufii* | low pH | 4 | high | *A. nectaris* | ancestral | low | 2 | 8 |
| *M. reukaufii* | conditioned | 4 | high | *A. nectaris* | ancestral | low | 2 | 8 |
| *M. reukaufii* | ancestral | none | low | *A. nectaris* | ancestral | high | 9 | 36 |
| *M. reukaufii* | normal | 1 | low | *A. nectaris* | ancestral | high | 2 | 8 |

#### Supplementary table 2 - Evolution treatments (contd.)

| Yeast | | | | Bacteria | | | Totals | |
|---|---|---|---|---|---|---|---|---|
| Species | Evolution | Evolutionary Replicate | Density | Species | Evolution | Density | Biological replicates per round | Total biological replicates |
| *M. reukaufii* | conditioned | 1 | low | *A. nectaris* | ancestral | high | 2 | 8 |
| *M. reukaufii* | ancestral | none | low | *A. nectaris* | ancestral | high | 9 | 36 |
| *M. reukaufii* | normal | 2 | low | *A. nectaris* | ancestral | high | 2 | 8 |
| *M. reukaufii* | low pH | 2 | low | *A. nectaris* | ancestral | high | 2 | 8 |
| *M. reukaufii* | conditioned | 2 | low | *A. nectaris* | ancestral | high | 2 | 8 |
| *M. reukaufii* | ancestral | none | low | *A. nectaris* | ancestral | high | 9 | 36 |
| *M. reukaufii* | normal | 3 | low | *A. nectaris* | ancestral | high | 2 | 8 |
| *M. reukaufii* | low pH | 3 | low | *A. nectaris* | ancestral | high | 2 | 8 |
| *M. reukaufii* | conditioned | 3 | low | *A. nectaris* | ancestral | high | 2 | 8 |
| *M. reukaufii* | ancestral | none | low | *A. nectaris* | ancestral | high | 9 | 36 |
| *M. reukaufii* | normal | 4 | low | *A. nectaris* | ancestral | high | 2 | 8 |
| *M. reukaufii* | low pH | 4 | low | *A. nectaris* | ancestral | high | 2 | 8 |
| *M. reukaufii* | conditioned | 4 | low | *A. nectaris* | ancestral | high | 2 | 8 |
| *M. reukaufii* | ancestral | none | low | *A. nectaris* | ancestral | high | 9 | 36 |
| *M. reukaufii* | normal | 1 | low | *A. nectaris* | ancestral | high | 2 | 8 |
| *M. reukaufii* | low pH | 1 | low | *A. nectaris* | ancestral | high | 2 | 8 |
| *M. reukaufii* | conditioned | 1 | low | *A. nectaris* | ancestral | high | 2 | 8 |
| *M. reukaufii* | ancestral | | low | *A. nectaris* | ancestral | high | 9 | 36 |
| *M. reukaufii* | ancestral | none | low | *A. nectaris* | ancestral | high | 2 | 8 |
| *M. reukaufii* | normal | 2 | low | *A. nectaris* | ancestral | high | 2 | 8 |
| *M. reukaufii* | low pH | 2 | low | *A. nectaris* | ancestral | high | 2 | 8 |
| *M. reukaufii* | conditioned | 2 | low | *A. nectaris* | ancestral | high | 9 | 36 |
| *M. reukaufii* | ancestral | none | low | *A. nectaris* | ancestral | high | 2 | 8 |
| *M. reukaufii* | normal | 3 | low | *A. nectaris* | ancestral | high | 2 | 8 |
| *M. reukaufii* | low pH | 3 | low | *A. nectaris* | ancestral | high | 2 | 8 |
| *M. reukaufii* | conditioned | 3 | low | *A. nectaris* | ancestral | high | 9 | 36 |
| *M. reukaufii* | ancestral | none | low | *A. nectaris* | ancestral | high | 2 | 8 |
| *M. reukaufii* | normal | 4 | low | *A. nectaris* | ancestral | high | 2 | 8 |
| *M. reukaufii* | low pH | 4 | low | *A. nectaris* | ancestral | high | 2 | 8 |
| *M. reukaufii* | conditioned | 4 | low | *A. nectaris* | ancestral | high | 2 | 8 |
| *M. reukaufii* | ancestral | none | high | nectar | nectar | nectar | 6 | 24 |
| *M. reukaufii* | normal | 1 | high | nectar | nectar | nectar | 2 | 8 |
| *M. reukaufii* | low pH | 1 | high | nectar | nectar | nectar | 2 | 8 |
| *M. reukaufii* | conditioned | 1 | high | nectar | nectar | nectar | 2 | 8 |
| *M. reukaufii* | ancestral | none | high | nectar | nectar | nectar | 6 | 24 |
| *M. reukaufii* | normal | 2 | high | nectar | nectar | nectar | 2 | 8 |

#### Supplementary table 2 - Evolution treatments (contd.)

| Yeast | | | | Bacteria | | | Totals | |
|---|---|---|---|---|---|---|---|---|
| Species | Evolution | Evolutionary Replicate | Density | Species | Evolution | Density | Biological replicates per round | Total biological replicates |
| *M. reukaufii* | conditioned | 2 | high | nectar | nectar | nectar | 2 | 8 |
| *M. reukaufii* | ancestral | none | high | nectar | nectar | nectar | 6 | 24 |
| *M. reukaufii* | normal | 3 | high | nectar | nectar | nectar | 2 | 8 |
| *M. reukaufii* | low pH | 3 | high | nectar | nectar | nectar | 2 | 8 |
| *M. reukaufii* | conditioned | 3 | high | nectar | nectar | nectar | 2 | 8 |
| *M. reukaufii* | ancestral | none | high | nectar | nectar | nectar | 6 | 24 |
| *M. reukaufii* | normal | 4 | high | nectar | nectar | nectar | 2 | 8 |
| *M. reukaufii* | low pH | 4 | high | nectar | nectar | nectar | 2 | 8 |
| *M. reukaufii* | conditioned | 4 | high | nectar | nectar | nectar | 2 | 8 |
| *M. reukaufii* | ancestral | none | low | nectar | nectar | nectar | 6 | 24 |
| *M. reukaufii* | normal | 1 | low | nectar | nectar | nectar | 2 | 8 |
| *M. reukaufii* | low pH | 1 | low | nectar | nectar | nectar | 2 | 8 |
| *M. reukaufii* | conditioned | 1 | low | nectar | nectar | nectar | 2 | 8 |
| *M. reukaufii* | ancestral | none | low | nectar | nectar | nectar | 6 | 24 |
| *M. reukaufii* | normal | 2 | low | nectar | nectar | nectar | 2 | 8 |
| *M. reukaufii* | low pH | 2 | low | nectar | nectar | nectar | 2 | 8 |
| *M. reukaufii* | conditioned | 2 | low | nectar | nectar | nectar | 2 | 8 |
| *M. reukaufii* | ancestral | none | low | nectar | nectar | nectar | 6 | 24 |
| *M. reukaufii* | normal | 3 | low | nectar | nectar | nectar | 2 | 8 |
| *M. reukaufii* | low pH | 3 | low | nectar | nectar | nectar | 2 | 8 |
| *M. reukaufii* | conditioned | 3 | low | nectar | nectar | nectar | 2 | 8 |
| *M. reukaufii* | ancestral | none | low | nectar | nectar | nectar | 6 | 24 |
| *M. reukaufii* | normal | 4 | low | nectar | nectar | nectar | 2 | 8 |
| *M. reukaufii* | low pH | 4 | low | nectar | nectar | nectar | 2 | 8 |
| *M. reukaufii* | conditioned | 4 | low | nectar | nectar | nectar | 2 | 8 |
| nectar | nectar | nectar | nectar | *A. nectaris* | ancestral | high | 6 | 24 |
| nectar | nectar | nectar | nectar | *A. nectaris* | ancestral | low | 6 | 24 |

#### Supplementary table 3 - Priority effect experiment results

Results from a linear mixed model testing the effect of arrival order on yeast growth, where BY and YB represents initial arrival by bacteria or yeast, respectively. -Y and Y- represent the comparable growth of yeast at either arrival time (day 0 or day 2). Bold text shows p-values less than or equal to 0.05.

##### (A) Yeast growth:

| Treatment | Estimate | Standard error | Degrees of freedom | t ratio | p value |
|---|---|---|---|---|---|
| YB - BY | 0.6213 | 0.0668 | 183 | 9.296 | **<.0001** |
| YB - (YB-) | 0.2429 | 0.0668 | 183 | 3.635 | **0.0048** |
| YB - (-YB) | 0.1195 | 0.0668 | 183 | 1.788 | 0.4763 |
| YB - (Y-) | 0.0352 | 0.0668 | 183 | 0.527 | 0.995 |
| YB - (-Y) | -0.2456 | 0.0668 | 183 | -3.674 | **0.0042** |
| BY - (YB-) | -0.3783 | 0.0668 | 183 | -5.661 | **<.0001** |
| BY - (-YB) | -0.5018 | 0.0668 | 183 | -7.508 | **<.0001** |
| BY - (Y-) | -0.586 | 0.0668 | 183 | -8.769 | **<.0001** |
| BY - (-Y) | -0.8668 | 0.0668 | 183 | -12.971 | **<.0001** |
| (YB-) - (-YB) | -0.1235 | 0.0668 | 183 | -1.848 | 0.438 |
| (YB-) - (Y-) | -0.2077 | 0.0668 | 183 | -3.108 | **0.0262** |
| (YB-) - (-Y) | -0.4885 | 0.0668 | 183 | -7.31 | **<.0001** |
| (-YB) - (Y-) | -0.0842 | 0.0668 | 183 | -1.26 | 0.806 |
| (-YB) - (-Y) | -0.365 | 0.0668 | 183 | -5.462 | **<.0001** |
| (Y-) - (-Y) | -0.2808 | 0.0668 | 183 | -4.202 | **0.0006** |

##### (B) Bacterial growth:

| Treatment | Estimate | Standard error | Degrees of freedom | t ratio | p value |
|---|---|---|---|---|---|
| YB - BY | -1.92858 | 0.0722 | 183 | -26.717 | <.0001 |
| YB - (YB-) | -1.40927 | 0.0722 | 183 | -19.523 | <.0001 |
| YB - (-YB) | -1.69876 | 0.0722 | 183 | -23.533 | <.0001 |
| YB - (B-) | -1.93511 | 0.0722 | 183 | -26.807 | <.0001 |
| YB - (-B) | -2.25898 | 0.0722 | 183 | -31.294 | <.0001 |
| BY - (YB-) | 0.51931 | 0.0722 | 183 | 7.194 | <.0001 |
| BY - (-YB) | 0.22982 | 0.0722 | 183 | 3.184 | 0.0209 |
| BY - (B-) | -0.00652 | 0.0722 | 183 | -0.09 | 1 |
| BY - (-B) | -0.3304 | 0.0722 | 183 | -4.577 | 0.0001 |
| (YB-) - (-YB) | -0.28949 | 0.0722 | 183 | -4.01 | 0.0012 |
| (YB-) - (B-) | -0.52583 | 0.0722 | 183 | -7.284 | <.0001 |
| (YB-) - (-B) | -0.84971 | 0.0722 | 183 | -11.771 | <.0001 |
| (-YB) - (B-) | -0.23634 | 0.0722 | 183 | -3.274 | 0.0158 |
| (-YB) - (-B) | -0.56022 | 0.0722 | 183 | -7.761 | <.0001 |
| (B-) - (-B) | -0.32388 | 0.0722 | 183 | -4.487 | 0.0002 |

#### Supplementary table 4 - Priority effect experiment results with wild strains

Results from a linear mixed model testing the effect of yeast strain (MR1, MY0182, MY0202) on the strength of priority effects exerted by *A. nectaris* calculated using this metric: PE = log(BY/(-Y)) - log(YB/(Y-)), where BY and YB represents initial dominance by bacteria or yeast, respectively. -Y and Y- represent the comparable growth of yeast at either density, alone and treatment densities were averaged by round of the experiment. Bold text shows p-values less than or equal to 0.05.

| Comparison | Estimate | Standard error | Degrees of freedom | t ratio | p value |
|---|---|---|---|---|---|
| MR1 - MY0182 | -0.453 | 0.0458 | 19 | -9.901 | **<.0001** |
| MR1 - MY0202 | -0.533 | 0.0473 | 19 | -11.285 | **<.0001** |
| MY0182 - MY0202 | -0.08 | 0.0457 | 19 | -1.752 | 0.2126 |

#### Supplementary table 5 - Priority effect experiment results with evolved strains

Results from a linear mixed model testing the effect of evolution treatment (ancestral, evolved in normal nectar, low-pH nectar, or bacteria-conditioned nectar) on the strength of priority effects calculated using one of two metrics:

##### (A) Priority effect metric #1: (Figure 5)

PE1 = log(BY/(-Y)) - log(YB/(Y-)), where BY and YB represents initial dominance by bacteria or yeast, respectively. -Y and Y- represent the comparable growth of yeast at either density, alone and treatment densities were averaged by round of the experiment. Bold text shows p-values less than or equal to 0.05.

| Comparison | Estimate | Standard error | Degrees of freedom | t ratio | p value |
|---|---|---|---|---|---|
| ancestral - normal | -0.11069 | 0.0614 | 77 | -1.803 | 0.28 |
| ancestral - low_pH | -0.18418 | 0.0614 | 77 | -2.999 | **0.0187** |
| ancestral - AN_con | -0.17655 | 0.0614 | 77 | -2.875 | **0.0263** |
| normal - low_pH | -0.07349 | 0.0723 | 77 | -1.017 | 0.74 |
| normal - AN_con | -0.06586 | 0.0723 | 77 | -0.911 | 0.7988 |
| low_pH - AN_con | 0.00762 | 0.0723 | 77 | 0.106 | 0.9996 |

##### (B) Priority effect metric #2: (figure supplement 5)

PE2 = log(BY/YB), where BY and YB represents initial dominance by bacteria or yeast, respectively. Treatment densities were averaged by round of the experiment. Bold text shows p-values less than or equal to 0.05.

| Comparison | Estimate | Standard error | Degrees of freedom | t ratio | p value |
|---|---|---|---|---|---|
| ancestral - normal | -0.11069 | 0.0614 | 77 | -1.803 | 0.28 |
| ancestral - low_pH | -0.18418 | 0.0614 | 77 | -2.999 | **0.0187** |
| ancestral - AN_con | -0.17655 | 0.0614 | 77 | -2.875 | **0.0263** |
| normal - low_pH | -0.07349 | 0.0723 | 77 | -1.017 | 0.74 |
| normal - AN_con | -0.06586 | 0.0723 | 77 | -0.911 | 0.7988 |
| low_pH - AN_con | 0.00762 | 0.0723 | 77 | 0.106 | 0.9996 |

#### Supplementary table 6 - Differences in growth between evolved strains with and without bacteria

Results from a linear mixed model testing the effect of yeast initial density (10,000 colony forming units/μL (“high”) or 10 cells/μL (“low”), yeast monoculture or competition with bacteria) treatment and evolution treatment (ancestral, evolved in normal nectar, low-pH nectar, or bacteria-conditioned nectar) on the difference in final yeast density between treatments with a high density of bacteria and a low density of yeast (BY) and yeast grown in monoculture at a low density (-Y). Growth difference was calculated as (BY) - (-Y). Bold text shows p-values less than or equal to 0.05.

| Comparison | Estimate | Standard error | Degrees of freedom | t ratio | p value |
|---|---|---|---|---|---|
| ancestral - normal | -0.0303 | 0.0969 | 127 | -0.313 | 0.9894 |
| ancestral - low_pH | -0.4636 | 0.1021 | 127 | -4.539 | **0.0001** |
| ancestral - AN_con | -0.3166 | 0.0969 | 127 | -3.267 | **0.0076** |
| normal - low_pH | -0.4333 | 0.1099 | 127 | -3.943 | **0.0008** |
| normal - AN_con | -0.2863 | 0.104 | 127 | -2.754 | **0.0337** |
| low_pH - AN_con | 0.147 | 0.1099 | 127 | 1.338 | 0.5409 |

#### Supplementary table 7 - Difference in final yeast densities between evolved strains with and without bacteria

Results from a linear mixed model testing the effect of yeast initial density (10,000 colony forming units/μL (“early”) or 10 cells/μL (“late”), yeast monoculture or competition with bacteria) treatment and evolution treatment (ancestral, evolved in normal nectar, low-pH nectar, or bacteria-conditioned nectar) on final density of yeast. Bold text shows p-values less than or equal to 0.05.

| Treatment | Comparison | Estimate | Standard error | Degrees of freedom | t ratio | p value |
|---|---|---|---|---|---|---|
| -Y | bacteria conditioned - ancestral | 0.1204 | 0.0967 | 593 | 1.244 | 0.5989 |
| | bacteria conditioned - low pH | 0.2535 | 0.1055 | 593 | 2.402 | 0.0778 |
| | bacteria conditioned - normal | -0.1001 | 0.1055 | 593 | -0.949 | 0.7785 |
| | ancestral - low pH | 0.1331 | 0.0967 | 593 | 1.376 | 0.515 |
| | ancestral - normal | -0.2205 | 0.0967 | 593 | -2.279 | 0.1041 |
| | low pH - normal | -0.3536 | 0.1055 | 593 | -3.351 | **0.0047** |
| BY | bacteria conditioned - ancestral | 0.4175 | 0.0903 | 593 | 4.624 | **<.0001** |
| | bacteria conditioned - low pH | 0.0967 | 0.1105 | 593 | 0.875 | 0.8176 |
| | bacteria conditioned - normal | 0.1862 | 0.1055 | 593 | 1.765 | 0.2916 |
| | ancestral - low pH | -0.3207 | 0.0959 | 593 | -3.344 | **0.0049** |
| | ancestral - normal | -0.2313 | 0.0903 | 593 | -2.562 | 0.0519 |
| | low pH - normal | 0.0895 | 0.1105 | 593 | 0.81 | 0.8499 |
| Y- | bacteria conditioned - ancestral | 0.1658 | 0.0967 | 593 | 1.714 | 0.3173 |
| | bacteria conditioned - low pH | -0.0013 | 0.1055 | 593 | -0.012 | 1 |
| | bacteria conditioned - normal | 0.07 | 0.1064 | 593 | 0.658 | 0.9127 |
| | ancestral - low pH | -0.1671 | 0.0967 | 593 | -1.727 | 0.3104 |
| | ancestral - normal | -0.0958 | 0.0977 | 593 | -0.981 | 0.7605 |
| | low pH - normal | 0.0713 | 0.1064 | 593 | 0.67 | 0.9083 |
| YB | bacteria conditioned - ancestral | 0.5153 | 0.0897 | 593 | 5.747 | **<.0001** |
| | bacteria conditioned - low pH | 0.3152 | 0.1055 | 593 | 2.987 | **0.0155** |
| | bacteria conditioned - normal | 0.1816 | 0.1055 | 593 | 1.721 | 0.3134 |
| | ancestral - low pH | -0.2001 | 0.0897 | 593 | -2.231 | 0.116 |
| | ancestral - normal | -0.3337 | 0.0897 | 593 | -3.721 | **0.0012** |
| | low pH - normal | -0.1336 | 0.1055 | 593 | -1.266 | 0.585 |

#### Supplementary table 8 - Sequencing and mapping QC

Mapping summary of evolved and ancestral genomes. Link to full MultiQC report. There was a total of 675M mapped reads and an average coverage depth of 444X.

| Treatment | Evolutionary replicate | M Reads Mapped | ≥ 30X | Median cov | Mean cov | % Aligned | % Dups | Error rate |
|---|---|---|---|---|---|---|---|---|
| low pH | 1 | 48 | 99.40% | 396.0X | 406.1X | 98.80% | 13.80% | 0.47% |
| bacteria conditioned | 2 | 55.4 | 99.40% | 475.0X | 479.8X | 98.70% | 15.00% | 0.48% |
| bacteria conditioned | 3 | 60.6 | 99.40% | 517.0X | 523.4X | 98.90% | 16.80% | 0.48% |
| bacteria conditioned | 4 | 51.6 | 99.40% | 428.0X | 438.4X | 98.30% | 19.90% | 0.48% |
| Ancestral | NA | 51.9 | 99.30% | 449.0X | 450.7X | 98.70% | 13.90% | 0.48% |
| low pH | 2 | 54.3 | 99.40% | 457.0X | 465.0X | 98.80% | 15.20% | 0.48% |
| low pH | 3 | 51.1 | 99.40% | 425.0X | 433.4X | 98.90% | 14.80% | 0.47% |
| low pH | 4 | 52.4 | 99.40% | 432.0X | 444.2X | 98.80% | 15.70% | 0.48% |
| Normal | 1 | 56.8 | 99.40% | 474.0X | 483.4X | 98.80% | 15.80% | 0.48% |
| Normal | 2 | 51.8 | 99.40% | 430.0X | 439.7X | 98.70% | 14.70% | 0.48% |
| Normal | 3 | 46.1 | 99.30% | 385.0X | 393.4X | 98.80% | 14.00% | 0.48% |
| Normal | 4 | 47.4 | 99.30% | 395.0X | 401.5X | 98.70% | 14.90% | 0.48% |
| bacteria conditioned | 1 | 47.2 | 99.30% | 398.0X | 406.8X | 98.70% | 14.60% | 0.49% |

#### Supplementary table 9 - Nearest annotated gene with treatment-specific divergence

Nearest annotated gene to each loci with treatment specific divergence (loss of heterozygosity permutation test) with p<0.1 and $F_{ST}$ = 0.3-0.5.

| Scaffold | SNP | Treatment comparison | $F_{ST}$ | LOH p-value | InterPro Description | Loci distance to gene (bp) |
|---|---|---|---|---|---|---|
| 2 | 192248 | low pH vs. bacteria-conditioned | 0.5 | NA | NA | 2131 |
| 2 | 721157 | low pH vs. bacteria-conditioned | 0.333333 | NA | Ribosomal protein L7Ae/L30e/S12e/Gadd45 | 241 |
| 2 | 977423 | low pH vs. bacteria-conditioned | 0.25 | 0.0969031 | Alpha/beta hydrolase fold-1 | 4633 |
| 2 | 1335729 | normal vs. low pH | 0.333333 | 0.06193806 | SH3 domain | 4956 |
| 2 | 1458114 | normal vs. low pH | 0.25 | 0.07992008 | Ribosomal protein S9 | 441 |
| 2 | 1484768 | low pH vs. bacteria-conditioned | 0.25 | 0.06993007 | Leucine-rich repeat, cysteine-containing subtype | 1502 |
| 2 | 1610283 | low pH vs. bacteria-conditioned | 0.25 | 0.02897103 | Chorismate mutase, AroQ class, eukaryotic type | 4335 |
| 2 | 1610283 | normal vs. low pH | 0.25 | 0.02897103 | Chorismate mutase, AroQ class, eukaryotic type | 4335 |
| 2 | 1613656 | normal vs. bacteria-conditioned | 0.333333 | NA | Phox homologous domain | 3370 |
| 2 | 1613656 | normal vs. low pH | 0.333333 | NA | Phox homologous domain | 3370 |
| 2 | 2103199 | low pH vs. bacteria-conditioned | 0.25 | 0.06993007 | HAT (Half-A-TPR) repeat | 4950 |
| 2 | 2378124 | normal vs. bacteria-conditioned | 0.25 | 0.06193806 | NOB1 zinc finger-like superfamily | 3480 |
| 3 | 17088 | normal vs. bacteria-conditioned | 0.5 | NA | Tubulin/FtsZ, GTPase domain superfamily | 2796 |
| 3 | 599816 | normal vs. low pH | 0.333333 | NA | HPP | 3373 |
| 3 | 2013434 | normal vs. bacteria-conditioned | 0.333333 | NA | NADP-dependent oxidoreductase domain superfamily | 3350 |
| 3 | 2621204 | normal vs. low pH | 0.333333 | NA | TRP, C-terminal | 4145 |
| 3 | 2957032 | low pH vs. bacteria-conditioned | 0.333333 | NA | Romo1/Mgr2 | 4081 |
| 4 | 16918 | normal vs. low pH | 0.25 | 0.08991009 | SUN domain | 1281 |
| 4 | 26373 | low pH vs. bacteria-conditioned | 0.333333 | NA | NA | 4551 |
| 4 | 124928 | normal vs. bacteria-conditioned | 0.333333 | NA | MIF4G-like, type 3 | 2444 |
| 4 | 376020 | normal vs. low pH | 0.333333 | NA | ATP synthase, F1 complex, epsilon subunit superfamily, mitochondrial | 2960 |
| 4 | 471834 | normal vs. low pH | 0.25 | 0.06193806 | Major facilitator superfamily domain | 2272 |
| 4 | 552417 | low pH vs. bacteria-conditioned | 0.333333 | NA | Hap4 transcription factor, heteromerisation domain | 200 |
| 4 | 1735646 | low pH vs. bacteria-conditioned | 0.5 | 0.01498501 | Something about silencing protein 4 domain | 3921 |

#### Supplementary table 9 - Nearest annotated gene with treatment-specific divergence (contd.)

| Scaffold | SNP | Treatment comparison | $F_{ST}$ | LOH p-value | InterPro Description | Loci distance to gene (bp) |
|---|---|---|---|---|---|---|
| 4 | 1735646 | normal vs. bacteria-conditioned | 0.166667 | 0.07592408 | Something about silencing protein 4 domain | 3921 |
| 4 | 1834583 | normal vs. bacteria-conditioned | 0.25 | 0.07092907 | Multicopper oxidase, type 3 | 4200 |
| 4 | 1968854 | normal vs. bacteria-conditioned | 0.333333 | NA | Aldo/keto reductase | 696 |
| 4 | 1968854 | normal vs. low pH | 0.5 | NA | Aldo/keto reductase | 696 |
| 4 | 2305144 | normal vs. low pH | 0.25 | 0.07192807 | FAS1 domain superfamily | 4783 |
| 5 | 38969 | normal vs. bacteria-conditioned | 0.333333 | NA | Longin-like domain superfamily | 3979 |
| 5 | 48790 | low pH vs. bacteria-conditioned | 0.25 | 0.07292707 | RNA recognition motif domain | 1920 |
| 5 | 236388 | low pH vs. bacteria-conditioned | 0.333333 | NA | Ran binding domain | 1329 |
| 5 | 355522 | low pH vs. bacteria-conditioned | 0.333333 | NA | NA | 3831 |
| 5 | 355522 | normal vs. low pH | 0.5 | NA | NA | 3831 |
| 5 | 678520 | normal vs. bacteria-conditioned | 0.25 | 0.06793207 | Fructose-bisphosphate aldolase, class-II | 3860 |
| 5 | 1184523 | low pH vs. bacteria-conditioned | 0.25 | 0.07392607 | Fatty acid desaturase domain | 4206 |
| 5 | 1232754 | low pH vs. bacteria-conditioned | 0.333333 | NA | Vicinal oxygen chelate (VOC) domain | 1376 |
| 5 | 1232754 | normal vs. low pH | 0.333333 | NA | Vicinal oxygen chelate (VOC) domain | 1376 |
| 6 | 639407 | low pH vs. bacteria-conditioned | 0.333333 | NA | Calcineurin-like phosphoesterase domain, ApaH type | 1782 |
| 6 | 639407 | normal vs. bacteria-conditioned | 0.333333 | NA | Calcineurin-like phosphoesterase domain, ApaH type | 1782 |
| 6 | 728208 | normal vs. bacteria-conditioned | 0.5 | NA | Carbohydrate kinase PfkB | 4911 |
| 6 | 728220 | normal vs. bacteria-conditioned | 0.333333 | NA | Carbohydrate kinase PfkB | 4923 |
| 7 | 206974 | normal vs. low pH | 0.25 | 0.08891109 | LicD family | 2246 |
| 8 | 15759 | low pH vs. bacteria-conditioned | 0.722222 | NA | Cytochrome c-like domain | 459 |
| 8 | 15759 | normal vs. bacteria-conditioned | 0.333333 | NA | Cytochrome c-like domain | 459 |
| 9 | 3961 | low pH vs. bacteria-conditioned | 0.333333 | NA | Chaperone DnaK | 2032 |
| 9 | 3961 | normal vs. bacteria-conditioned | 0.5 | NA | Chaperone DnaK | 2032 |
| 9 | 27928 | normal vs. bacteria-conditioned | 0.333333 | NA | Rho protein GDP-dissociation inhibitor | 1130 |
| 10 | 36322 | low pH vs. bacteria-conditioned | 0.333333 | 0.01598402 | DASH complex subunit Ask1 | 2195 |
| 10 | 36322 | normal vs. bacteria-conditioned | 0.5 | 0.00599401 | DASH complex subunit Ask1 | 2195 |

#### Supplementary table 9 - Nearest annotated gene with treatment-specific divergence (contd.)

| Scaffold | SNP | Treatment comparison | $F_{ST}$ | LOH p-value | InterPro Description | Loci distance to gene (bp) |
|---|---|---|---|---|---|---|
| 50 | 14804 | normal vs. bacteria-conditioned | 0.333333 | NA | Exocyst complex component EXOC3/Sec6, C-terminal domain | 4348 |
| 50 | 14804 | normal vs. low pH | 0.333333 | NA | Exocyst complex component EXOC3/Sec6, C-terminal domain | 4348 |
| 50 | 16264 | normal vs. bacteria-conditioned | 0.25 | 0.0999001 | NA | 4606 |
| 50 | 16264 | normal vs. low pH | 0.25 | 0.0999001 | NA | 4606 |
| 117 | 65833 | low pH vs. bacteria-conditioned | 0.25 | 0.02897103 | NA | 4754 |
| 117 | 65833 | normal vs. low pH | 0.25 | 0.02897103 | NA | 4754 |
| 117 | 1316972 | normal vs. bacteria-conditioned | 0.25 | 0.08991009 | Rho GTPase-activating protein domain | 4144 |
| 117 | 1443605 | low pH vs. bacteria-conditioned | 0.333333 | NA | 2-isopropylmalate synthase | 3828 |
| 117 | 1476712 | normal vs. low pH | 0.333333 | 0.06893107 | TCP-1-like chaperonin intermedi-ate domain superfamily | 3740 |
| 117 | 1481377 | low pH vs. bacteria-conditioned | 0.333333 | NA | Protein kinase domain | 4037 |
| 117 | 1481377 | normal vs. bacteria-conditioned | 0.333333 | NA | Protein kinase domain | 4037 |
| 117 | 1981533 | normal vs. bacteria-conditioned | 0.25 | 0.06993007 | Thioredoxin domain | 3792 |
| 117 | 1984793 | normal vs. bacteria-conditioned | 0.5 | NA | Eukaryotic translation initiation factor 3 subunit J | 2334 |
| 117 | 2030654 | normal vs. bacteria-conditioned | 0.25 | 0.08791209 | Leucine-rich repeat domain superfamily | 4296 |
| 117 | 2034922 | normal vs. bacteria-conditioned | 0.333333 | NA | Mss4 | 4890 |
| 117 | 2097637 | low pH vs. bacteria-conditioned | 0.333333 | NA | Cysteine-rich transmembrane CYSTM domain | 3186 |
| 117 | 2097637 | normal vs. low pH | 0.333333 | NA | Cysteine-rich transmembrane CYSTM domain | 3186 |
| 117 | 2517317 | normal vs. bacteria-conditioned | 0.5 | NA | Protein kinase domain | 4074 |
| 117 | 3462569 | low pH vs. bacteria-conditioned | 0.333333 | NA | Histidine phosphatase superfami-ly, clade-1 | 4722 |

#### Supplementary table 10 - Nearest annotated gene with punitive de novo singleton mutation

Nearest annotated gene to each putative *de novo* singleton mutation.

| Scaffold | SNP | Treatment | Gene ID | Signature Description | InterPro Description | Loci distance to gene (bp) |
|---|---|---|---|---|---|---|
| 2 | 142172 | bacteria conditioned | 122804 | SHNi-TPR | Tetratricopeptide, SHNi-TPR domain | 4330 |
| 2 | 152222 | low pH | 122810 | Zn(2)-C6 fungal-type DNA-binding domain profile. | Zn(2)-C6 fungal-type DNA-binding domain | 4580 |
| 2 | 581610 | low pH | 123038 | - | Mitochondrial import inner membrane translocase subunit Tim21 | 3315 |
| 2 | 874406 | normal | 158686 | SET domain profile. | SET domain | 3525 |
| 2 | 983739 | bacteria conditioned | 143523 | Beta-lactamase | Beta-lactamase-related | 3091 |
| 2 | 1193730 | bacteria conditioned | 123348 | SEC7 domain profile. | Sec7 domain | 3703 |
| 2 | 1214463 | bacteria conditioned | 104900 | RNA-binding domain, RBD | RNA-binding domain superfamily | 4296 |
| 2 | 1387782 | low pH | 158874 | Taurine catabolism dioxygen-ase TauD, TfdA family | TauD/TfdA-like domain | 4020 |
| 2 | 1801552 | ancestral | 143718 | - | Exocyst complex com-ponent EXOC6/Sec15, C-terminal, domain 2 | 3974 |
| 2 | 2270572 | low pH | 165272 | ABC transporter integral membrane type-1 fused domain profile. | ABC transporter type 1, transmembrane domain | 4315 |
| 2 | 2613284 | bacteria conditioned | 140298 | HCP-like | NA | 3499 |
| 2 | 2654116 | bacteria conditioned | 124080 | CYTOCHROME C OXIDASE SUBUNIT 6A, MITOCHON-DRIAL | NA | 2528 |
| 2 | 2654116 | bacteria conditioned | 124080 | CYTOCHROME C OXIDASE POLYPEPTIDE VIA | Cytochrome c oxidase, subunit VIa | 2528 |
| 2 | 2945111 | bacteria conditioned | 159455 | ER MEMBRANE PROTEIN COMPLEX SUBUNIT 6 | NA | 2215 |
| 2 | 2945111 | bacteria conditioned | 159455 | UNCHARACTERIZED | ER membrane protein complex subunit 6 | 2215 |
| 3 | 15656 | bacteria conditioned | 125316 | - | Tubulin/FtsZ, GTPase domain superfamily | 4228 |
| 3 | 387084 | bacteria conditioned | 125523 | Ferredoxin reductase-type FAD binding domain profile. | FAD-binding domain, ferredoxin reduc-tase-type | 4644 |
| 3 | 792251 | low pH | 140953 | Classic Zinc Finger | NA | 4652 |
| 3 | 1477772 | low pH | 141117 | Protein kinase domain profile. | Protein kinase domain | 3437 |
| 3 | 1535525 | normal | 156507 | Nnf1 | Nuclear MIS12/MIND complex subunit PMF1/Nnf1 | 3517 |
| 3 | 2075720 | low pH | 149825 | ALG6, ALG8 glycosyltrans-ferase family | Glycosyl transferase, ALG6/ALG8 | 3878 |

#### Supplementary table 10 - Nearest annotated gene with punitive de novo singleton mutation (contd.)

| Scaffold | SNP | Treatment | Gene ID | Signature Description | InterPro Description | Loci distance to gene (bp) |
|---|---|---|---|---|---|---|
| 3 | 2144184 | normal | 156735 | Cytidine and deoxycytidylate deaminases domain profile. | Cytidine and deoxy-cytidylate deaminase domain | 3764 |
| 3 | 2353536 | low pH | 117782 | ARM repeat | Armadillo-type fold | 2219 |
| 3 | 2436246 | normal | 117810 | NNP-1 PROTEIN NOVEL NUCLEAR PROTEIN 1 NOP52 | Nucleolar, Nop52 | 4021 |
| 3 | 2671944 | bacteria conditioned | 117893 | Aldehyde Dehydrogenase; Chain A | Aldehyde dehydroge-nase, N-terminal | 4924 |
| 3 | 2671944 | bacteria conditioned | 117893 | Aldehyde Dehydrogenase; Chain A | Aldehyde dehydroge-nase, C-terminal | 4924 |
| 3 | 2753650 | low pH | 156963 | Fe(2+) 2-oxoglutarate dioxy-genase domain profile. | Oxoglutarate/iron-de-pendent dioxygenase | 4708 |
| 3 | 2909079 | bacteria conditioned | 117984 | Solute carrier (Solcar) repeat profile. | Mitochondrial sub-strate/solute carrier | 4378 |
| 3 | 3082088 | normal | 118051 | TTL domain profile. | Tubulin-tyrosine ligase/ Tubulin polyglutam-ylase | 4200 |
| 4 | 79542 | normal | 118841 | t-SNARE coiled-coil homolo-gy domain profile. | Target SNARE coiled-coil homology domain | 2595 |
| 4 | 262947 | normal | 141816 | Regulator of G-protein sig-naling, RGS | RGS domain super-family | 4830 |
| 4 | 286255 | normal | 112888 | Prokaryotic membrane lipoprotein lipid attachment site profile. | NA | 1486 |
| 4 | 414723 | bacteria conditioned | 138051 | Eukaryotic RNA Recognition Motif (RRM) profile. | RNA recognition motif domain | 2087 |
| 4 | 756357 | low pH | 138127 | - | Fe-S cluster assembly domain superfamily | 2038 |
| 4 | 1536342 | low pH | 160157 | Acetyl-coenzyme A (CoA) carboxyltransferase N-termi-nal domain profile. | Acetyl-coenzyme A carboxyltransferase, N-terminal | 3642 |
| 4 | 1692875 | low pH | 113406 | PRA1 family protein | Prenylated rab accep-tor PRA1 | 325 |
| 4 | 1780889 | bacteria conditioned | 160246 | Forkhead-associated (FHA) domain profile. | Forkhead-associated (FHA) domain | 3677 |
| 4 | 1966106 | normal | 142199 | AKR | Aldo/keto reductase | 3444 |
| 4 | 2163139 | low pH | 154278 | Pyoverdine/dityrosine bio-synthesis protein | Pyoverdine/dityrosine biosynthesis protein | 636 |
| 4 | 2190640 | bacteria conditioned | 119880 | Ubiquitin specific protease (USP) domain profile. | Ubiquitin specific protease domain | 3341 |
| 5 | 38989 | normal | 124511 | SNARE-like | Longin-like domain superfamily | 3959 |

#### Supplementary table 10 - Nearest annotated gene with punitive de novo singleton mutation (contd.)

| Scaffold | SNP | Treatment | Gene ID | Signature Description | InterPro Description | Loci distance to gene (bp) |
|---|---|---|---|---|---|---|
| 5 | 39790 | normal | 124511 | SNARE-like | Longin-like domain superfamily | 3158 |
| 5 | 252114 | low pH | 154487 | Candida agglutinin-like (ALS) | Agglutinin-like protein repeat | 3285 |
| 5 | 252114 | low pH | 154487 | Candida agglutinin-like (ALS) | Agglutinin-like protein repeat | 3285 |
| 5 | 252114 | low pH | 154487 | Candida agglutinin-like (ALS) | Agglutinin-like protein repeat | 3285 |
| 5 | 549641 | bacteria conditioned | 116652 | Translation proteins SH3-like domain | Translation protein SH3-like domain superfamily | 2612 |
| 6 | 319227 | normal | 161928 | MFS general substrate transporter like domains | NA | 3992 |
| 6 | 319531 | ancestral | 161928 | MFS general substrate transporter like domains | NA | 3878 |
| 6 | 427980 | bacteria conditioned | 118319 | TLDc domain profile. | NA | 1247 |
| 6 | 528907 | normal | 141612 | PLA2c domain profile. | Lysophospholipase, catalytic domain | 3786 |
| 6 | 686853 | low pH | 141643 | TLC domain profile. | TRAM/LAG1/CLN8 homology domain | 3970 |
| 6 | 699796 | normal | 118472 | Major facilitator superfamily (MFS) profile. | Major facilitator superfamily domain | 1840 |
| 6 | 980857 | bacteria conditioned | 118638 | alpha/beta-Hydrolases | Alpha/Beta hydrolase fold | 862 |
| 6 | 992775 | normal | 118648 | SIGMA 1-TYPE OPIOID RECEPTOR-RELATED | ERG2/sigma1 receptor-like | 4387 |
| 6 | 1127358 | low pH | 118711 | Mu homology domain (MHD) profile. | Mu homology domain | 4349 |
| 8 | 2881 | normal | 157102 | - | NA | 3261 |
| 8 | 64919 | low pH | 103781 | TB2/DP1, HVA22 family | TB2/DP1/HVA22-related protein | 3451 |
| 9 | 26756 | bacteria conditioned | 125192 | RHO protein GDP dissociation inhibitor | Rho protein GDP-dissociation inhibitor | 2302 |
| 54 | 7419 | normal | 120011 | Serine proteases, subtilase domain profile. | NA | 4015 |
| 54 | 11135 | low pH | 120011 | Serine proteases, subtilase domain profile. | NA | 299 |
| 54 | 11152 | bacteria conditioned | 120011 | Serine proteases, subtilase domain profile. | NA | 282 |
| 54 | 11862 | low pH | 120011 | Serine proteases, subtilase domain profile. | NA | 737 |
| 54 | 12493 | normal | 120011 | Serine proteases, subtilase domain profile. | NA | 1368 |
| 117 | 20485 | bacteria conditioned | 160460 | ARM repeat | Armadillo-type fold | 974 |
| 117 | 20595 | bacteria conditioned | 160460 | ARM repeat | Armadillo-type fold | 864 |

#### Supplementary table 10 - Nearest annotated gene with punitive de novo singleton mutation (contd.)

| Scaffold | SNP | Treatment | Gene ID | Signature Description | InterPro Description | Loci distance to gene (bp) |
|---|---|---|---|---|---|---|
| 117 | 341822 | bacteria conditioned | 120229 | Transcription mediator complex subunit Med12 | Mediator complex, subunit Med12 | 3644 |
| 117 | 592420 | bacteria conditioned | 120359 | Zinc finger C2H2 type domain profile. | Zinc finger C2H2-type | 4695 |
| 117 | 668209 | bacteria conditioned | 120406 | Glycosyl hydrolases family 18 | Glycoside hydrolase family 18, catalytic domain | 3687 |
| 117 | 925961 | bacteria conditioned | 163279 | Nucleotide-diphospho-sugar transferases | Nucleotide-diphospho-sugar transferases | 4190 |
| 117 | 1330373 | low pH | 163422 | Protein kinase domain profile. | Protein kinase domain | 4774 |
| 117 | 1330453 | low pH | 163422 | Protein kinase domain profile. | Protein kinase domain | 4854 |
| 117 | 1331269 | bacteria conditioned | 120799 | Allantoicase repeat | Allantoicase domain | 3260 |
| 117 | 1392788 | normal | 142560 | Zn(2)-C6 fungal-type DNA-binding domain profile. | Zn(2)-C6 fungal-type DNA-binding domain | 4298 |
| 117 | 1550134 | normal | 120918 | Transmembrane amino acid transporter protein | Amino acid transporter, transmembrane domain | 3766 |
| 117 | 1578280 | low pH | 120929 | - | NA | 1510 |
| 117 | 1685479 | bacteria conditioned | 163532 | USP_Like | NA | 2067 |
| 117 | 1685494 | bacteria conditioned | 163532 | USP_Like | NA | 2052 |
| 117 | 1685560 | normal | 163532 | USP_Like | NA | 1986 |
| 117 | 1698523 | low pH | 163538 | UNCHARACTERIZED | Pre-rRNA-processing protein Esf1 | 4656 |
| 117 | 1850710 | ancestral | 121089 | - | NA | 1113 |
| 117 | 1914706 | bacteria conditioned | 121132 | SAM-dependent MTase RsmB/NOP-type domain profile. | SAM-dependent methyltransferase RsmB/NOP2-type | 4575 |
| 117 | 2011919 | normal | 114390 | - | NA | 3312 |
| 117 | 2322449 | low pH | 121409 | Cyclin_C_H_G | NA | 2345 |
| 117 | 2375781 | low pH | 161383 | - | HAD superfamily | 3555 |
| 117 | 2393784 | bacteria conditioned | 161388 | WD40 repeat-like | WD40-repeat-containing domain superfamily | 1282 |
| 117 | 2395046 | normal | 121448 | Histone-fold | Histone-fold | 4332 |
| 117 | 2499913 | low pH | 163813 | Universal stress protein signature | Universal stress protein A family | 2742 |
| 117 | 2499913 | low pH | 163813 | Universal stress protein signature | Universal stress protein A family | 2742 |
| 117 | 2499913 | low pH | 163813 | Universal stress protein signature | Universal stress protein A family | 2742 |
| 117 | 2681072 | bacteria conditioned | 161494 | Protein kinase domain profile. | Protein kinase domain | 1015 |
| 117 | 2925974 | low pH | 163965 | Heavy-metal-associated domain profile. | Heavy metal-associated domain, HMA | 2173 |

#### Supplementary table 10 - Nearest annotated gene with punitive de novo singleton mutation (contd.)

| Scaffold | SNP | Treatment | Gene ID | Signature Description | InterPro Description | Loci distance to gene (bp) |
|---|---|---|---|---|---|---|
| 117 | 3039368 | normal | 147163 | Vps54-like protein | Vacuolar protein sorting-associated protein 54, C-terminal | 4292 |
| 117 | 3198640 | low pH | 103079 | - | WD40/YVTN repeat-like-containing domain superfamily | 2 |
